# Embedding kernels for sequence-function relationships ^*^

**DOI:** 10.64898/2026.11.15.766687

**Authors:** Waverly Carabba, Carlos Martí-Gómez, David M. McCandlish, Samantha Petti

**Author notes:** Co-first authors (Waverly Carabba, Carlos Martí-Gómez). Submitted to the editors November 15, 2026.

## Abstract

A central challenge in biology is understanding how variation in biological sequences (DNA, RNA, and protein) impacts function. Recent advances in high-throughput techniques have enabled the measurement of such sequence–function relationships on an unprecedented scale. However, analyzing these complex datasets requires models that are both flexible and mathematically tractable. We consider Gaussian process regression, a Bayesian method that specifies a Gaussian prior distribution over sequence–function relationships. The prior is defined by a kernel function that encodes the expected covariance between pairs of sequences. We study a family of embedding kernels in which the covariance depends on the distance between sequences in a learned embedding space, making these kernels both expressive and interpretable. We show that linear embedding kernels generalize previously studied classes of kernels and can more flexibly express variability in the expected similarity between mutational effects across genetic backgrounds, allowing the resulting priors to encode more complex biological mechanisms such as RNA base-pairing. We implement these methods using a GPU-accelerated Gaussian process framework, allowing us to learn linear embeddings and make predictions with datasets containing hundreds of thousands of measurements. We find that our embedding kernels match or improve the predictive performance of other kernels across various high-throughput sequence–function datasets, including protein datasets ranging from short motifs to full-length proteins. Furthermore, the learned embedding spaces provide insight into biological mechanisms underlying sequence–function relationships.

## 1. Introduction

Understanding how genetic sequences (DNA, RNA, and proteins) determine biological function remains a central challenge in biology. Characterizing this sequence– function map would improve our understanding of how mutations alter biological function [18, 38, 28, 64, 29], helping to uncover relevant biological mechanisms for a range of applications [36, 40], including understanding genetic and infectious diseases [12, 47], crop improvement [84, 9], and protein engineering for therapeutics and biomanufacturing [11, 82, 51].

Learning this map is difficult because sequence space, a discrete space whose size depends on the alphabet size and sequence length, grows astronomically large even for moderate-length sequences. This high dimensionality makes it impractical to measure every possible sequence in sequence space for most datasets [65, 23]. Even when experiments exhaustively measure the phenotypes of all possible variants across a small number of sequence positions [80, 57, 42, 32, 14, 85, 34, 25], the resulting sequence–function relationship can still be complex due to a phenomenon known as epistasis, in which the effect of a mutation varies depends on the genetic background in which it is introduced. These epistatic interactions include both global or non-specific epistasis, when the effect of a mutation depends systematically on the phenotype of the genetic background in which it is introduced [50, 13, 39], as well as specific epistasis, when the effect of a mutation depends not just on the phenotype of the background but also on the identities of the other mutations present in that background [70, 78, 10]. Global epistasis often arises from biophysical nonlinearities (e.g. the nonlinear relationship between folding energy and the probability a protein is folded [5, 70, 50]) even when mutations have additive effects on the underlying biophysical trait. Specific epistasis often reflects structural or functional couplings between particular residues or positions [62, 66]. As a result of this complexity, it is not obvious a priori which features of a sequence are most useful for predicting its function [70, 55, 64, 13].

A variety of approaches have been used to model sequence–function relationships, including linear regression [26, 57, 54], Gaussian process regression [63, 83, 33], and neural networks trained directly on sequence–function data [21, 6, 43, 74, 19, 67]. Linear models are highly interpretable, but capturing complex patterns of epistasis can require large numbers of interaction terms. Neural networks, on the other hand, can capture complex, higher-order structure but are difficult to interpret. Gaussian processes (GPs) offer a natural middle ground between these two extremes. Rather than finding a particular function that best describes the data, a GP uses a Bayesian framework to provide a distribution of plausible functions that are compatible with the data. In particular, these models work by first defining a prior distribution describing our belief about the plausibility of each possible genotype-phenotype map before seeing any data. Then, conditioning on the observed data yields a posterior distribution, which reflects our updated knowledge about the particular sequence–function relationship we are studying [60]. A key modeling choice is specifying the prior distribution, which is defined through a covariance or kernel function that encodes our expectations about the phenotypic similarity of two arbitrary sequences. Consequently, the choice of kernel and how well it can capture the biological mechanisms driving phenotypic variation will govern both the predictive performance of the model and the interpretability of its underlying parameters.

GPs have been used in different ways to model sequence–function relationships. One approach is to embed sequences in a continuous space and apply classical kernels for GP regression directly on this space to define a prior [63, 83, 4, 24, 30, 33]. Alternatively, we can construct specific priors for sequence–function relationships whose parameters control relevant properties, such as the average magnitude of local epistatic coefficients [87, 8], the variance explained by genetic interactions of different orders [88, 44] or different sets of sites [45], and the effects that mutations at different positions have on the predictability of one another through General Product kernels (GPKs) [86]. In particular, GPKs can be computed efficiently and implemented within GPU-accelerated frameworks [20, 76, 7], enabling scalable inference for sequences of arbitrary length and estimation of kernel parameters from the data through evidence maximization. The resulting parameter estimates can then be used both to interpret and to guide the exploration of the sequence–function relationship. However, the expressivity of these priors is limited because the expected covariance between two sequences is determined by independent contributions from the positions at which they differ, thus restricting the range of molecular mechanisms that they can represent.

In this work, we develop a principled mathematical framework for understanding and applying classical embedding kernels designed for continuous spaces to functions defined over discrete sequence spaces. In particular, we focus on the family of linear embedding kernels, also known as Factor Analysis (FA) kernels [75, 60], in which the prior covariance between two sequences decays exponentially with the squared Euclidean distance in a space defined by linear functions of their one-hot representations. We explain the mathematical relationship between FA kernels and GPKs and describe how the FA can encode a richer structure of dependencies in mutational effects across backgrounds and positions. We implement the FA kernel within a GPU-accelerated Gaussian process framework [20, 76, 7] to estimate these linear embeddings from data via evidence maximization. Our formulation of FA kernels complements an alternative latent-space approach, LANTERN [73], by providing a direct mathematical relationship to GPKs and removing the requirement of a prior on the latent space. For a detailed comparison of these two methods, see Supplementary Materials Section SM1.7.

The rest of the paper is structured as follows. In Section 2, we provide a short introduction to Gaussian process models. In Section 3, we define the FA kernel and show that it generalizes a class of GPKs, which can be represented by an FA kernel where each embedding dimension contains information about only a single position in the sequence. By relaxing this restriction, FA kernels give us more flexibility in controlling both the expected similarity between the effects of mutations at different positions within the same background and the expected similarity of a fixed mutation’s effect across backgrounds that differ at a specific set of positions. This interpretable representation enables us to identify aspects of sequence–function relationships that cannot be directly represented by GPKs, including biologically relevant phenomena such as global epistasis and RNA base-pairing configurations. In Section 5, we describe the implementation of these kernels in a GPU-accelerated framework for GPs [20, 76, 7], which allows estimation of the kernel’s hyperparameters via evidence maximization and computation of the posterior distribution for phenotypic prediction of unknown sequences. We identify two sources of nonidentifiability in the parametrization of these kernels: the translational and rotational invariance of distances in the embedding space, and the gauge freedoms associated with one-hot representations [59, 58]. We address both by developing a gauge-fixed optimization procedure together with a systematic and unique representation of the learned embeddings for interpretation. Finally, in Sections 6.1-6.4, we apply our FA models to analyze four sequence–function datasets: the binding affinity of GB1 variants [80], the splicing efficiency of a self-splicing group I intron [68], the AAV2 capsid assembly [6], and the fluorescence of GFP [65, 23] and explore their predictive capabilities and the biological interpretations of their learned representations. We find that the FA kernel improves upon or provides comparable predictive performance to GPKs but learns features that can be interpreted to make hypotheses about the driving biological mechanisms in these datasets. Furthermore, we find that learning these embeddings directly, rather than precomputing them from patterns found from natural sequence data or protein language models [41], substantially improves predictive performance (see Supplementary Materials Section SM1.8).

## 2. Gaussian process models

Let *s* be a sequence of length *ℓ* with characters drawn from an alphabet *A* = {*c*_1_, …, *c*_*α*_} composed of *α* different characters (*α* = 4 for DNA sequences and *α* = 20 for protein sequences). A sequence–function map can be defined as a function that assigns a continuous phenotypic value to every possible sequence in the space *s* ∈ *A*^*ℓ*^ given by *f* : *A*^*ℓ*^ → ℝ. Since there are *α*^*ℓ*^ possible sequences, *f* can also be viewed as an 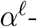 dimensional vector 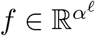. In this work, our goal is to infer sequence–function relationships from experimental measurements of a subset of sequences using a class of Bayesian methods called Gaussian process (GP) regression [60].

Before any data is observed, we encode our assumptions about the sequence–function relationship in a Gaussian prior, *f* ~ *N* (*µ, K*), with mean *µ* and covariance matrix *K*. For simplicity, we assume that *µ*(*s*) = 0. The entries of the covariance matrix *K* are defined by a kernel function *K*_*s,s*_*′* = *k*(*s, s*^*′*^ *θ*) that characterizes the expected covariance between any pair of sequences *s* and *s*^*′*^ under the prior, where *θ* represents the parameters of this kernel function. We then obtain a vector of measurements *y* for a subset of sequences *S* ⊆ *A*^*ℓ*^ and assume that they are noisy Gaussian observations of their corresponding function values 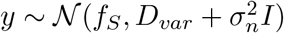, where *D*_*var*_ represents a diagonal matrix with the known variance of the experimental measurements and 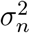 represents an additional learnable noise term. We first learn the hyperparameters *θ* that define the prior and the noise variance 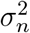 by maximizing the marginal likelihood:

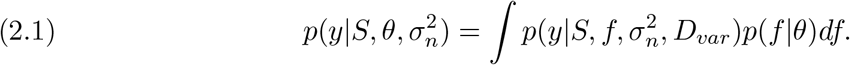

This integral can be evaluated analytically, making evidence maximization computationally tractable. In practice, we maximize the log marginal likelihood:

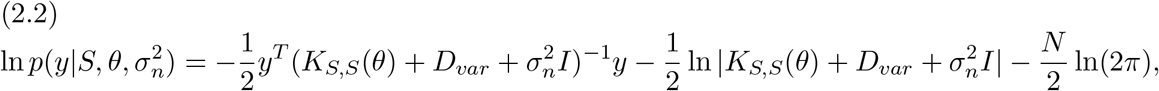

where *N* = |*S*|, and *K*_*S,S*_(*θ*) is the prior covariance matrix defined by the training sequences *S* and the hyperparameters *θ*. Let *S*_∗_ be the sequences in the test set and *f*_∗_ = *f* (*S*_∗_) be the function values of the test sequences. With the hyperparameters fixed at the values that maximize the log marginal likelihood, we can compute the posterior distribution *f*_∗_|*S, y, S*_∗_ of the function values at the test sequences *S*_∗_. We thus have *f*_∗_|*S, y, S*_∗_ ~ *N* (*m*_∗_, Σ_∗,∗_) where

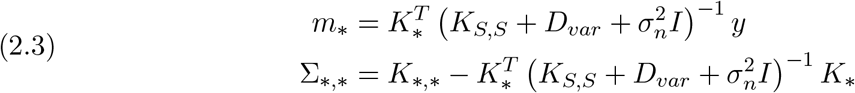

where 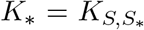 and 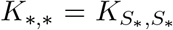. To then make predictions on the test set, we use the maximum a posteriori (MAP) estimate 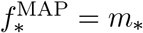.

## 3. Embedding kernels

An embedding kernel defines the covariance between two sequences as a decreasing function of the distance between the sequences in an embedding space. Let *σ*^2^ be a constant, *s, s*^*′*^ be sequences, and *ϕ* be an embedding function. We define the embedding kernel as

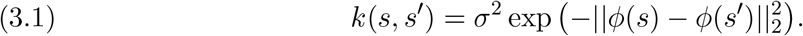

Here we consider linear embedding kernels, the special case when the embedding *ϕ* is a linear function of the one-hot representation of the sequence *s, x*(*s*): *ϕ*(*s*) = *Ax*(*s*) for a matrix *A*, also known as Factor Analysis (FA) kernels [75, 60]. The embedding matrix *A* can be inferred from data via evidence maximization or, alternatively, can be precomputed from patterns found in families of natural sequences. In this section, we discuss the expressivity and mathematical details of the model for learning linear embeddings *A*, but in Supplementary Materials Section SM1.8.1 we also explore embedding kernels that use precomputed linear embeddings derived from multiple sequence alignments and nonlinear embeddings derived from a protein language model [41].

### 3.1. Relationship to other kernels for sequence–function relationships

The embedding matrix *A* determines the arrangements of sequences in the embedding space. Constraining *A* restricts which arrangements of sequences are possible, giving rise to different kernel families with varying flexibility, some of which correspond to previously established models [60, 86]. We show that FA kernels with *A* restricted to being diagonal are Automatic-Relevance Determination (ARD) kernels, and FA kernels with *A* restricted to being block diagonal are General Product kernels (GPKs). We describe the geometry of the embedding space under these constraints on *A* and explain how removing the constraint on *A* allows for more flexible arrangements of sequences in the embedding space, enabling the model to potentially capture important features driving sequence functionality.

We define the ARD and GPKs before discussing their geometric intuition and connections to the FA kernel. The ARD kernel can be written as

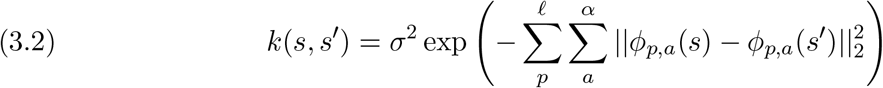

where 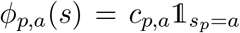 is a scaled indicator for whether allele *a* is present at position *p* in sequence *s*, and *c*_*p,a*_ is the learnable embedding scale parameter for each allele-position pair (*p, a*) that quantifies the importance of each pair. While the ARD kernel learns one parameter per allele-position pair, the GPK learns a covariance matrix for each position that expresses the extent to which changing between each pair of alleles at that position impacts the predictability of the phenotype. The covariance between a pair of sequences is the product of the position-specific covariances:

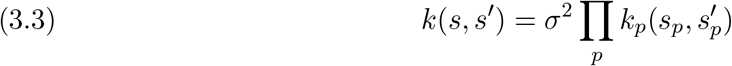

where 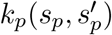 is the position-specific correlation at position *p* between alleles *s*_*p*_ and 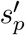.

For an FA kernel with a diagonal matrix *A*, each axis in the embedding space corresponds to a position-allele combination. The embedding coordinate for a sequence along the axis corresponding to the position-allele pairs (*p, a*) is determined by the presence or absence of allele *a* at position *p*. The FA kernel with *A* = *cI*, a diagonal matrix with *c* along the diagonal, is the Radial-Basis Function (RBF) kernel, an ARD kernel with the same lengthscale for each position-allele pair (*p, a*). The kernel value for a pair of sequences depends only on their Hamming distance and is therefore equivalent to a GPK where each position-specific correlation is weighted equally, 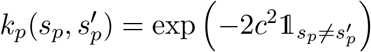, giving us exactly the Geometric model [48, 88, 86, 33]. Figure 1 (left) illustrates the embedding of this kernel where each single mutant is at distance *c* away from the wild-type along the axis corresponding to the mutant allele. For simplicity, we do not illustrate the axes associated with the wild-type alleles, at which single mutants away from the wild-type are also separated by distance *c*, resulting in a contribution of 2*c*^2^ to the squared distance for each mutation. The embedding of the double mutant *ab* is the vector sum of these two single mutant embeddings, geometrically completing the square.

**Figure 1.**
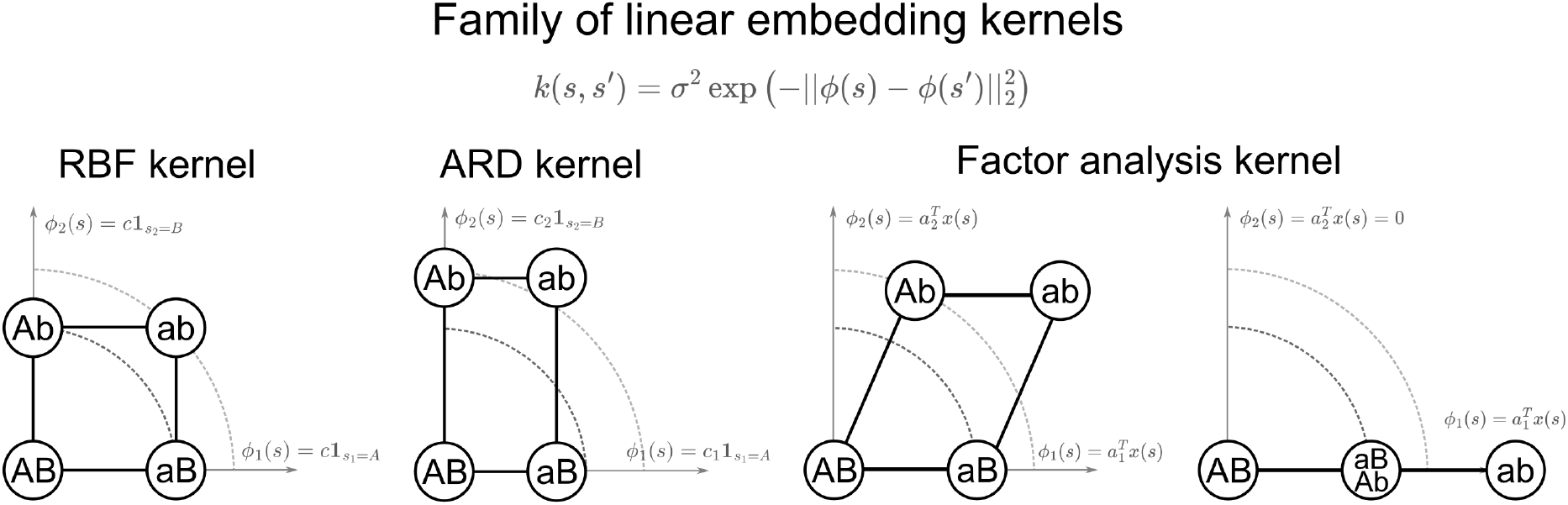
Geometric flexibility that FA kernels and ARD kernels. The axes of the RBF kernel and ARD kernel are defined by the scaled indicator of the presence of the sequences’ alleles at each position. The dotted circles of various radii are contours of k(·, AB). The RBF kernel sets all parameters c_p,a_ = c for all positions and alleles and the ARD example in this figure just distinguishes the parameter based on positions, c_p,a_ = c_p_. This restricts the embedding of the double mutant ab, embedded at the completion of the rectangle of the two single mutant axes. The FA kernel defines its axes as an inner product between a factor a_i_ and one-hot encoded sequence x(s). This enables more flexibility in embedding the double mutant by changing the angle between the two single mutant axes.

When we restrict *A* to be diagonal but allow the entries on the diagonal to vary, the FA kernel becomes an ARD kernel. Under this assumption, each position-allele embedding scale, *c*_*p,a*_, corresponds to the diagonal entry of *A*. Geometrically, the distances along the embedding axes now vary depending on this diagonal entry, allowing us to embed different alleles at different distances from the origin depending on which position was mutated. In Figure 1 (middle left), *aB* is embedded closer to the wild-type than *Ab*, so that the phenotype of the single mutant *aB* is expected to be more similar to the wild-type *AB* than the other single-mutant *Ab*. While learning different embedding scale parameters for different positions and alleles provides more flexibility with the single mutant correlations, the location of the double mutant is still geometrically the completion of the rectangle, and therefore completely determined by the covariance of the single mutants that comprise it.

The diagonal *A* case corresponds to two classes of product kernels. When *A* is a diagonal matrix whose entries are *c*_*p*_ for each position *p* regardless of allele, i.e. *A* = diag(*c*_1_*I*_*α*_, …, *c*_*ℓ*_*I*_*α*_), it forms a Connectedness kernel with 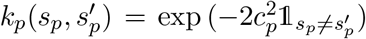. For a bi-allelic alphabet this is equivalent to the Connectedness model for fitness landscapes proposed by Reddy and Desai [61]. When *A* is a diagonal matrix whose entries depend on positions and alleles, *c*_*p,a*_ along the diagonal, it forms a Jenga model [86] 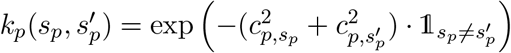.

Next we consider FA kernels with block diagonal embedding matrices *A*. In this case, each position retains its own *α*-dimensional embedding subspace, but its coordinates can now depend on arbitrary linear combinations of the allele indicators at that position rather than only individual alleles. The embedding along each of these axes can be any linear function of the one-hot encoding at that position. This defines a special case of GPKs with strictly positive entries, 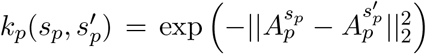 where *A* is the diagonal block of *A* at position *p* and 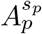 is the column of *A*_*p*_ corresponding to allele *s*_*p*_. Note that not all GPKs can be represented in this form, for instance, those where site-specific kernels contain negative entries [86].

Finally, we consider the most general FA kernel, i.e. one with no restrictions on the embedding matrix *A*. General FA kernels are not GPKs because each embedding dimension can combine contributions from alleles at multiple positions, rather than being restricted to a single position. This design enables FA kernels to capture sets of interactions that the ARD kernels and GPKs cannot. To clarify this, we first can rewrite the GPK in Equation 3.3 to reflect the contribution that each position has to the overall covariance via decay factors *δ*, where 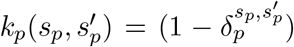 for position *p* and sequences *s* and *s*^*′*^. This enables us to rewrite the FA kernel in terms of these factors depending on individual positions and pairs of positions, highlighting this additional complexity

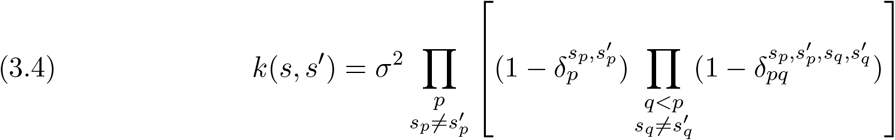

where:

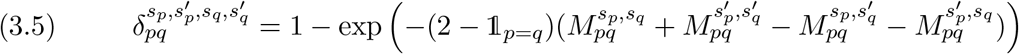

and 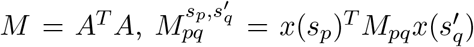 for block matrix *M*_*pq*_ at position pair (*p, q*), and 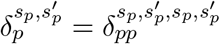. From this perspective, if we let 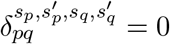, we recover a GPK in which different mutations have a constant multiplicative contribution to the covariance between two sequences given by their mutation-specific decay factors 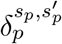. Thus, the mutation-pair specific 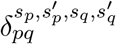 factors control the deviation from this multiplicative expectation positive 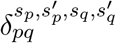, where values results in a smaller correlation between pairs of sequences than expected by the mutation specific contributions, while negative 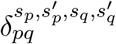 values result in higher correlation values. Note that the mutation pair specific 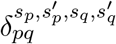 factors cannot take arbitrary values but are jointly constrained by *M* being positive semi-definite. For more details on the derivation, see the Supplementary Materials Section SM1.3.

Thus, the most general FA kernel can be understood as a relaxation of a subclass of GPKs. This additional flexibility can be interpreted geometrically, as shown by Figure 1 (right). With the unrestricted FA kernel, each axis is defined by an additive function of the sequence. This enables each dimension of the embedding space to depend on the genotype at multiple positions, which provides flexibility for different configurations of the double mutant in the embedding space. Now single mutant embeddings no longer need to lie on orthogonal axes. On the left side of the FA plot from Figure 1, the two single mutants are embedded so that they form an acute angle, rather than being orthogonal, so that the double mutant is further, geometrically completing the parallelogram. On the right FA plot in Figure 1, the angle has completely vanished to describe a scenario in which the single mutants *Ab* and *aB* are determined only on the first factor. As a consequence, the double mutant is at twice the distance from the wild-type in the embedding space than the single mutants. The FA kernel’s additional expressivity comes from the ability to model covariance among double mutants and the wild-type in a way that is not completely determined by the covariance between the single-mutants and the wild-type.

We have thus far seen that the unrestricted FA kernel provides maximal flexibility. However, its number of parameters grows rapidly with sequence length. The most general form of the FA kernel has 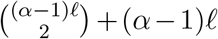 degrees of freedom, which is impractical in some settings. Therefore, sometimes we will consider the full-rank *A* and look at low-rank approximations of the learned *A* for interpretation, and sometimes we will directly learn a low-dimensional embedding space *A* with only a few factors, either with or without an added diagonal term to account for single-site, position-allele combinations.

### 3.2. Predictability of mutational effects

In this section, we characterize the flexibility of the FA prior to capture variation in the predictability of mutational effects across different genetic backgrounds [16, 2, 88, 86], as well as the similarity between the effects of different mutations introduced in the same genetic background.

Let *p* : *c*_1_ → *c*_2_ denote the mutation from allele *c*_1_ to allele *c*_2_ at position *p* and 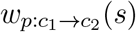 be the effect of introducing this mutation in the genetic background determined by the sequence *s*. Then, for a given sequence–function map *f* :

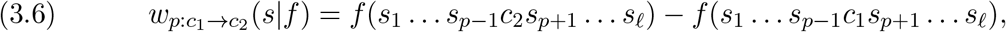

where *f* (*s*) is the phenotype of sequence *s*. In the following subsections, we consider different statistics that quantify the predictability of these mutational effects throughout the landscape under the FA prior.

#### Predictability of the effect of a fixed mutation across backgrounds that differ by a set of mutations

Here, we show that the expected predictability of a mutation’s effect across genetic backgrounds under the FA prior depends not only on the mutations separating those backgrounds, as with GPKs, but also on the specific mutation whose effect is being considered.

For a given sequence–function map *f*, we quantify the similarity between the effects of mutation *c*_1_ → *c*_2_ at position *p* across backgrounds differing only by mutation *c*_3_ → *c*_4_ at position *q*, with *p*≠ *q*, using the statistic

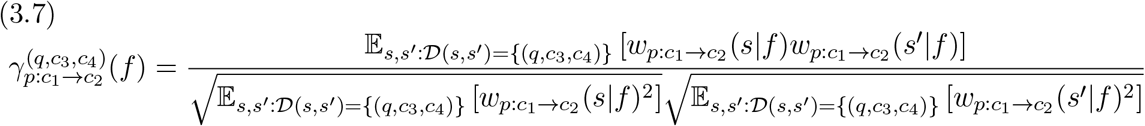

where 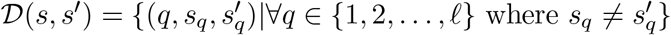 is the set of mutations at which two sequences *s* and *s* differ, and averages are taken uniformly over the indicated background pairs. This statistic averages the product of the paired mutational effects and normalizes it by the geometric mean of the mutation’s average squared effect across backgrounds containing either *c*_3_ or *c*_4_ at position *q*. Effects with the same sign in the presence of the *c*_3_ and *c*_4_ allele at position *q* contribute positively, whereas effects with opposite signs contribute negatively, with contributions weighted by their magnitudes. Unlike a centered correlation, this measure retains the contribution of effects that are consistent across backgrounds; in particular, it equals 1 when the focal mutation has the same nonzero effect in every background, whereas strict correlations are undefined in the absence of variability. These statistics correspond to cosine similarities between the vectors of mutational effects across the sets of backgrounds separated by one mutation and are closely related to the previously proposed *γ*(*A*_*i*_, *B*_*i*_)→(*A*_*j*_, *B*_*j*_) statistics [2] (see Supplementary Materials Section SM1.4 for details).

We approximate the expected value of this statistic under the FA prior, *f* ~ *N* (0, *K*), and express it in terms of the decay factors:

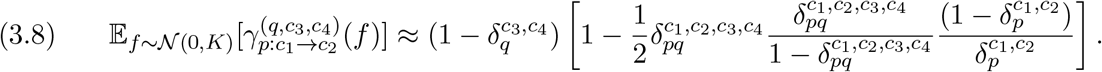

Throughout, ≈ denotes this approximation of the expectation of a ratio by the ratio of expectations [88]. Equation 3.8 illustrates the additional flexibility of FA kernels relative to GPKs. Under a GPK prior, 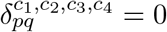, so the approximated expected value of the statistic depends only on the decay factor associated with mutation *q* : *c*_3_ → *c*_4_ separating the two backgrounds but not on the specific mutation *p* : *c*_1_ → *c*_2_. The FA kernel relaxes this restriction by allowing nonzero pairwise decay factors. Consequently, a given background mutation can have different effects on the predictability of different focal mutations, depending on the specific sites and alleles they involve. Interestingly, when the mutation-specific decay factors are fixed, the constraints on the FA decay factors imply that the FA prior cannot increase the predictability of mutational effects across backgrounds differing by a single mutation, relative to the corresponding GPK. See Supplementary Materials Section SM1.4 for more details and a generalization of Equation 3.8 where backgrounds are separated by an arbitrary set of mutations *D*.

#### Predictability of the effects of two mutations when introduced in the same background

Here, we examine the expected similarity between the effects of different mutations introduced in the same genetic background under the FA prior. We first consider the similarity in the effects of mutations at the same position, for which FA kernels and GPKs offer comparable expressivity. We then consider the similarity in the effects of mutations at different positions, where the FA prior provides additional flexibility.

For a given sequence–function map *f*, we quantify the similarity between the effects of mutation *p* : *c*_1_ → *c*_2_ and the effects of mutation *q* : *c*_3_ → *c*_4_ when introduced across all possible genetic backgrounds *s* containing alleles *c*_1_ and *c*_3_ at positions *p* and *q* using the following statistic

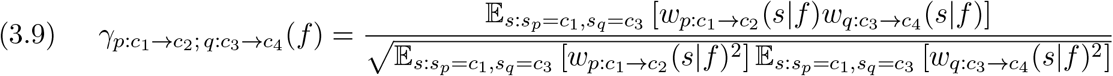

As before, this statistic also corresponds to the cosine similarity between the two vectors of mutational effects across backgrounds, and takes value 1 when the effect of one mutation is positively proportional to the effect of the second mutation and value 0 when these vectors are orthogonal to each other.

We approximate its expected value under the FA prior by replacing each background-averaged product or squared effect in Equation 3.9 with its expectation under the prior, as before. The results depend on whether the two mutations are introduced in the same site or across different sites. In the first case, given by *p* = *q* and *c*_1_ = *c*_3_, we redefine *c*_3_ to be the second mutant allele and find that

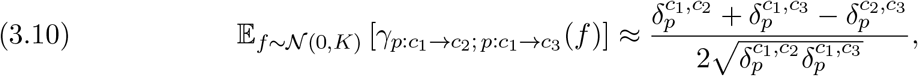

which depends only on the mutation specific decay factors at site *p* involving all pairs of the involved alleles, but not on the decay factors for pairs of positions. From this perspective, the FA kernel is as expressive as the GPKs, and both allow the effects of different mutations at the same site to be exactly proportional to each other when 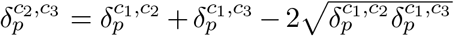, assuming that those decay factors are compatible with a valid kernel.

If we consider mutations at different positions *p≠ q*, we find that

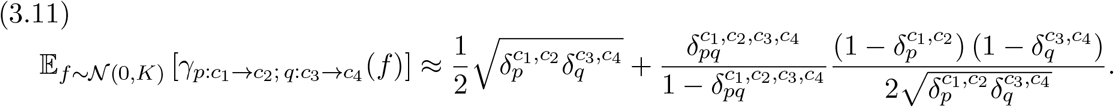

This equation reflects the additional flexibility of the FA prior relative to GPKs. Under a GPK prior, the pairwise decay factors vanish, and the approximation to the expected similarity between the effects of different mutations reduces to 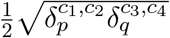. For GPKs with nonnegative site-specific correlations, the mutation-specific decay factors lie between 0 and 1, so this expression is bounded above by 0.5. This bound is approached as both decay factors approach 1, when sequence phenotypes are independent and identically distributed Gaussian variables, as in the classical House-of-Cards limit [37, 86]. Although the phenotypes are independent in this limit, two mutations introduced in the same sequence share the initial phenotype that is subtracted when calculating their effects. This shared term induces a positive association between the effects, yielding the value 0.5 in our approximation. For fixed single-mutation decay factors, the FA prior allows this approximate expected similarity to increase when 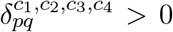 or decrease when 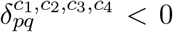, relative to the GPK base-line. The kernel embeddings provide an intuitive explanation for this flexibility. When two mutations produce the same displacement in every embedding dimension, introducing either mutation into a shared background leads to identical embedding coordinates and, therefore, identical mutational effects under the prior. In contrast, within the embedding representation of GPKs considered here, mutations at different sites contribute to disjoint sets of embedding dimensions, preventing this equivalence for mutations with nonzero effects (Figure 1).

### 3.3. Precomputed features

For each dataset we analyze, we will learn a linear embedding *A* applied to one-hot encoded sequences from lab-derived sequence–function data. Alternatively, embeddings can be precomputed from patterns observed in natural sequence data. Early work built such embeddings from structural or local context-dependent information: [63] augmented one-hot encoded sequences with a residue-residue contact map to capture pairwise structural features, while [83] broke sequences into non-overlapping k-mers and embedded them using a doc2vec-style approach. More recent approaches pair precomputed embeddings from protein language models [41], which capture the context of each residue against the entire sequence and are trained on much larger-scale datasets with a smooth kernel [4, 27, 24, 30]. Other methods instead use a different class of kernels that builds evolutionary or structural information directly into the kernel itself [24, 33].

Motivated by this work, we also consider precomputed embeddings from naturally occurring sequences or from patterns identified by protein language models to compare with our learned FA embedding and determine whether patterns derived from natural sequence variation offer information beyond what is learned directly from mutagenesis data. We consider two approaches for constructing such embeddings: extracting directions of greatest variation from a multiple sequence alignment (MSA) [3], or using embeddings derived from a protein language model [41]. We then define kernels using these precomputed embeddings as follows. Let *X* be a matrix containing the one-hot encodings for the sequences in a MSA or the pre-computed language model embedding for the set of experimentally characterized sequences. Then, *V*_*k*_ denotes the matrix containing the top *k* right singular vectors of *X*. We define the FA kernel using the embedding 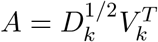 where 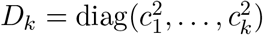 is a diagonal matrix containing the learnable *c*_*i*_ scale associated with the embedding dimension *i* specified by each row of the precomputed 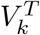. We apply these methods in Supplementary Materials Section SM1.8.

## 4. Examples of what the FA kernel can encode

Here, we illustrate how the additional flexibility of the FA embedding space allows the prior to encode covariance structures that reflect biological phenomena but cannot be expressed by site-factorized GPKs. We first show how a low-dimensional FA embedding can represent global epistasis, as previously demonstrated by LANTERN [73], and then illustrate how shared embedding dimensions can encode patterns of expected functional similarity between sequences that reflect RNA base-pairing interactions.

### 4.1. Global Epistasis

Global epistasis (GE), also called non-specific epistasis, describes the phenomenon in which the effect of a mutation changes systematically with the phenotype of the background in which it is introduced, rather than depending on the specific allelic composition of that background [50, 13, 71]. This arises when mutations combine additively with respect to an unobserved or latent phenotype *ϕ*(*s*), such as folding or binding affinity, which is nonlinearly related to the measured phenotype *y* = *g*(*ϕ*(*s*)) + *ϵ* [50, 13] for some noise *ϵ*. In practice, *g* is typically constrained to be monotonic to ensure identifiability the nonlinear function and the additive latent phenotype [50, 65].

The FA kernel can encode a similar type of model by letting the embedding space have a single dimension. In this case, the matrix *A* has a single row and can then represent the latent additive function *ϕ*(*s*) = *Ax*(*s*) and capture the nonlinear relationship *g* through the Gaussian process defined on that space. Note that this construction does not constrain *g* to be monotonic, but instead favors smooth *g* functions. When the embedding space *A* has more than one row, then it extends *ϕ* to a vector of additive latent traits. As a result, the model can learn nonlinearities in a multidimensional latent space without assuming monotonicity. Related approaches such as LANTERN show that learned latent representations can capture global nonlinearities in sequence–function landscapes [73]. We verify this behavior on simulated data (Supplementary Materials Section SM1.6) and observe similar latent structure in the GB1, AAV2, and GFP datasets (Sections 6.1, 6.3, 6.4). Because the FA kernel and LANTERN allow flexible nonlinear functions of additive latent traits, they impose fewer restrictions on the form of the nonlinear relationship than biophysical models, such as the commonly used two- or three-state thermodynamic models [49, 15] or Fisher’s geometric model [17, 72], where the form of the nonlinear relationship is assumed to have a specific parametric form.

### 4.2. RNA base-pairing

A second example in which the FA kernel’s additional flexibility allows the prior to encode useful relationships between sequences concerns base-pairing interactions in RNA. RNA function often depends on secondary structures maintained by base-pairing between specific positions. Different allele combinations can preserve these interactions and therefore support similar functions. In such cases, we may expect sequences separated by two compensatory mutations that preserve base-pairing to be more functionally similar than sequences separated by either of the individual mutations that disrupt it. GPKs with nonnegative site-specific correlations require the prior covariance to decrease or remain unchanged with each additional mutation, preventing them from directly encoding this expectation. In contrast, the FA kernel allows alleles at different positions to contribute to shared embedding coordinates, so that sequences at Hamming distance two can be close to each other in the embedding space (Figure 1). This way, sequences with complementary alleles, including canonical Watson–Crick *AU/UA* and *GC/CG* pairs, or wobble *GU/UG* pairs, can be placed close together while separating them from noncomplementary configurations.

We can consider an example of an embedding matrix that maps sequences with base pairs at a fixed pair of positions to the same point in the embedding space and the sequences without base pairs dispersed along the axes. Let *s* be a sequence of length two. We assume the one-hot encoding of *s* via *x*(*s*) is ordered *ACUG*. In one dimension, we define this embedding matrix in Figure 2A as:

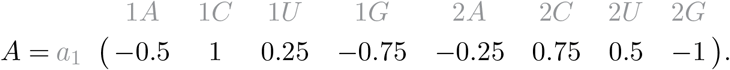

**Figure 2.**
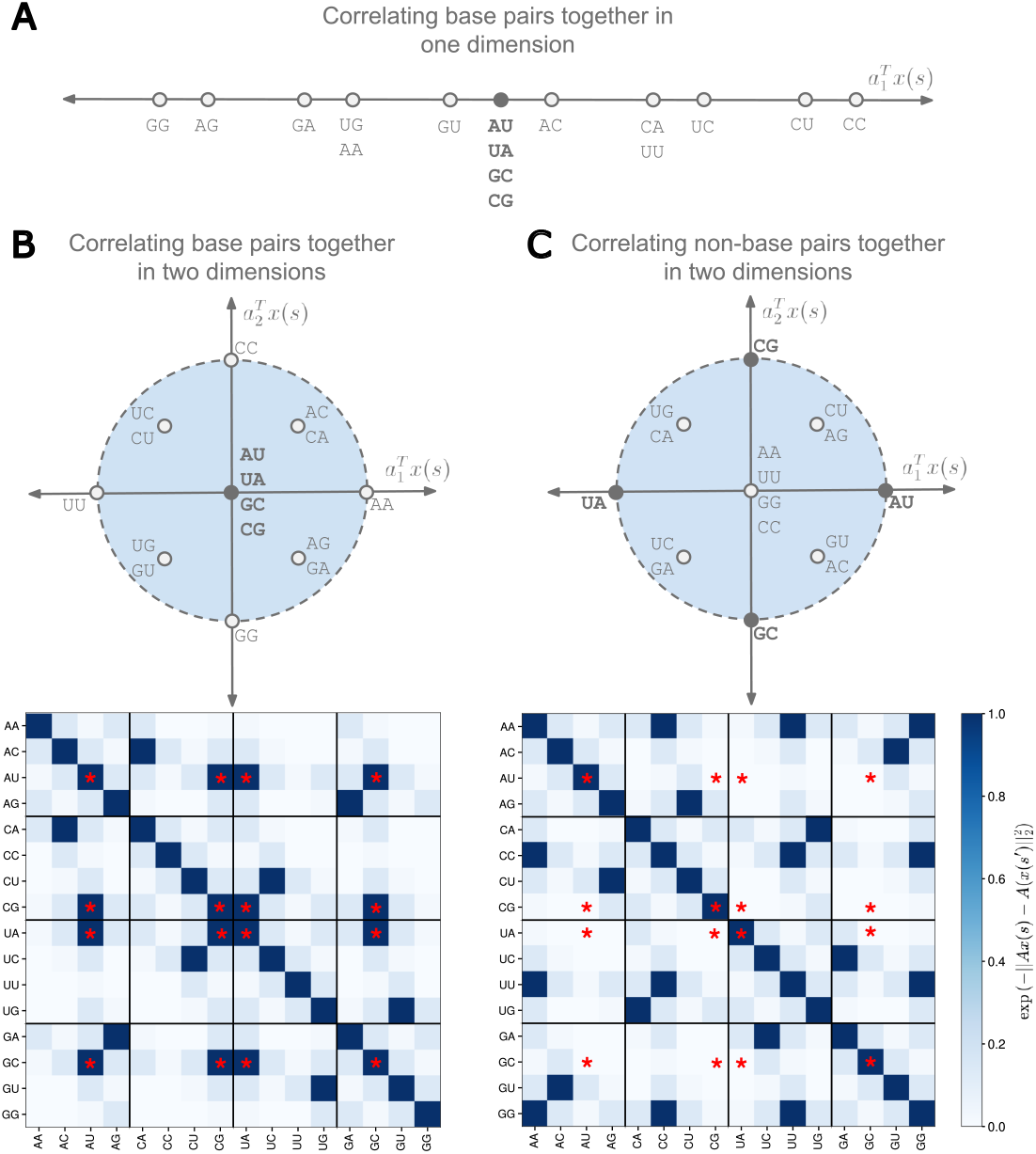
Embedding RNA sequences so that base pairs are separate from non-base pairs. (A) Top plot shows the embedding space in one dimension of all the sequences where all the base pairs (AU, UA, GC and CG) are embedded together at the origin and all non-base pairs are spread along the axis. (B) Now we demonstrate embedding base pairs together at the origin in two dimensions. Each factor now defines AU,UA and GC,CG pairings. This gives rise to a correlation matrix where all the base pairs (denoted by the red star) are correlated to one another. (C) Now each base pair is embedded onto a particular axis rather than at the origin. This gives us a correlation matrix where the base pairs are nearly uncorrelated with one another except of course when the two sequences are the same.

Here, we embed sequences that base pair at the origin so that they are completely correlated with each other. On the other hand, the non-base pairs are spread across the axis in both directions. By parameterizing this matrix with variables *a, b, c, d* in place of 0.5, 1, 0.25, 0.75 respectively, we can change the extent to which the non-base pairs are correlated with the base pairs. Furthermore, we observe a similar embedding structure in one of the four datasets we study (Section 6.2), highlighting the importance of base-pairing configurations in RNA structures.

We can extend this concept to higher dimensions for added expressivity. In Figure 2B, we define an embedding matrix with two factors:

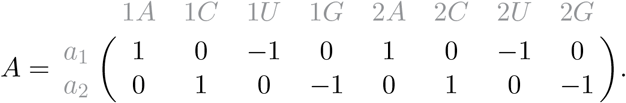

where the first row of *A, a*_1_, encodes the *AU* and *UA* pairs and the second row of *A, a*_2_, encodes the *CG* and *GC* pairs. The non-base pairs are distributed around the origin within a circle of radius two, creating distinct distances from the base pair configurations.

It is also possible to design an embedding matrix where different base pair configurations have low correlations under the prior, allowing the functional effects of the different base pairs to vary more independently. For instance, we can define the *AU/UA*] pairs on the positive and negative directions of the x-axis at [2 0] and [−2 0 and *CG/GC* on the positive and negative *y*-axis directions at [0 2] and [0 −2], respectively. The remaining nucleotide combinations lie closer to the origin than these base pairs. Sequences with the same nucleotide at both positions are mapped to the origin and therefore have perfect prior correlation with one another, while the other non-base pair configurations lie at the corners of a square centered at the origin.

This embedding can be achieved by switching the sign of position 2 in the *A* matrix above:

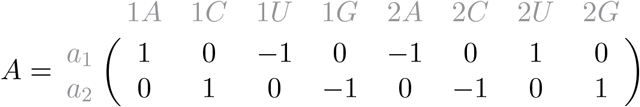

which we can visualize in Figure 2C.

Just as we can scale these embeddings in the single dimension to change the magnitude of the correlation, we can also do so in these two-dimensional examples. The added flexibility of multiple dimensions illustrates how more factors can encode more complex relationships between sequences.

## 5. Optimizing and interpreting FA factors

To learn the optimal embedding matrix *A* and interpret the corresponding factors, we must address two sources of nonidentifiability: the non-uniqueness of the embedding representation and additional gauge freedoms introduced by one-hot encoded sequence spaces [59].

Consider the alternative representation of the FA kernel:

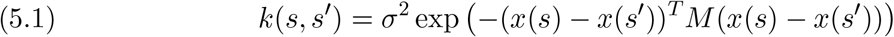

where 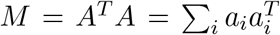 is a positive semi-definite matrix and *a*_*i*_ are the factors. The kernel is completely determined by the matrix *M*, rather than any particular factorization of *M*. There are many decompositions *M* = *A*^*T*^ *A*, meaning that there are many embeddings *A* that can provide the same distance structure and hence the same kernel. This non-uniqueness poses a challenge in optimizing and interpreting the factors.

### 5.1. Optimizing FA factors

To optimize the factors, we remove the redundant directions in which the parameters change but the kernel does not. These directions, known as gauge freedoms [59, 58], can hinder optimization. For one-hot encoded sequences, this gauge freedom arises because there are multiple *M* matrices that can result in the same covariance for sequences *s* and *s*^*′*^. The learned representation is not uniquely identifiable because adding an arbitrary offset to all allele coefficients of a factor at any given position does not change the kernel value: 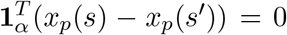, where *x*_*p*_(*s*) ∈ ℝ^*α*^ describes the vector indicating each allele at position *p*. Thus, translating the allele coefficient vectors at individual positions produces the same kernel (see Supplementary Material Section SM1.2).

We remove this gauge freedom by selecting a representative from the equivalence class of matrices *M*. Specifically, we impose a zero-sum gauge by requiring the learned coefficients *a*_*p,b*_ (the position-allele entries of *A*) to satisfy 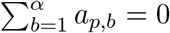 for each position *p*. This constraint is enforced by applying a gauge-fixing transformation to *M* : 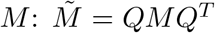, where *Q* is block diagonal with *ℓ* blocks 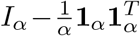 along its diagonal. This projection removes the components of the allele coefficient vectors in the all-ones direction, leaving only the zero-sum components. Therefore, rather than learning an arbitrary *M*, we learn a gauge-fixed representation 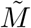.

A gauge-fixed matrix 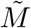 is rank deficient because *Q* removes the all-ones directions. Therefore, we parameterize the matrix in a lower-dimensional gauge-fixed subspace and lift it back to the original space. We define *P* ∈ R^*αℓ×*(*α*−1)*ℓ*^ by taking the eigendecomposition of *Q* so that its columns define a basis for the column space of *Q*. The matrix *P* is a basis for the gauge-fixed subspace, so it lets us parameterize a full-rank matrix in the reduced (*α* ™ 1)*ℓ*-dimensional space before lifting it back to the original parameter space. Letting *Q* = *PP* ^+^, where *P* ^+^ is the left pseudo-inverse of *P*, we can rewrite 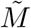 as 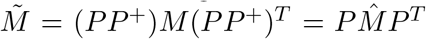, where 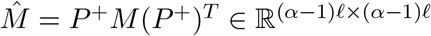.

With the *M* matrix expressed in the reduced gauge-fixed space, we can parameterize 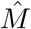 directly and recover the embedding matrix *A*. We consider two approaches for parameterizing 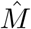. First, when we want to learn a full-rank 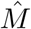, we parameterize the Cholesky factors *L* using the log-Cholesky parameterization [56] so that 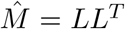 is symmetric and positive definite. This gives us our embedding matrix *A* = *L*^*T*^ *P* ^*T*^ and consequently our gauge-fixed and positive semi-definite matrix 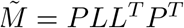 defining the kernel. Second, when we want to learn a low-rank 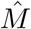, we instead parameterize the reduced 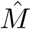 matrix through a set of orthogonal latent directions and positive scaling parameters. We learn an unconstrained matrix whose QR decomposition provides orthonormal directions *U*, and independently parameterize positive scales collected in the diagonal matrix, Λ. This yields 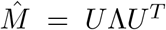 with corresponding embedding matrix *A* = Λ^1*/*2^*U* ^*T*^ *P* ^*T*^ and gauge-fixed and positive semi-definite matrix 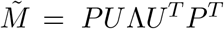.

### 5.2. Interpreting FA factors

Since the embedding matrix *A* is not uniquely determined by 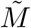, we must be cautious when interpreting individual factors. In particular, rotations of *A* produce equivalent embeddings that preserve the distance structure of 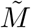. Therefore, to systematically decompose 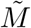, we take the eigendecomposition 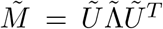, where the embedding matrix is 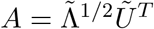. The eigenvectors of 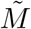 define orthogonal directions in the space of one-hot sequence encodings, with larger eigenvalues corresponding to directions along which the kernel similarity decreases more rapidly.

Although the eigendecomposition provides a systematic representation, the resulting orthogonal directions may not be the most interpretable directions to understand biological features. We also consider sparse decompositions of 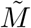 to evaluate whether the embedding space can be described by a set of axes, each depending on a small number of positions. To do so, we modify a sparse matrix decomposition method [53] to penalize position-allele blocks.

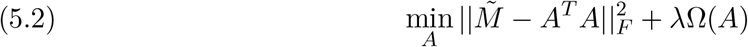

where the sparsity penalty is:

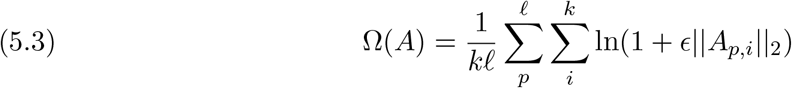

for parameters *ϵ* and *λ*. This cost function penalizes position-allele blocks for each factor or embedding dimension *i, A*_*p,i*_, where 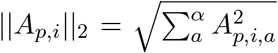. This penalty encourages sparse factors, influenced by only a small number of dominant positions.

To interpret the learned factors via sparse representations of 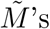 decomposition, we first apply a rotation test [53] to determine whether the sparsity exhibited by the optimizer of Equation 5.2 reflects sparse structure inherent in the learned matrix 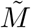. A random rotation of 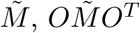, simply changes the basis of the eigenvectors of 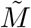. Therefore, refitting Equation 5.2 with the same parameters to the rotated matrix tests whether the apparent sparsity is tied to the specific position-allele coordinates. If so, the rotated 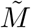 will have a larger reconstruction error for the same penalty and therefore support the existence of an underlying sparse decomposition in that coordinate system [53]. Of the three datasets in which we learned more than one latent dimension, only one exhibited evidence of a sparse decomposition as determined by this rotation test (Section 6.2).

## 6. Applying the FA kernel to data

In this section, we apply our method to four high-throughput experimental datasets, ranging from short sequence motifs in which we can reconstruct the phenotypes for every possible sequence in the space to full length proteins with over two hundred amino acids. We compare how the FA kernel performs in predicting the phenotypes of unobserved variants to other methods and how its estimated hyperparameters can be used to interpret the specific sequence–function relationships learned by the model.

### 6.1. GB1 landscape

We start by applying the FA kernel to the binding landscape of a four amino acid region in the protein G binding domain B1 with measurements of enrichment under selection for IgG-Fc binding for 149, 361 (93.4%) of the 20^4^ = 160, 000 possible sequences [80]. We use this previously characterized landscape to test the behavior of Gaussian process regression models under the FA kernel and illustrate how the parameters of the prior relate to our understanding of this landscape [87, 88, 86].

To do so, we apply the most general form of the FA kernel (defined by an arbitrary positive definite 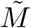) to this dataset and compare it with the other site-specific product kernels previously described [86]. In order to characterize the predictive power of these models under different training quantities, we sample sets of measured sequences of increasing size, fit the model’s hyperparameters via evidence maximization, and evaluate the predictive power of each model by computing the *R*^2^ between the predicted and observed measurements in the held-out measurements. Figure 3A shows that with sufficient amounts of training data, the performance of the FA kernel is similar to the GPK.

**Figure 3.**
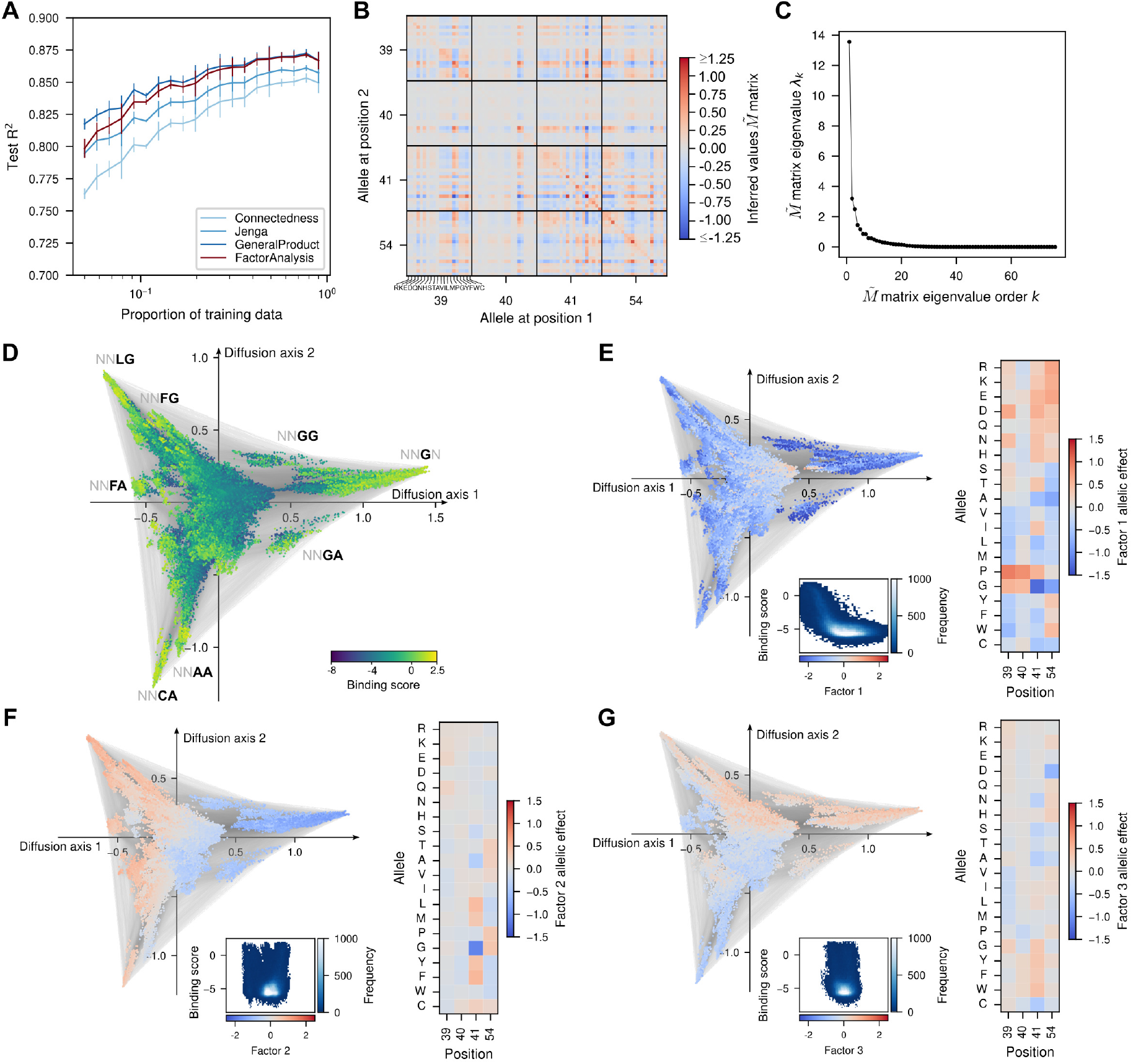
Application of Gaussian process regression with an FA kernel to the GB1 landscape. (A) Predictive performance evaluated by the R^2^ between the predicted and measured binding scores of held-out test sequences when using different amounts of training data for different models. Predicted values are the maximum a posteriori (MAP) estimate given by each method. Error bars represent the standard deviation across three different random samples for each fraction of training data. (B) Heatmap representing the entries of the 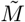 matrix estimated via evidence maximization under the FA kernel. Color scale is limited to values between −1.25 and 1.25 for visualization purposes. (C) Eigenspectrum of the learned 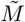 matrix, showing that the top three eigenvectors have the strongest effect. (D) Visualization of the predicted sequence–function map using the MAP estimate under the FA kernel on the complete dataset. Points represent genotypes and their color defines their predicted binding scores. Edges denote single mutations between genotypes. Squared distances between points optimally approximate the commute times between genotypes under a weak-mutation evolutionary model with selection for high phenotypic values [46, 44]. (E-G) Interpreting the FA kernel by overlaying the factor values from the top three eigenvectors of the 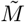 matrix on the commute time embedding from (D). Inset contains a two-dimensional histogram comparing the factor values with the posterior mean binding scores across all possible sequences. Heatmap on the right of each panel represents the independent contributions that each allele at each position has on each of the orthogonal factors.

Although the predictive performance of the FA kernel seems to be indistinguishable from the GPK, we can interpret the sequence–function relationship by analyzing the estimated hyperparameters of the prior. Figure 3B shows the learned 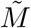 matrix under the FA kernel. The largest entry in this matrix corresponds to 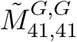, highlighting the importance of amino acid G at position 41 for the inferred covariance structure, consistent with previous findings [86]. The 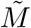 matrix found is noticeably different from the block-diagonal matrix assumed by a GPK, and suggests that sequences separated by two or more mutations may be correlated more or less than expected from the correlations between sequences separated by the individual mutations. Moreover, the eigenspectrum of the learned 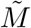 matrix indicates that this matrix may be well-approximated and interpreted by its top three eigenvectors (Figure 3C, Section 5.2). This result suggests that it may be reasonable to learn a lower rank matrix in other cases, especially for longer sequences (see Sections 6.3, 6.4).

To understand how the factors of the learned embedding matrix relate to the structure of the sequence–function relationship, we use the MAP estimate of the binding scores for every possible sequence to construct an evolutionary model in which selection favors higher values of the inferred binding phenotype. We then visualize the resulting landscape as a graph whose vertices represent genotypes and whose edges represent single mutations. The positions of the vertices are chosen so that squared distances approximate the commute times (the expected time to evolve from one sequence to another and back) between pairs of sequences under this evolutionary model (see [46, 44] and Supplementary Materials Section SM2.4 for details). This technique separates highly functional sequences into distinct peaks that are separated by regions of lower functionality and reveals three main regions of functional sequences, characterized by 41*G*, 41*A/C* with 54*A*, and 41*F/L* with 54*G* (Figure 3D) [87]. Then, we visualize each factor on top of this commute time embedding by specifying each genotype’s color as the factor value associated with that sequence. This enables us to explore how these learned factors relate to the known structure of the landscape and further interpret the independent effects of each allele at each position (Figure 3E-G).

Factor 1 shows typically low values in regions with high predicted binding scores, particularly for the 41*G* region at high values of diffusion axis 1 (Figure 3E). In fact, the inset at Figure 3E shows a very strong nonlinear relationship between Factor 1 and the predicted binding scores. Thus, Factor 1 may be interpreted as a latent additive function that is nonlinearly relative to the phenotype, capturing global or non-specific epistasis [50]. Moreover, Factor 1 shows strong negative contributions of the alleles that define the main sets of functional sequences i.e. *G, F, A, C* at position 41 and *A, G* at position 54, whereas alleles that are generally deleterious like *P, G* at positions 39 and 40 or the charged amino acids *R, K, E, D* at positions 41 and 54, show strong positive contributions.

Factor 2 aligns strongly with Diffusion axis 1 and clearly separates functional sequences characterized by 41*G* from those characterized by 41*F/L*, 54*G*, with intermediate values for sequences characterized by 41*C/A*, 54*A* (Figure 3F). While Factor 2 does not show any direct systematic relationship to the predicted binding scores, the three sets of functional sequences appear as three modes in the factor values at sequences with high predicted binding scores. Consistent with these observations, 41*G* shows a strong negative contribution to Factor 2, whereas 41*F/L* and 54*G* have strong positive contributions.

Finally, Factor 3 aligns with Diffusion axis 2, separating the third region of functional sequences characterized by 41*C/A*, 54*A* from the others (Figure 3G). Like Factor 2, Factor 3 does not show any clear relationship with the estimated binding scores. Its negative allelic effects for the amino acids *C* and *A* at position 41 and for *A* at position 54, among others, are nevertheless consistent with the sequence requirements of that region.

In summary, the learned FA embedding captures a pattern of global epistasis in the GB1 landscape and distinguishes the three regions of functional sequences previously identified by [87, 80]. This suggests that functional sequences form distinct clusters in the embedding space, which motivates applying the approach to landscapes that cannot be visualized via commute times but where groups of sequences with similar binding scores could still be identified as clusters in the embedding *A*.

### 6.2. Self-splicing intron

Next, we test the FA kernel on a sequence–function dataset that contains all 4^8^ possible sequence combinations at a key region in the *Tetrahymena thermophila* self-splicing group I intron [68, 45]. The phenotype is the experimentally reported log_2_ fold-change at 30^°^C. As before, we compare the predictive performance of the FA kernel and other models across a range of training set sizes (Figure 4A). Unlike what we observed in the GB1 dataset, the FA kernel in this dataset improves the prediction accuracy over the other product kernels when given enough training sequences to learn the hyperparameters accurately. This requirement to make accurate predictions reflects the FA kernel having significantly more hyperparameters to learn than the GPK (as described in Section 3.1).

**Figure 4.**
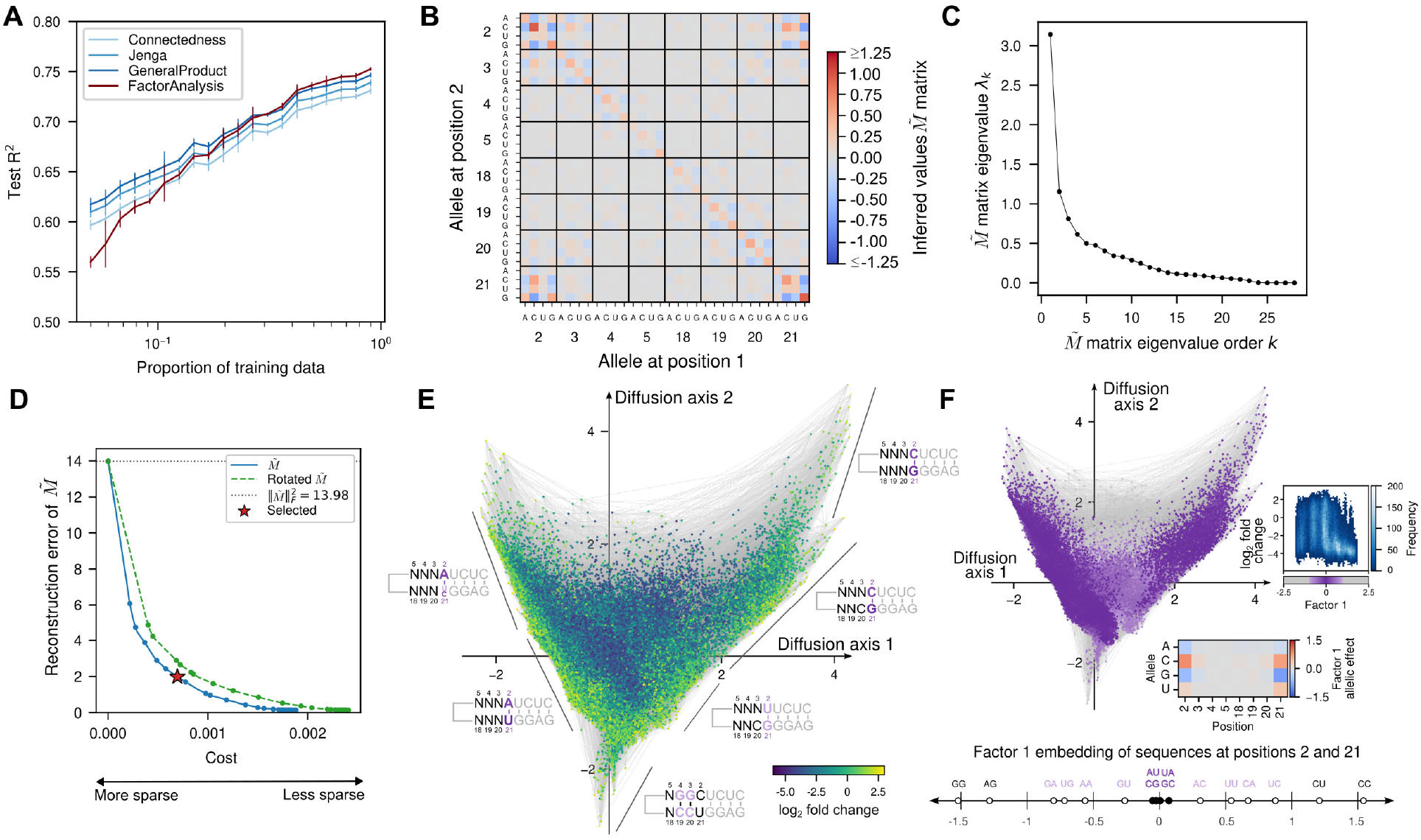
Application of embedding kernels to the self-splicing intron landscape. (A) Predictive performance evaluated by the R^2^ between the predicted and the measured phenotypic values of held-out test sequences when using different amounts of training data for different models. Predicted values are the maximum a posteriori estimate given by each method. Error bars represent the standard deviation across 3 different random samples for each fraction of training data. (B) Heatmap representing the entries of the 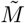 matrix estimated via evidence maximization under the FA kernel. Color scale is limited to values between −1.25 and 1.25 for visualization purposes. (C) Eigenspectrum of the 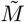 matrix, dominated by the first eigenvector. (D) Evaluation of the 12-component sparse structure decomposition (SSD) of the learned 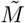 matrix. Comparison of the reconstruction error and the total cost function for a wide range of sparsity penalties of the 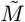 matrix (solid blue line) and a rotated version (dashed red line). The difference between the two lines supports the sparse decomposition [53]. Error is quantified by the squared Frobenius norm of the difference between the 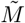 matrix and the reconstructed matrix from the sparse factors. The star denotes the embedding matrix A with 14% reconstruction error that was used to interpret the first factor of 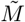 in panel (F). (E) Visualization of the predicted sequence–function map using the MAP estimate under the FA kernel on the complete dataset. Points represent genotypes and their color defines their predicted log_2_ fold-change values. Edges denote single mutations between genotypes. Squared distances between points optimally approximate the commute times between genotypes under a weak-mutation evolutionary model with selection for high phenotypic values [46, 44]. (F) Interpreting the FA kernel by visualizing Factor 1 values (the factor with the largest contribution in the sparse decomposition) on the commute-time embedding from (E). The color of each genotype is now associated with these factor values. Top inset contains a two-dimensional histogram comparing Factor 1 values with the posterior mean log_2_ fold-change across all possible sequences. Bottom inset heatmap represents the independent contributions that each allele at each position has on this sparse factor. The bottom axis shows the embedding coordinates for allelic combinations at positions 2 and 21, capturing base-pair configurations near zero with other allele combinations spread along the axis: 2G and 21G having the most negative value and 2C and 21C having the most positive value.

To understand the structure of the estimated prior, we investigate the structure of the inferred 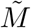 matrix (Figure 4B). We find that it has a strong block-diagonal structure corresponding to the individual sites, as we would expect to find under the simpler product kernels considered here. However, we also find large values at the off-diagonal blocks corresponding to the pairs of positions (2, 21) and (3, 20), known to be involved in base-pairing interactions for the P1 helix extension during the first step of the splicing reaction [68]. This indicates that the inferred 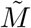 matrix recovers known pairwise interactions between positions without prior knowledge of the RNA structure.

The eigenspectrum of 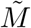 is dominated by its first eigenvector, and the first 12 eigenvectors account for 88% of its structure (Figure 4C). While the eigendecomposition of 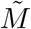 provides a systematic way to interpret the factors, we find that in this dataset, unlike the GB1 landscape, these orthogonal directions are less interpretable than a sparse decomposition of 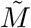, which yields factors that depend on fewer positions. This is consistent with the dataset having a few key sites that are biologically important, such as those involved in base-pairing.

To obtain a more interpretable embedding from this sparse decomposition, we constrain the embedding matrix *A* to have 12 components, corresponding to the top 12 eigenvectors of 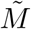, and penalize the overall contribution of each site to each embedding dimension (Section 5.2). To test whether the estimated 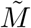 has an underlying sparse structure, we apply the rotation test [53] described in Section 5.2. If 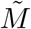 has a sparse decomposition, the rotated 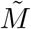 should have a larger reconstruction error at the same level of sparsity. This is what we find (Figure 4D), suggesting that the inferred 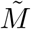 admits a decomposition that is sparse with respect to individual sites. We select the particular sparse decomposition (indicated by the star) with an error of 2*/*13.98 ≈ 14% described by the squared Frobenius norm, which balances the sparsity and structure of 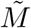 to highlight important position-allele combinations inherent in 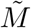.

We again use the MAP estimates under the FA model to construct an evolutionary model, this time assuming selection for higher values of the predicted splicing-related phenotype. We then visualize the resulting landscape using the same dimensionality reduction technique as for GB1 [46, 44] (Figure 4E). This visualization highlights regions of sequence space that correspond to different base-pairing configurations between positions 2 and 21, arranged according to the relative accessibility between them, with 2*C* and 21*G* and the reciprocal 2*G* and 21*C* base pairs at extreme values of Diffusion axis 1. We find that Factor 1, ordered first for having the largest variation of the sparse factors, has contributions from sites 2 and 21 only, and embeds sequences with base-pairing configurations near zero (Figure 4F, bottom). This finding recapitulates the expected patterns from Section 4.2 (Figure 2A). In addition to the highly interpretable contribution of Factor 1, the higher predictive power of this model provides greater resolution to the structure of the sequence–function relationships than previous estimates [45]. This improvement results in a visualization that more clearly shows how the different sets of functional sequences are connected to each other in sequence space. Moreover, the visualization highlights a set of functional sequences at low values for Diffusion axis 2, characterized by a typically deleterious combination of pyrimidines at positions 2 and 21 that can be compensated by the combination of 3*G*, 4*G*, 19*C*, 20*C*. This compensation could enable the formation of two strong, potentially compensating, *GC* base pairs, a mode of action that was not previously characterized to our knowledge (Figure 4E).

In summary, we find that the FA kernel can capture the base-pairing configurations that are most relevant for the self-splicing efficiency of the group I intron [77, 68], as anticipated by our theory. Moreover, the increased predictive power provided by this more informative prior enables the characterization of a local peak that could correspond to a new mode of action for the recognition of the 5’ splice site.

### 6.3. AAV2 capsid production

The previous datasets covered small, nearly complete combinatorial sequence spaces, which enabled us to predict the phenotype of every possible sequence. We could then understand how the latent dimensions relate to the structure of these sequence–function relationships by visualizing the factors on the commute-time embedding derived by assuming that natural selection favors greater phenotypic values [44]. Visualizing the landscape in this way requires computing a prediction for every sequence, which is infeasible for sufficiently long sequences. Instead, we can visualize and analyze the learned factors to understand the sequence–function map. This allows us to analyze longer sequences without needing predictions for all possible combinations of mutations.

Here, we consider a deep mutational scanning (DMS) dataset containing over 40,000 variants in a 28-amino-acid region (positions 561–588) of the AAV2 capsid protein [6]. The measured phenotype is a viral production score, defined as the log enrichment of each variant in the viral output library relative to the input plasmid library, with higher scores indicating more efficient production of genome-containing viral particles. Since constructing a commute-time visualization of the complete sequence space is infeasible, we explore how the learned factors under an FA kernel can be used to guide the interpretation of these high-dimensional maps, in a similar way to the mutation-specific decay factors under GPKs [86].

One challenge when fitting FA kernels to datasets composed of longer sequences is the large number of kernel hyperparameters, given by 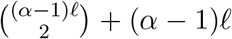. For example, a completely general FA kernel for protein sequences of length 28 would have a total of 141,778 hyperparameters, whereas the most general GPK has only 5,320. Since we observed in our empirical results that the GB1 and self-splicing intron datasets yielded 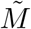 matrices whose eigenspectra were dominated by the first few eigenvectors, we estimate a rank-eight 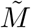 matrix for the AAV2 dataset. We find that the FA kernel’s predictive performance, with enough training data, is comparable to that of the GPK (Figure 5A). Moreover, the eigenspectrum of 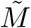 indicates that the eigenvectors with the largest eigenvalues capture the additive functions along which the covariance decays fastest (Figure 5B).

**Figure 5.**
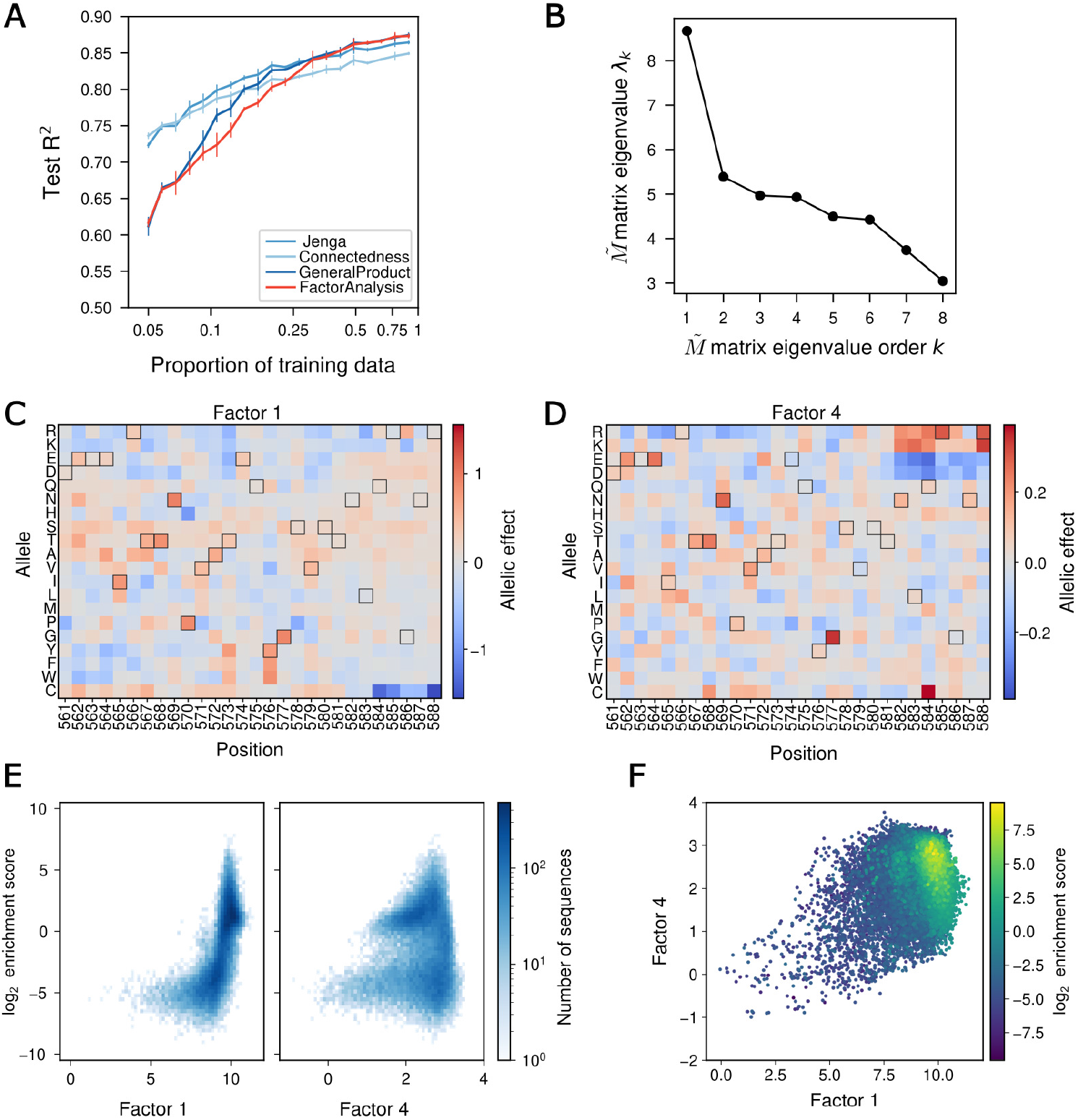
Applying the FA kernel to AAV2 capsid production. (A) Predictive performance evaluated by the R^2^ between the predicted and the measured viral production scores of held-out test sequences when using different amounts of training data for different models. Predicted values are the MAP estimate given by each method. Error bars represent the standard deviation across three different random samples for each fraction of training data. (B) Eigenspectrum of the estimated low-rank 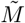 matrix. (C-D) Heatmaps representing the independent contributions that each allele at each position has for Factors 1 (C) and 4 (D). (E) Two-dimensional histogram comparing Factors 1 (left) and 4 (right) values with the posterior mean of the predicted log_2_ enrichment score across all measured sequences. (F) Scatterplot showing the relationship between the posterior mean of the log_2_ enrichment score and corresponding embedding values for Factors 1 and 4 across all measured sequences.

As with the self-splicing intron dataset, we tried a sparse decomposition of the 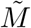 matrix to better interpret its factors. However, when applying the rotation test [53] to this decomposition, we find that the reconstruction error of the arbitrary rotations of the 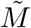 matrix are comparable to the reconstruction error of the non-rotated 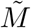 matrix (see Supplementary Materials Section SM3 for details). This suggests that there is no true underlying sparse structure in this case. Therefore, we instead use the eigendecomposition of 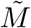 to interpret the factors via these orthogonal directions. Of the eight factors, we further analyze Factors 1 and 4, as their interpretations are the most apparent.

Factor 1, corresponding to the dominant eigenvector, shows independent allelic contributions with high values for alleles at 569*N* and 576*Y/F/W*, both of which were previously shown to be associated with high functional virus production (Figure 5C) [86]. We find, furthermore, that Factor 1 shows a strong nonlinear relationship with the predictions for the measured sequences (Figure 5E, left), consistent with a strong global epistasis pattern in which low embedded values are associated with low scores and high embedded values with high scores.

Among the next most relevant factors (Figure SM4 in Supplementary Materials), we find that Factor 4 shows a particularly interesting pattern of allelic effects. At positions 582–588, within a surface loop of the AAV2 capsid, positively charged amino acids (R/K) tend to have high embedding values, whereas negatively charged amino acids (E/D) tend to have low embedding values (Figure 5D). A charge-balanced combination of these alleles would therefore tend to result in intermediate values along this embedding axis, potentially capturing the previously proposed charge-balance hypothesis for this loop [86]. In addition to the main contribution of the positively and negatively charged residues, we also find large contributions from other specific alleles, such as 577*G* and 584*C* (Figure 5D), suggesting that these alleles may also have a role in the orientation or function of this particular loop. Consistent with the charge-balance hypothesis, comparing Factor 4 with the predicted viral production scores shows that both high and low values of Factor 4 are associated with reduced functionality, whereas intermediate values are associated with higher viral production (Figure 5E, right).

Although we cannot reconstruct the complete sequence–function relationship to compute the commute-time embedding as in the previous datasets, we can still use the learned factors as embedding dimensions for arbitrary sets of sequences. In the full embedding space, the prior covariance is proportional to the exponential of the negative squared distance between sequences. We therefore visualize the sequences by their Factor 1 and 4 values (Figure 5F) which allows us to see how the inferred phenotype varies across these two dimensions. This representation shows that both high values of Factor 1 and intermediate values of Factor 4 are associated with highly functional sequences.

In summary, the FA kernel provides comparable predictive performance to previous models while also learning factors that capture biologically meaningful patterns, including global epistasis and the previously identified charge-balance constraint on AAV2 capsid production.

### 6.4. Green fluorescent protein

In this section, we examine how the FA kernel can be used to learn sequence–function relationships defined over full-length protein sequences by applying it to deep mutational scanning data that reports the log fluorescence of green fluorescent protein (GFP) variants [65, 23]. This dataset comprises measurements of four libraries, each obtained by random mutagenesis of different wild-type GFP sequences (avGFP, amacGFP, cgreGFP, and ppluGFP2). These four wild-types differ from one another with varying degrees of divergence, with sequence identities ranging from 82% (avGFP and amacGFP) to 17% (amacGFP and ppluGFP2). Previous analyses of these datasets suggest that avGFP is well described by a simple global epistasis model with a single additive latent trait [50], whereas specific higher-order epistasis may contribute substantially to other datasets such as cgreGFP and ppluGFP2 [67].

To evaluate the performance of the FA model relative to simpler GPKs, we need to limit the number of hyperparameters of the priors for these 222 to 238 amino acid protein sequences. To do so, we restrict our analyses to the Jenga kernel, which contains one parameter per allele and position, and to an FA kernel with a one-dimensional embedding, which should be able to capture the patterns of non-specific or global epistasis. Furthermore, we also consider a kernel resulting from the product of the Jenga kernel and one-dimensional FA kernel that we call Jenga-FA, equivalent to an FA kernel with 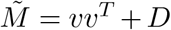, which generalizes both types of models. We find that the FA and Jenga-FA kernels match or improve predictive performance when compared with other product kernels in all four GFP datasets given a sufficient amount of data (Figure 6A). We find that the Jenga-FA kernel performs especially well relative to the FA and other models in the cgreGFP dataset in the high data regime. Interestingly, the estimated 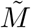 matrix under the Jenga-FA kernel is dominated by the diagonal, rather than the rank-one component, indicating that the model relies primarily on the position-specific structure of the Jenga kernel rather than additional interactions learned by the FA embedding. This suggests that specific epistasis captured by the Jenga model component may be particularly important in this dataset, consistent with the findings of [67].

**Figure 6.**
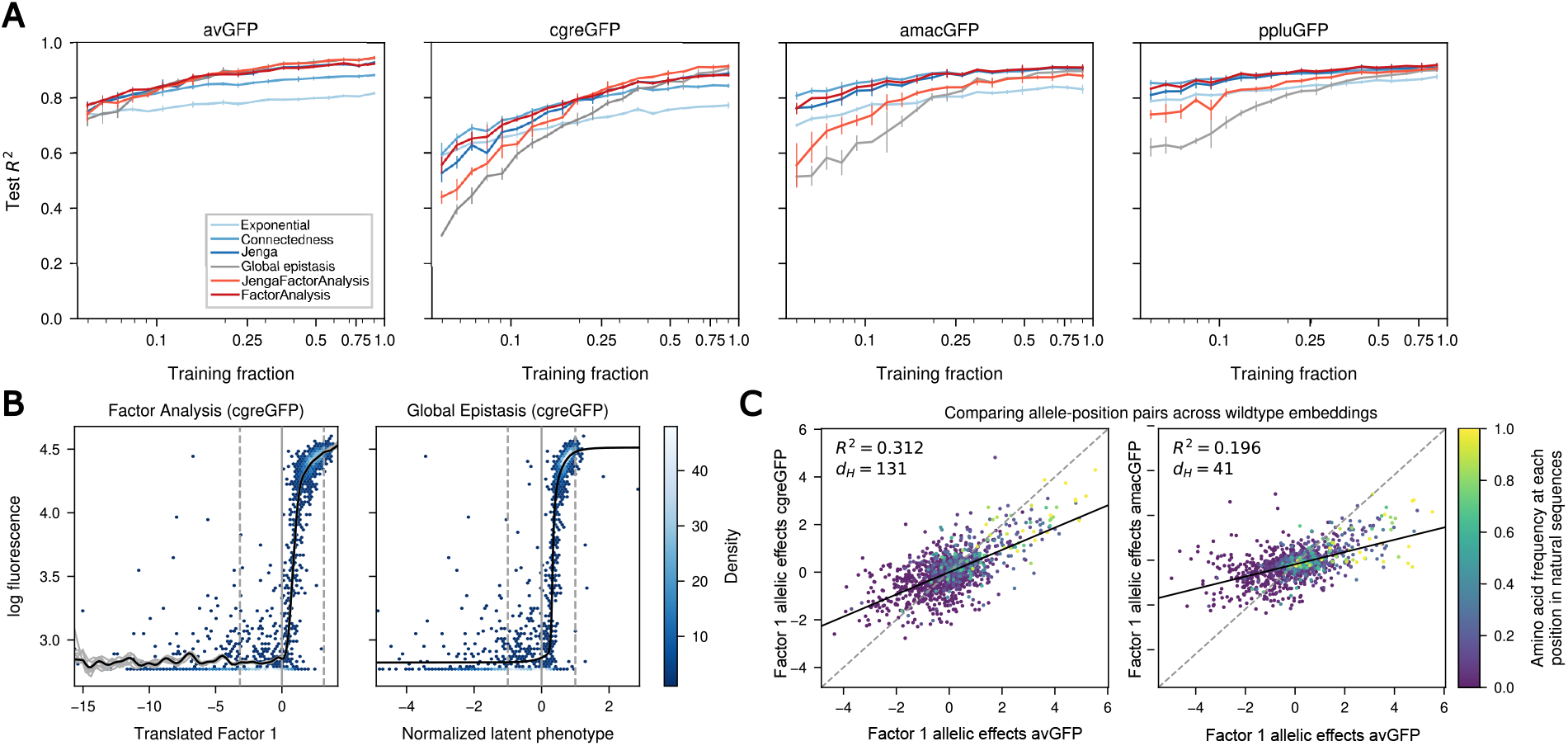
Application of the FA kernel to the analysis of four full-length protein datasets. (B) Comparison of the measurements and predictions with the translated inferred additive embedding (Factor 1) so that the mean is centered at the origin and latent phenotype under the FA kernel (left) and global epistasis (right) models, respectively, for held-out test sequences of the cgreGFP dataset. On the left, the grey lines represent samples from the posterior over the log fluorescence values of sequences with different values of Factor 1, and the solid black line represents the posterior mean plotted on the same sequences as the grey lines (see Supplementary Materials Section SM2.5 for more details). On the right, the solid black line represents the learned nonlinear function relating the additive latent phenotype with the predicted log fluorescence. (C) Scatter plots comparing the estimated allelic effects at each position between avGFP and cgreGFP (left) and between avGFP and amacGFP (right) in relation to the allele-position frequency of naturally occurring GFP sequences from a constructed MSA (see Supplementary Materials Section SM2.1.4 for more details). Dots represent specific allele-position combinations with color representing their frequency in the MSA. The dashed line represents y = x, the solid black line represents the least squares regression line, and d_H_ indicates Hamming distance between wild-type GFP sequences.

In Section 4, we hypothesized that the FA kernel could capture patterns of global or non-specific epistasis, which is known to explain a large fraction of the interactions in these datasets [50, 67]. Therefore, we also fit an additive latent trait global epistasis model [71] to these datasets (Supplementary Materials Section SM2.3) and compare its performance with the one-dimensional FA model. Notably, we see that the one-dimensional FA kernel tends to perform better than the global epistasis model (GE), particularly with smaller training sets (Figure 6A). This surprising improvement suggests that this GP framework learns more efficiently on limited data. Figure 6B compares the nonlinear relationships learned by these two models for the cgreGFP dataset, showing the fitted GE function alongside the posterior mean and posterior samples from the FA model (see Supplementary Materials Section SM2.5 for more details on how we sampled the sequences).

Thus far, we have analyzed the four different GFP datasets independently of each other. Therefore, to further understand the relationship between the learned priors across the different datasets, we compare the estimated independent allelic effects at each position that define the one-dimensional embedding at the homologous positions from an MSA [23] (Supplementary Materials Section SM2.1.4). We define the Hamming distance between aligned wild-types as the number of positions at which both sequences contain an amino acid and those amino acids differ, so positions with a gap in either sequence are excluded. This metric enables us to compare the similarity of wild-types in Hamming distance with their learned embeddings. Interestingly, while avGFP and amacGFP are closer to each other in Hamming distance (82% identity, Hamming distance of 41) than avGFP and cgreGFP (41% identity, Hamming distance of 131) [23], we find that the coefficients of the learned embeddings are more strongly correlated between avGFP and cgreGFP than between avGFP and amacGFP (Figure 6C). This suggests that similarities between learned representations of sequence–function relationships are not necessarily determined by overall sequence identity.

Finally, in Supplementary Materials Section SM1.8, we consider kernels that use embeddings derived from distributions of natural sequence data, rather than embeddings learned directly from the data as we have done here. We consider both linear and nonlinear embeddings. The precomputed linear embeddings are derived from the top *k* right singular vectors of the one-hot encoded sequences from a constructed MSA of natural GFP proteins (see Supplementary Materials Section SM2.1.4 for how it was created). The precomputed nonlinear embeddings are derived from the ESM protein language model [29, 41] on the lab-derived sequences in two different ways: (i) ESM embeddings averaged over positions and (ii) the top *k* right singular vectors of the per-position ESM embeddings. Our results show that both the FA kernel, as well as the other product kernels, performed substantially better than kernels using these precomputed features, even when trained on a small number of sequences. These results suggest that, for the GFP datasets considered here, learning simple site-specific or additive embeddings directly from the experimental data provides better predictive performance than using precomputed embeddings derived from natural sequence diversity or protein language models.

## 7. Discussion

In this work, we applied the FA kernel, a classical kernel on continuous spaces, to discrete sequence spaces to better understand sequence–function maps. We characterized the kernel as a generalization of ARD kernels and a subclass of GPKs and demonstrated its added expressivity in comparison to these existing kernel families. We describe our GPU-accelerated implementation that resolves the nonidentifiability of the embedding matrix, including its non-uniqueness and the additional gauge freedom introduced by one-hot encodings. Finally, we evaluated FA kernels on four diverse sequence–function datasets, spanning from short motifs to full-length proteins: the binding domain of GB1, the self-splicing group I intron, the AAV2 capsid assembly, and the fluorescence of GFP. Across these datasets, FA kernels learned latent representations that reflect known biological structure, including global epistasis for GB1, AAV2, and GFP; RNA base-pairing in the group I intron; and charge-balance constraints in AAV2. Comparing learned embeddings across GFP proteins also revealed greater similarity between avGFP and cgreGFP than between avGFP and amacGFP, despite the greater sequence divergence of avGFP and cgreGFP. Across all four datasets, FA kernels achieved predictive performance comparable to or better than GPKs, demonstrating that the learned embeddings both offer a means of interpretation and practical improvements in predicting sequence–function relationships.

Our work contributes to the ongoing integration of theoretical and empirical approaches to understanding sequence–function relationships, in which well-characterized families of random fitness landscapes [35, 1, 69, 31, 61] can be used as priors for inference from experimental data [87, 88, 86, 45]. In particular, we show that the classical FA kernel [60] generalizes a subclass of GPKs [61, 86], allowing the prior covariance between sequences to depend on combinations of mutations at different positions. From the perspective of the *γ* statistics, which describe how mutations affect the predictability of other mutations’ effects [16, 2, 88, 45], this prior allows the same mutation to have different effects on the predictability of other mutations. This variation can occur not only between mutations at different sites, as in [45], but also between alternative mutations at the same site. Unlike [45] and [86], the FA kernel can flexibly model heterogeneity in the predictability of the effects of different mutations from one another when introduced in a shared genetic background. These statistics, defined in Equation 3.9, describe another aspect of the local landscape geometry: the extent to which different mutations have similar effects. This provides a discrete analogue of the interpretation of the matrix *M* in terms of the second moments of local gradients when the function is defined on a continuous space [60, 81]. Associations among outgoing mutational effects underlie calculations of the expected number of local peaks in Gaussian random field models [31, 22]. Our work suggests an additional useful perspective: these covariances can help interpret the parameters of the FA prior in terms of relationships among mutational effects and guide exploration of the underlying sequence–function relationship.

Our FA kernels learn sequence space embeddings, connecting our approach to a line of work on continuous space embeddings for biological sequences [63, 83, 4, 24, 30, 33]. We investigated whether embeddings precomputed by an independent method could be used in place of learned embeddings by applying FA kernels with embeddings from the protein language model ESM rather than by a one-hot encoding. On the four GFP datasets, these models performed poorly (Supplementary Materials Section SM1.8), suggesting that Euclidean distances in the ESM latent space are not good proxies for fluorescence similarity. This could reflect misalignment between the masked language modeling objective of ESM and the assayed fluorescence phenotype. Alternatively, the ESM latent space may encode information related to fluorescence in a way that is not well captured by Euclidean distance. Learning embeddings as a linear transformation from site-specific embeddings derived from pretrained, structure-aware positional embeddings has been previously shown to improve Gaussian process regression performance [33]. Thus, learning nonlinear functions of the ESM embeddings involving all sites through deep kernel learning (DKL) [79] has the potential to extract further information. Comparing these models with their counterparts trained on one-hot encodings would help assess whether precomputed features provide useful prior information for learning a particular sequence–function relationship. Exploring these extensions and how their learned representations can be interpreted in terms of mutational effects provides an interesting direction for future research.

### Further Reading

The following references provide an introduction to some key mathematical topics relevant to this work: Gaussian processes and kernels [60]. Closely related work, including the LANTERN model [73], General Product kernels [86], and gauge-fixing [59, 58, 52]. Readers may also find it useful to read an overview of models that capture sequence–function relationships beyond Gaussian processes [38].

## Author Contributions

All authors have made substantial intellectual contributions to the study conception, execution, and design of the work. All authors have read and approved the final manuscript. In addition, the following contributions occurred: Conceptualization: Samantha Petti, Carlos Martí-Gómez; Methodology: Waverly Carabba, Carlos Martí-Gómez, David M. McCandlish, Samantha Petti; Formal analysis and investigation: Waverly Carabba and Carlos Martí-Gómez; Writing - original draft preparation: Waverly Carabba; Writing - review and editing: Waverly Carabba, Carlos Martí-Gómez, David M. McCandlish, Samantha Petti; Funding acquisition: Samantha Petti; Supervision: David M. McCandlish, Samantha Petti.

## Access to Code

The general code to infer sequence–function relationships using Gaussian process models under the Factor Analysis kernel is available on the EpiK GitHub repository: https://github.com/cmarti/epik. Code to recreate all the figures and the results presented in this manuscript is publicly available on a different GitHub repository: https://github.com/cmarti/embedding_kernels.

## Acknowledgments

SNP is supported by the Burroughs-Wellcome Fund’s Career Awards at the Scientific Interface (CASI). CMG and DMM acknowledge support from US National Institutes of Health Grant R35GM133613 with computational support from US National Institutes of Health Grant S10OD028632 and additional support from the Simons Center for Quantitative Biology at Cold Spring Harbor Laboratory. AI was used to help with basic coding of plots and KeOps implementation [7]. All code produced has been reviewed by the authors to ensure correctness.

## REFERENCES

[1] T. Aita and Y. Husimi, Fitness Landscape of a Biopolymer Participating in a Multi-step Reaction, Journal of Theoretical Biology, 191 (1998), pp. 377–390, 10.1006/jtbi.1997.0595, https://www.sciencedirect.com/science/article/pii/S0022519397905956 (accessed 2025-06-11).

[2] C. Bank, S. Matuszewski, R. T. Hietpas, and J. D. Jensen, On the (un)predictability of a large intragenic fitness landscape, Proceedings of the National Academy of Sciences, 113 (2016), pp. 14085–14090, 10.1073/pnas.1612676113, https://pnas.org/doi/full/10.1073/pnas.1612676113 (accessed 2024-01-09).

[3] A. R. Baxter-Koenigs, G. El Nesr, and D. Barrick, Singular value decomposition of protein sequences as a method to visualize sequence and residue space, Protein Science, 31 (2022), p. e4422, 10.1002/pro.4422, https://onlinelibrary.wiley.com/doi/10.1002/pro.4422.

[4] T. Bepler and B. Berger, Learning the protein language: Evolution, structure, and function, Cell Systems, 12 (2021), pp. 654–669.e3, 10.1016/j.cels.2021.05.017, https://linkinghub.elsevier.com/retrieve/pii/S2405471221002039.

[5] J. D. Bloom, J. J. Silberg, C. O. Wilke, D. A. Drummond, C. Adami, and F. H. Arnold, Thermodynamic prediction of protein neutrality, Proceedings of the National Academy of Sciences, 102 (2005), pp. 606–611.

[6] D. H. Bryant, A. Bashir, S. Sinai, N. K. Jain, P. J. Ogden, P. F. Riley, G. M. Church, L. J. Colwell, and E. D. Kelsic, Deep diversification of an AAV capsid protein by machine learning, Nature Biotechnology, 39 (2021), pp. 691–696, 10.1038/s41587-020-00793-4, https://www.nature.com/articles/s41587-020-00793-4.

[7] B. Charlier, J. Feydy, J. A. Glaunés, F.-D. Collin, and G. Durif, Kernel operations on the GPU, with autodiff, without memory overflows, Journal of Machine Learning Research, 22 (2021), pp. 1–6.

[8] W.-c. Chen, J. Zhou, J. M. Sheltzer, J. B. Kinney, and D. M. Mccandlish, Field-theoretic density estimation for biological sequence space with applications to 5 splice site diversity and aneuploidy in cancer, Proc. Natl. Acad. Sci. USA, (2021), 10.1073/pnas.2025782118.

[9] J. N. Cobb, G. DeClerck, A. Greenberg, R. Clark, and S. McCouch, Next-generation phenotyping: Requirements and strategies for enhancing our understanding of genotype–phenotype relationships and its relevance to crop improvement, Theoretical and Applied Genetics, 126 (2013), pp. 867–887, 10.1007/s00122-013-2066-0, http://link.springer.com/10.1007/s00122-013-2066-0.

[10] K. Crona, A. Gavryushkin, D. Greene, and N. Beerenwinkel, Inferring genetic interactions from comparative fitness data, eLife, 6 (2017), p. e28629, 10.7554/eLife.28629, https://elifesciences.org/articles/28629.

[11] A. Currin, N. Swainston, P. J. Day, and D. B. Kell, Synthetic biology for the directed evolution of protein biocatalysts: Navigating sequence space intelligently, Chemical Society Reviews, 44 (2015), pp. 1172–1239, 10.1039/C4CS00351A, https://pubs.rsc.org/cs/article/44/5/1172-1239/434816.

[12] K. Dasari, J. A. Somarelli, S. Kumar, and J. P. Townsend, The somatic molecular evolution of cancer: Mutation, selection, and epistasis, Progress in Biophysics and Molecular Biology, 165 (2021), pp. 56–65, 10.1016/j.pbiomolbio.2021.08.003, https://linkinghub.elsevier.com/retrieve/pii/S0079610721000961.

[13] J. Domingo, P. Baeza-Centurion, and B. Lehner, The Causes and Consequences of Genetic Interactions (Epistasis), Annual Review of Genomics and Human Genetics, 20 (2019), pp. 433– 460, 10.1146/annurev-genom-083118-014857, https://www.annualreviews.org/doi/10.1146/annurev-genom-083118-014857.

[14] A. Escobedo, G. Voigt, A. J. Faure, and B. Lehner, Genetics, energetics and allostery during a billion years of hydrophobic protein core evolution, May 2024, 10.1101/2024.05.11.593672, http://biorxiv.org/lookup/doi/10.1101/2024.05.11.593672 (accessed 2024-06-09).

[15] A. J. Faure and B. Lehner, Mochi: neural networks to fit interpretable models and quantify energies, energetic couplings, epistasis, and allostery from deep mutational scanning data, Genome Biology, 25 (2024), p. 303, https://doi.org/10.1186/s13059-024-03444-y, 10.1186/s13059-024-03444-y (accessed 2025-03-09).

[16] L. Ferretti, B. Schmiegelt, D. Weinreich, A. Yamauchi, Y. Kobayashi, F. Tajima, and G. Achaz, Measuring epistasis in fitness landscapes: The correlation of fitness effects of mutations, Journal of Theoretical Biology, 396 (2016), pp. 132–143, 10.1016/j.jtbi.2016.01.037.

[17] R. A. Fisher, The genetical theory of natural selection, Clarendon Press, Oxford, 1930, 10.5962/bhl.title.27468, https://www.biodiversitylibrary.org/bibliography/27468.

[18] D. M. Fowler and S. Fields, Deep mutational scanning: A new style of protein science, Nature Methods, 11 (2014), pp. 801–807, 10.1038/nmeth.3027, https://www.nature.com/articles/nmeth.3027.

[19] C. R. Freschlin, S. A. Fahlberg, P. Heinzelman, and P. A. Romero, Neural network extrapolation to distant regions of the protein fitness landscape, Nature Communications, 15 (2024), p. 6405.

[20] J. Gardner, G. Pleiss, K. Q. Weinberger, D. Bindel, and A. G. Wilson, Gpytorch: Blackbox matrix-matrix gaussian process inference with gpu acceleration, Advances in neural information processing systems, 31 (2018).

[21] S. Gelman, S. A. Fahlberg, P. Heinzelman, P. A. Romero, and A. Gitter, Neural networks to learn protein sequence–function relationships from deep mutational scanning data, Proceedings of the National Academy of Sciences, 118 (2021), p. e2104878118, 10.1073/pnas.2104878118, https://pnas.org/doi/full/10.1073/pnas.2104878118.

[22] M. Ghafari, A. C. Cabrera, A. Lage-Castellanos, G. Achaz, J. Krug, and L. Ferretti, Epistatic strength, modularity, and locus heterogeneity shape the number of local optima in fitness landscapes, (2026), 10.48550/arXiv.2605.03046, http://arxiv.org/abs/2605.03046 (accessed 2026-11-06). arXiv:2605.03046 [q-bio.PE].

[23] L. Gonzalez Somermeyer, A. Fleiss, A. S. Mishin, N. G. Bozhanova, A. A. Igolkina, J. Meiler, M.-E. Alaball Pujol, E. V. Putintseva, K. S. Sarkisyan, and F. A. Kondrashov, Heterogeneity of the GFP fitness landscape and data-driven protein design, eLife, 11 (2022), p. e75842, 10.7554/eLife.75842, https://elifesciences.org/articles/75842.

[24] P. Groth, M. Kerrn, L. Olsen, J. Salomon, and W. Boomsma, Kermut: Composite kernel regression for protein variant effects, in Advances in Neural Information Processing Systems 37, Vancouver, BC, Canada, 2024, Neural Information Processing Systems Foundation, Inc. (NeurIPS), pp. 29514–29565, 10.52202/079017-0929, http://www.proceedings.com/079017-0929.html.

[25] S. Herrera-Alvarez, J. E. J. Patton, and J. W. Thornton, The structure of an ancient genotype– phenotype map shaped the functional evolution of a protein family, Nature Ecology & Evolution, 9 (2025), pp. 1656–1669, 10.1038/s41559-025-02777-6, https://www.nature.com/articles/s41559-025-02777-6.

[26] T. Hinkley, J. Martins, C. Chappey, M. Haddad, E. Stawiski, J. M. Whitcomb, C. J. Petropoulos, and S. Bonhoeffer, A systems analysis of mutational effects in hiv-1 protease and reverse transcriptase, Nature genetics, 43 (2011), pp. 487–489.

[27] T. Hoffbauer and B. Strodel, TransMEP: Transfer learning on large protein language models to predict mutation effects of proteins from a small known dataset, Jan. 2024, 10.1101/2024.01.12.575432, http://biorxiv.org/lookup/doi/10.1101/2024.01.12.575432.

[28] M. K. Howard and W. Coyote-Maestas, Mechanistic Mutational Scanning to Uncover the Secret Life of Proteins, Annual Review of Biomedical Data Science, (2026), 10.1146/annurev-biodatasci-092624-104436, https://www.annualreviews.org/content/journals/10.1146/annurev-biodatasci-092624-104436.

[29] C. Hsu, H. Nisonoff, C. Fannjiang, and J. Listgarten, Learning protein fitness models from evolutionary and assay-labeled data, Nature Biotechnology, 40 (2022), pp. 1114–1122, 10.1038/s41587-021-01146-5, https://www.nature.com/articles/s41587-021-01146-5.

[30] M. Huot, D. Wang, J. Liu, and E. I. Shakhnovich, Predicting high-fitness viral protein variants with Bayesian active learning and biophysics, Proceedings of the National Academy of Sciences, 122 (2025), p. e2503742122, 10.1073/pnas.2503742122, https://pnas.org/doi/10.1073/pnas.2503742122.

[31] S. Hwang, B. Schmiegelt, L. Ferretti, and J. Krug, Universality Classes of Interaction Structures for NK Fitness Landscapes, Journal of Statistical Physics, 172 (2018), pp. 226–278, 10.1007/s10955-018-1979-z, https://doi.org/10.1007/s10955-018-1979-z. arXiv: 1708.06556 Publisher: Springer US.

[32] A. S. Jalal, N. T. Tran, C. E. Stevenson, E. W. Chan, R. Lo, X. Tan, A. Noy, D. M. Lawson, and T. B. Le, Diversification of DNA-Binding Specificity by Permissive and Specificity-Switching Mutations in the ParB/Noc Protein Family, Cell Reports, 32 (2020), p. 107928,10.1016/j.celrep.2020.107928, https://linkinghub.elsevier.com/retrieve/pii/S2211124720309098 (accessed 2023-10-30).

[33] M. Jankowiak, Y. Ordabayev, R. Tuwani, H. N. Ward, H. Nisonoff, J. M. Mcfarland, and G. Grigoryan, Flexible kernels for protein property prediction, in Proceedings of the 43rd International Conference on Machine Learning, T. Zhang, M. Dudik, M. Jaggi, A. Agarwal, S. Li, D. Schuurmans, J. Zhu, F. Berkenkamp, H. Dong, and A. Bietti, eds., vol. 306 of Proceedings of Machine Learning Research, PMLR, 06–11 Jul 2026, pp. 50779–50829, https://proceedings.mlr.press/v306/jankowiak26a.html.

[34] K. E. Johnston, P. J. Almhjell, E. J. Watkins-Dulaney, G. Liu, N. J. Porter, J. Yang, and F. H. Arnold, A combinatorially complete epistatic fitness landscape in an enzyme active site, Proceedings of the National Academy of Sciences of the United States of America, 121 (2024), p. e2400439121, 10.1073/pnas.2400439121.

[35] S. A. Kauffman and E. D. Weinberger, The NK model of rugged fitness landscapes and its application to maturation of the immune response, Journal of Theoretical Biology, 141 (1989), pp. 211–245, 10.1016/S0022-5193(89)80019-0, https://doi.org/10.1016/S0022-5193(89)80019-0.

[36] H. Kemble, P. Nghe, and O. Tenaillon, Recent insights into the genotype–phenotype relationship from massively parallel genetic assays, Evolutionary Applications, 12 (2019), pp. 1721–1742, 10.1111/eva.12846, https://onlinelibrary.wiley.com/doi/10.1111/eva.12846.

[37] J. F. C. Kingman, A simple model for the balance between selection and mutation, Journal of Applied Probability, 15 (1978), pp. 1–12, 10.2307/3213231.

[38] J. B. Kinney and D. M. McCandlish, Massively Parallel Assays and Quantitative Sequence– Function Relationships, Annual Review of Genomics and Human Genetics, 20 (2019), pp. 99–127, 10.1146/annurev-genom-083118-014845, https://www.annualreviews.org/doi/10.1146/annurev-genom-083118-014845.

[39] S. Kryazhimskiy, D. P. Rice, E. R. Jerison, and M. M. Desai, Global epistasis makes adaptation predictable despite sequence-level stochasticity, Science, 344 (2014), pp. 1519–1522.

[40] M. Lässig, V. Mustonen, and A. Nourmohammad, Steering and controlling evolution — from bioengineering to fighting pathogens, Nature Reviews Genetics, 24 (2023), pp. 851–867,10.1038/s41576-023-00623-8, https://www.nature.com/articles/s41576-023-00623-8.

[41] Z. Lin, H. Akin, R. Rao, B. Hie, Z. Zhu, W. Lu, N. Smetanin, R. Verkuil, O. Kabeli,Y. Shmueli, A. Dos Santos Costa, M. Fazel-Zarandi, T. Sercu, S. Candido, and A. Rives, Evolutionary-scale prediction of atomic-level protein structure with a language model, Science, 379 (2023), pp. 1123–1130, 10.1126/science.ade2574, https://www.science.org/doi/10.1126/science.ade2574.

[42] T. L. V. Lite, R. A. Grant, I. Nocedal, M. L. Littlehale, M. S. Guo, and M. T. Laub, Uncovering the basis of protein-protein interaction specificity with a combinatorially complete library, eLife, 9 (2020), pp. 1–57, 10.7554/eLife.60924.

[43] Y. Luo, G. Jiang, T. Yu, Y. Liu, L. Vo, H. Ding, Y. Su, W. W. Qian, H. Zhao, and J. Peng, ECNet is an evolutionary context-integrated deep learning framework for protein engineering, Nature Communications, 12 (2021), p. 5743, 10.1038/s41467-021-25976-8, https://www.nature.com/articles/s41467-021-25976-8.

[44] C. Martí-Gómez, J. Zhou, W.-C. Chen, A. Stoltzfus, J. B. Kinney, and D. M. McCandlish, Inference and Visualization of Complex Genotype–Phenotype Maps, Molecular Biology and Evolution, 43 (2026), p. msag023, 10.1093/molbev/msag023, https://academic.oup.com/mbe/article/doi/10.1093/molbev/msag023/8456298.

[45] C. Martí-Gómez and D. M. McCandlish, Inference of fitness landscapes with heterogeneous patterns of epistasis across sites, June 2026, 10.64898/2026.06.25.734428, https://www.biorxiv.org/content/10.64898/2026.06.25.734428v1 (accessed 2026-09-11). ISSN: 2692-8205 Pages: 2026.06.25.734428 Section: New Results.

[46] D. M. McCandlish, Visualizing Fitness Landscapes, Evolution, 65 (2011), pp. 1544–1558, 10.1111/j.1558-5646.2011.01236.x, https://academic.oup.com/evolut/article/65/6/1544/6854390.

[47] A. Moulana, T. Dupic, A. M. Phillips, J. Chang, A. A. Roffler, A. J. Greaney, T. N. Starr, J. D. Bloom, and M. M. Desai, The landscape of antibody binding affinity in SARS-CoV-2 Omicron BA.1 evolution, eLife, 12 (2023), p. e83442, 10.7554/eLife.83442, https://elifesciences.org/articles/83442.

[48] J. Neidhart, I. G. Szendro, and J. Krug, Exact results for amplitude spectra of fitness landscapes, Journal of Theoretical Biology, 332 (2013), pp. 218–227, 10.1016/j.jtbi.2013.05.002, http://dx.doi.org/10.1016/j.jtbi.2013.05.002. arXiv: 1301.1923 Publisher: Elsevier.

[49] J. Otwinowski, Biophysical inference of epistasis and the effects of mutations on protein stability and function, Molecular Biology and Evolution, 35 (2018), pp. 2345–2354, 10.1093/molbev/msy141. arXiv: 1802.08744.

[50] J. Otwinowski, D. M. McCandlish, and J. B. Plotkin, Inferring the shape of global epistasis, Proceedings of the National Academy of Sciences, 115 (2018), 10.1073/pnas.1804015115, https://pnas.org/doi/full/10.1073/pnas.1804015115.

[51] R. Paul, A. Krishnan, and A. Dasgupta, Omics, CRISPR, and Artificial Intelligence: The Triad Revolutionizing Bioprocess Engineering, in Industrial Microbiology and Biotechnology, P. Verma, ed., Springer Nature Singapore, Singapore, 2026, pp. 57–84, 10.1007/978-981-95-5378-5_3, https://link.springer.com/10.1007/978-981-95-5378-5_3.

[52] S. Petti, C. Martí-Gómez, J. B. Kinney, J. Zhou, and D. M. McCandlish, On learning functions over biological sequence space: relating gaussian process priors, regularization, and gauge fixing, Journal of Mathematical Biology, 92 (2026), p. 78.

[53] S. Petti, G. Reddy, and M. M. Desai, Inferring sparse structure in genotype–phenotype maps, GENETICS, 225 (2023), p. iyad127, 10.1093/genetics/iyad127, https://academic.oup.com/genetics/article/doi/10.1093/genetics/iyad127/7223405.

[54] A. M. Phillips, K. R. Lawrence, A. Moulana, T. Dupic, J. Chang, M. S. Johnson, I. Cvijovic, T. Mora, A. M. Walczak, and M. M. Desai, Binding affinity landscapes constrain the evolution of broadly neutralizing anti-influenza antibodies, Elife, 10 (2021), p. e71393.

[55] P. C. Phillips, Epistasis — the essential role of gene interactions in the structure and evolution of genetic systems, Nature Reviews Genetics, 9 (2008), pp. 855–867, 10.1038/nrg2452, https://www.nature.com/articles/nrg2452.

[56] J. C. Pinheiro and D. M. Bates, Unconstrained parametrizations for variance-covariance matrices, Statistics and Computing, 6 (1996), pp. 289–296, 10.1007/BF00140873.

[57] F. J. Poelwijk, M. Socolich, and R. Ranganathan, Learning the pattern of epistasis linking genotype and phenotype in a protein, Nature Communications, 10 (2019), p. 4213, 10.1038/s41467-019-12130-8.

[58] A. Posfai, D. M. McCandlish, and J. B. Kinney, Symmetry, gauge freedoms, and the interpretability of sequence-function relationships, Physical Review Research, 7 (2025), p. 023005.

[59] A. Posfai, J. Zhou, D. M. McCandlish, and J. B. Kinney, Gauge fixing for sequence-function relationships, PLOS Computational Biology, 21 (2025), p. e1012818, 10.1371/journal.pcbi.1012818, https://dx.plos.org/10.1371/journal.pcbi.1012818.

[60] C. E. Rasmussen and C. K. I. Williams, Gaussian Processes for Machine Learning, Adaptive Computation and Machine Learning, MIT Press, Cambridge, Mass., 3. print ed., 2008.

[61] G. Reddy and M. M. Desai, Global epistasis emerges from a generic model of a complex trait, eLife, 10 (2021), p. e64740, 10.7554/eLife.64740, https://elifesciences.org/articles/64740.

[62] N. J. Rollins, K. P. Brock, F. J. Poelwijk, M. A. Stiffler, N. P. Gauthier, C. Sander, and D. S. Marks, Inferring protein 3d structure from deep mutation scans, Nature genetics, 51 (2019), pp. 1170–1176.

[63] P. A. Romero, A. Krause, and F. H. Arnold, Navigating the protein fitness landscape with Gaussian processes, Proceedings of the National Academy of Sciences, 110 (2013), 10.1073/pnas.1215251110, https://pnas.org/doi/full/10.1073/pnas.1215251110.

[64] T. B. Sackton and D. L. Hartl, Genotypic Context and Epistasis in Individuals and Populations, Cell, 166 (2016), pp. 279–287, 10.1016/j.cell.2016.06.047, https://linkinghub.elsevier.com/retrieve/pii/S0092867416308558.

[65] K. S. Sarkisyan, D. A. Bolotin, M. V. Meer, D. R. Usmanova, A. S. Mishin, G. V. Sharonov, D. N. Ivankov, N. G. Bozhanova, M. S. Baranov, O. Soylemez, N. S. Bogatyreva,P. K. Vlasov, E. S. Egorov, M. D. Logacheva, A. S. Kondrashov, D. M. Chudakov,E. V. Putintseva, I. Z. Mamedov, D. S. Tawfik, K. A. Lukyanov, and F. A. Kondrashov, Local fitness landscape of the green fluorescent protein, Nature, 533 (2016), pp. 397–401, 10.1038/nature17995, https://www.nature.com/articles/nature17995.

[66] J. M. Schmiedel and B. Lehner, Determining protein structures using deep mutagenesis, Nature genetics, 51 (2019), pp. 1177–1186.

[67] P. Sethi and J. Zhou, An interpretable neural network unveils higher-order epistasis in large protein sequence-function relationships, Mar. 2026, 10.7554/eLife.108005.2, https://elifesciences.org/reviewed-preprints/108005v2.

[68] V. W. C. Soo, J. B. Swadling, A. J. Faure, and T. Warnecke, Fitness landscape of a dynamic RNA structure, PLOS Genetics, 17 (2021), p. e1009353, 10.1371/journal.pgen.1009353, https://dx.plos.org/10.1371/journal.pgen.1009353.

[69] P. F. Stadler and R. Happel, Random field models for fitness landscapes, Journal of Mathematical Biology, 38 (1999), pp. 435–478, 10.1007/s002850050156.

[70] T. N. Starr and J. W. Thornton, Epistasis in protein evolution, Protein Science, 25 (2016), pp. 1204– 1218, 10.1002/pro.2897, https://onlinelibrary.wiley.com/doi/10.1002/pro.2897.

[71] A. Tareen, M. Kooshkbaghi, A. Posfai, W. T. Ireland, D. M. McCandlish, and J. B. Kinney, MAVE-NN: Learning genotype-phenotype maps from multiplex assays of variant effect, Genome Biology, 23 (2022), p. 98, 10.1186/s13059-022-02661-7, https://genomebiology.biomedcentral.com/articles/10.1186/s13059-022-02661-7.

[72] O. Tenaillon, The utility of fisher’s geometric model in evolutionary genetics, Annual review of ecology, evolution, and systematics, 45 (2014), pp. 179–201.

[73] P. D. Tonner, A. Pressman, and D. Ross, Interpretable modeling of genotype–phenotype landscapes with state-of-the-art predictive power, Proceedings of the National Academy of Sciences, 119 (2022), p. e2114021119, 10.1073/pnas.2114021119, https://pnas.org/doi/full/10.1073/pnas.2114021119.

[74] E. D. Vaishnav, C. G. de Boer, J. Molinet, M. Yassour, L. Fan, X. Adiconis, D. A. Thompson,J. Z. Levin, F. A. Cubillos, and A. Regev, The evolution, evolvability and engineering of gene regulatory DNA, Nature, (2022), 10.1038/s41586-022-04506-6. Publisher: Springer US.

[75] F. Vivarelli and C. Williams, Discovering hidden features with gaussian processes regression, in Advances in Neural Information Processing Systems, M. Kearns, S. Solla, and D. Cohn, eds., vol. 11, MIT Press, 1998, https://proceedings.neurips.cc/paperfiles/paper/1998/file/ed4227734ed75d343320b6a5fd16ce57-Paper.pdf.

[76] K. Wang, G. Pleiss, J. Gardner, S. Tyree, K. Q. Weinberger, and A. G. Wilson, Exact Gaussian Processes on a Million Data Points, NeurIPS, (2019).

[77] R. B. Waring, P. Towner, S. J. Minter, and R. W. Davies, Splice-site selection by a self-splicing RNA of Tetrahymena, Nature, 321 (1986), pp. 133–139, 10.1038/321133a0, https://www.nature.com/articles/321133a0 (accessed 2026-11-14). Publisher: Nature Publishing Group.

[78] D. M. Weinreich, Y. Lan, J. Jaffe, and R. B. Heckendorn, The Influence of Higher-Order Epistasis on Biological Fitness Landscape Topography, Journal of Statistical Physics, 172 (2018), pp. 208–225, 10.1007/s10955-018-1975-3, http://link.springer.com/10.1007/s10955-018-1975-3.

[79] A. G. Wilson, Z. Hu, R. Salakhutdinov, and E. P. Xing, Deep Kernel Learning, in Proceedings of the 19th International Conference on Artificial Intelligence and Statistics, A. Gretton and C. C. Robert, eds., vol. 51 of Proceedings of Machine Learning Research, Cadiz, Spain, May 2016, PMLR, pp. 370–378, https://proceedings.mlr.press/v51/wilson16.html.

[80] N. C. Wu, L. Dai, C. A. Olson, J. O. Lloyd-Smith, and R. Sun, Adaptation in protein fitness landscapes is facilitated by indirect paths, eLife, 5 (2016), p. e16965, 10.7554/eLife.16965, https://elifesciences.org/articles/16965.

[81] N. Wycoff, M. Binois, and S. M. Wild, Sequential learning of active subspaces, Journal of Computational and Graphical Statistics, 30 (2021), pp. 1224–1237, 10.1080/10618600.2021.1874962, https://doi.org/10.1080/10618600.2021.1874962, https://arxiv.org/abs/https://doi.org/10.1080/10618600.2021.1874962

[82] K. K. Yang, Z. Wu, and F. H. Arnold, Machine-learning-guided directed evolution for protein engineering, Nature methods, 16 (2019), pp. 687–694.

[83] K. K. Yang, Z. Wu, C. N. Bedbrook, and F. H. Arnold, Learned protein embeddings for machine learning, Bioinformatics, 34 (2018), pp. 2642–2648, 10.1093/bioinformatics/bty178, https://doi.org/10.1093/bioinformatics/bty178.

[84] W. Yang, H. Feng, X. Zhang, J. Zhang, J. H. Doonan, W. D. Batchelor, L. Xiong, and J. Yan, Crop Phenomics and High-Throughput Phenotyping: Past Decades, Current Challenges, and Future Perspectives, Molecular Plant, 13 (2020), pp. 187–214, 10.1016/j.molp.2020.01.008, https://linkinghub.elsevier.com/retrieve/pii/S1674205220300083.

[85] T. Zarin and B. Lehner, A complete map of specificity encoding for a partially fuzzy protein interaction, Apr. 2024, 10.1101/2024.04.25.591103, http://biorxiv.org/lookup/doi/10.1101/2024.04.25.591103 (accessed 2024-06-09).

[86] J. Zhou, C. Martí-Gómez, S. Petti, and D. M. McCandlish, Learning sequence-function relationships with scalable, interpretable Gaussian processes, Nov. 2025, 10.7554/eLife.108964.1, https://elifesciences.org/reviewed-preprints/108964v1.

[87] J. Zhou and D. M. McCandlish, Minimum epistasis interpolation for sequence-function relationships, Nature Communications, 11 (2020), p. 1782, 10.1038/s41467-020-15512-5, https://www.nature.com/articles/s41467-020-15512-5.

[88] J. Zhou, M. S. Wong, W.-C. Chen, A. R. Krainer, J. B. Kinney, and D. M. McCandlish, Higher-order epistasis and phenotypic prediction, Proceedings of the National Academy of Sciences, 119 (2022), p. e2204233119, 10.1073/pnas.2204233119, https://pnas.org/doi/full/10.1073/pnas.2204233119.

