## Supplementary Materials for "Embedding kernels for sequence-function relationships ^*^"

### SM1. Supplementary Information.

**SM1.1. FA kernel and its relationship to automatic relevance determination kernels and general product kernels.** This section provides more detail for Section 3 of the main text. We show that automatic relevance determination kernels (ARD) and a subclass of general product kernels (GPKs) arise as subclasses of FA kernels under restrictions on the embedding matrix $A$. We will describe these models with increasing flexibility.

When the FA kernel's embedding matrix $A$ is restricted to $A = cI$ for some constant $c$, it is an RBF kernel (an ARD kernel restricted to learning the same embedding scale for each allele-position pair). This corresponds to the Geometric kernel restricted to positive correlations:

$$k(s,s') = \exp\left(-||cIx(s) - cIx(s')||_2^2\right) = \exp\left(-2c^2 \cdot d_H(s,s')\right) = (1-\delta)^{d_H(s,s')} \tag{SM1.1}$$

where $d_H$ is the Hamming distance and $(1-\delta) = \exp(-2c^2)$.

When the FA kernel's embedding matrix $A$ is restricted to a diagonal matrix, whose diagonal sub-matrices are $c_p I_\alpha$ for each position $p$, we have an ARD kernel that is a prior on the Connectedness model [SM16]. This corresponds to our Connectedness kernel restricted to positive correlations [SM24]:

$$\begin{aligned} k(s,s') &= \sigma^2 \exp\left(-\sum_p ||c_p I_\alpha x(s_p) - c_p I_\alpha x(s'_p)||_2^2\right) \\ &= \sigma^2 \prod_p \exp\left(-c_p^2 \sum_a |\mathbb{1}_{s_p=a} - \mathbb{1}_{s'_p=a}|^2\right) \\ &= \sigma^2 \prod_p \exp\left(-2c_p^2 \cdot \mathbb{1}_{s_p \neq s'_p}\right) \\ &= \sigma^2 \prod_{p:s_p\neq s'_p} (1-\delta_p) \end{aligned} \tag{SM1.2}$$

where $(1-\delta_p) = \exp(-2c_p^2)$.

Next, we consider when $A$ is a diagonal matrix with $c_{p,a}$, an ARD kernel with distinct embedding scales for allele-position pairs $(a,p)$. This exactly coincides with the Jenga kernel,

† Co-first authors.

* Submitted to the editors November 15, 2026.

‡ Department of Mathematics, Tufts University.
§ Cold Spring Harbor Laboratory.

which generalizes the Connectedness model. As before we have:

$$
\begin{aligned}
k(s,s') &= \sigma^2 \prod_p \exp\left(-\sum_a |c_{p,a}\mathbb{1}_{s_p=a} - c_{p,a}\mathbb{1}_{s'_p=a}|^2\right) \\
&= \sigma^2 \prod_p \exp\left(-(c^2_{p,s_p} + c^2_{p,s'_p}) \cdot \mathbb{1}_{s_p\neq s'_p}\right) \\
&= \sigma^2 \prod_{p,s_p\neq s'_p} (1-\delta_p^{s_p})(1-\delta_p^{s'_p})
\end{aligned} \tag{SM1.3}
$$

where $(1-\delta_p^{s_p}) = \exp\left(-c^2_{p,s_p}\right)$ and $(1-\delta_p^{s'_p}) = \exp\left(-c^2_{p,s'_p}\right)$.

Each of these kernels is a type of general product kernel (GPK). We will show that the subset of GPKs that admit a site-specific Gaussian euclidean embedding are FA kernels with a block-diagonal $A$ matrix, with each $\alpha$-block being $A_p$ for position $p$ where $\alpha$ is the alphabet size. Now, rather than having an axis in the embedding space for each allele-position combination, we have $\alpha$ axes per position to account for every possible allele. This gives us:

$$
\begin{aligned}
k(s,s') &= \sigma^2 \prod_p \exp\left(-||A_p x(s_p) - A_p x(s'_p)||_2^2\right) \\
&= \sigma^2 \prod_p \exp\left(-||A_p^{s_p} - A_p^{s'_p}||_2^2\right) \\
&= \sigma^2 \prod_p (1-\delta_p^{s_p,s'_p})
\end{aligned} \tag{SM1.4}
$$

where $A_p^{s_p}$ is a vector of length $\alpha$, the corresponding $s_p$th column of $A_p$ block.

**SM1.2. Gauge freedoms in the additive embeddings.** In the main text, we discussed the gauge freedom arising from using the one-hot sequence encoding to parametrize the additive embeddings [SM13] and the fact that the kernel only depends on the distances between sequences in those embeddings. These freedoms are apparent from rewriting the FA kernel of Equation 3.1 in the main text as

$$
k(s,s') = \sigma^2 \exp\left(-(x(s)-x(s'))^T M (x(s)-x(s'))\right), \tag{SM1.5}
$$

where $M = A^T A$. There are many $A$ matrices resulting in the same $M$ matrix, corresponding to orthogonal rotations of $A$, that encode the same kernel. However, the dependent nature of the one-hot encoding allows translations of blocks of $A$ that also preserve the kernel. Here, we provide an example of the translation invariance of $A$ and how we address it.

We can translate the entire matrix $A$ or any combination of $\alpha$-block columns or rows of $A$. Without loss of generality, let $A_{i,j}$ be an $\alpha$-block row of $A$ in position $(i,j)$. Then we define the translation $\Delta_c$ to be $c\mathbf{1}_\alpha^T$ at $(i,j)$ and zero everywhere else. Let $A' = A + \Delta_c$. Clearly, $A' = A$ at every block except for $(i,j)$. Therefore, we only need to consider $A'_{i,j} = A_{i,j} + c\mathbf{1}_\alpha^T$. We see that when applied to differences of one-hot encoded sequences, we get the same result.

Let $x_j(s)$ be the $\alpha$-dimensional block of $x(s)$ associated to position $j$. Then:

$$(x_j(s) - x_j(s'))^T A'^{\,T}_{i,j} A'_{i,j}(x_j(s) - x_j(s')) \tag{SM1.6}$$
$$= (x_j(s) - x_j(s'))^T (A_{i,j} + c\mathbf{1}^T_\alpha)^T (A_{i,j} + c\mathbf{1}^T_\alpha)(x_j(s) - x_j(s')) \tag{SM1.7}$$
$$= (x_j(s) - x_j(s'))^T (A^T_{i,j}A_{i,j} + c\mathbf{1}_\alpha A_{i,j} + cA^T_{i,j}\mathbf{1}^T_\alpha + c_2\mathbf{1}_\alpha\mathbf{1}^T_\alpha)(x_j(s) - x_j(s')) \tag{SM1.8}$$
$$= (x_j(s) - x_j(s'))^T A^T_{i,j}A_{i,j}(x_j(s) - x_j(s')), \tag{SM1.9}$$

where we used the fact that $\mathbf{1}^T_\alpha(x_j(s) - x_j(s')) = 0$ for any pair of sequences $s$ and $s'$.

Equation SM1.6 shows that adding the same constant to all allele coefficients within a position block row of $A$ does not change the kernel. We remove this freedom by requiring these coefficients to sum to zero within each block, defined as a zero-sum gauge [SM13]. To achieve this, we apply $Q$ as a block diagonal matrix with $\ell$ blocks being $I_\alpha - \frac{1}{\alpha}\mathbf{1}_\alpha\mathbf{1}^T_\alpha$ described in Section 5 of the main text to $A$, as $AQ$. This subtracts the mean coefficient within each position block of every row of $A$. Since each position block of $x(s) - x(s')$ sums to zero, we have

$$Q(x(s) - x(s')) = x(s) - x(s'),$$

and therefore,

$$AQ(x(s) - x(s')) = A(x(s) - x(s')).$$

Thus, applying $Q$ removes the block-specific translation freedom while preserving all latent distances and hence the FA kernel.

**SM1.3. Decay factors for pairs of sites.** In this section, we derive Equation 3.4 from the main text by redefining the FA kernel in terms of its decay factors $\delta_p^{c_1,c_2}$ and $\delta_{pq^{c_1,c_2,c_3,c_4}}$, which describe the kernel by its single site and pairwise contributions to the overall covariance.

We can achieve this by writing the FA kernel from Equation SM1.5 by grouping $M$ into its $\alpha \times \alpha$ block matrices $M_{pq}$ for positions $p$ and $q$:

$$M = \begin{bmatrix} M_{11} & M_{12} & .. & M_{1\ell} \\ M_{21} & M_{22} & \dots & M_{2\ell} \\ \vdots & \ddots & & \vdots \\ M_{\ell 1} & M_{\ell 2} & \dots & M_{\ell\ell} \end{bmatrix}$$

This enables us to expand the quadratic form in the exponential as:

$$\begin{aligned}(x(s) - x(s'))^T M(x(s) - x(s')) &= \sum_{p,q}(x(s_p) - x(s'_p))^T M_{pq}(x(s_q) - x(s'_q)) \\ &= \sum_{p,q} x(s_p)^T M_{pq}x(s_q) + x(s'_p)^T M_{pq}x(s'_q) - x(s_p)^T M_{pq}x(s'_q) - x(s'_p)^T M_{pq}x(s_q).\end{aligned} \tag{SM1.10}$$

Define $M_{pq}^{s_p,s'_q} = x(s_p)^T M_{pq}x(s'_q)$ and the other terms accordingly. Then we can rewrite the kernel in the following way to extract the decay factors:

$$\begin{aligned} k(s,s') &= \sigma^2 \exp\left(-(x(s)-x(s'))^T M (x(s)-x(s'))\right) \\ &= \sigma^2 \prod_{\substack{p,q \\ s_p \neq s'_p \\ s_q \neq s'_q}} \exp\left(-(M_{pq}^{s_p,s_q} + M_{pq}^{s'_p,s'_q} - M_{pq}^{s_p,s'_q} - M_{pq}^{s'_p,s_q})\right) \end{aligned}$$

We use the symmetry of $M$ to combine contributions from $(p,q)$ and $(q,p)$ for $p \neq q$ so that we can separate this product into two cases: when $p = q$ and when $q < p$. By defining our factors $\delta_p^{s_p,s'_p} = \delta_{pp}^{s_p,s'_p,s_p,s'_p}$ when $p = q$ and more generally:

$$\delta_{pq}^{s_p,s'_p,s_q,s'_q} = 1 - \exp\left(-(2 - \mathbb{1}_{p=q})(M_{pq}^{s_p,s_q} + M_{pq}^{s'_p,s'_q} - M_{pq}^{s_p,s'_q} - M_{pq}^{s'_p,s_q})\right) \tag{SM1.11}$$

we can rewrite the equation using these factors:

$$\begin{aligned} k(s,s') &= \sigma^2 \prod_{\substack{p=q \\ s_p \neq s'_p}} (1 - \delta_p^{s_p,s'_p}) \prod_{\substack{q<p \\ s_p \neq s'_p \\ s_q \neq s'_q}} (1 - \delta_{pq}^{s_p,s'_p,s_q,s'_q}) \\ &= \sigma^2 \prod_{\substack{p \\ s_p \neq s'_p}} \left[ (1 - \delta_p^{s_p,s'_p}) \prod_{\substack{q<p \\ s_q \neq s'_q}} (1 - \delta_{pq}^{s_p,s'_p,s_q,s'_q}) \right] \end{aligned} \tag{SM1.12}$$

Now we have defined the FA kernel as a product of factors that depend on single site positions $\delta_p^{s_p,s'_p}$ and pairs of positions $\delta_{pq}^{s_p,s'_p,s_q,s'_q}$.

While these decay factors help interpret how individual mutations and their combinations affect the prior covariance between sequences, they cannot be chosen independently. Counting the nontrivial single-site factors and both relative orientations of each pair of mutations gives $\ell\binom{\alpha}{2} + 2\binom{\ell}{2}\binom{\alpha}{2}^2$ decay factors. For $\ell \geq 2$ and $\alpha \geq 2$, this exceeds the $\ell(\alpha-1) + \binom{\ell(\alpha-1)}{2}$ free parameters in the reduced, gauge-fixed $M$ matrix without a rank restriction. The decay factors therefore satisfy relationships inherited from their dependence on the same entries of $M$ (Equation SM1.11). They are further constrained by the requirement that $M$ be positive semi-definite. One such relationship follows by reversing one of two mutations at different positions $p \neq q$:

$$1 - \delta_{pq}^{c_1,c_2,c_3,c_4} = \exp\left[-2\left(M_{pq}^{c_1,c_3} + M_{pq}^{c_2,c_4} - M_{pq}^{c_1,c_4} - M_{pq}^{c_2,c_3}\right)\right] = \left(1 - \delta_{pq}^{c_1,c_2,c_4,c_3}\right)^{-1}. \tag{SM1.13}$$

Consequently,

$$\delta_{pq}^{c_1,c_2,c_4,c_3} = \frac{\delta_{pq}^{c_1,c_2,c_3,c_4}}{\delta_{pq}^{c_1,c_2,c_3,c_4} - 1}. \tag{SM1.14}$$

Thus, the two factors have opposite signs whenever they are nonzero: if $0 < \delta_{pq}^{c_1,c_2,c_3,c_4} < 1$, then $\delta_{pq}^{c_1,c_2,c_4,c_3} < 0$, and conversely.

**SM1.4. Predictability of mutational effects across genetic backgrounds.** In this section, we derive the expected predictability of mutational effects across genetic backgrounds separated by a particular mutation, and its generalization to backgrounds separated by an arbitrary set of mutations. This section provides additional details to our discussion of Section 3.2 in the main text.

Let $w_{p:c_1\to c_2}(s|f)$ be the effect of mutation $c_1 \to c_2$ at position $p$ in genetic background $s$ obtained by subtracting the function $f$ evaluated at sequence $s$ with allele $c_1$ compared to sequence $s$ with allele $c_2$ at position $p$ for an arbitrary sequence-function map $f$:

$$w_{p:c_1\to c_2}(s|f) = f(t_p^{c_2}(s)) - f(t_p^{c_1}(s)), \tag{SM1.15}$$

where $t_p^c(s) = s_1 \dots s_{p-1} c s_{p+1} \dots s_\ell$ is the sequence resulting from placing allele $c$ at position $p$ in sequence $s$.

Let $\mathcal{D}(s, s')$ be a function that returns the directed set of mutations at which any two sequences $s$ and $s'$ differ, where mutations are defined by their position and the pair of different alleles at sequences $s$ and $s'$, respectively:

$$\mathcal{D}(s, s') = \{(p, s_p, s'_p) \mid s_p \neq s'_p\}. \tag{SM1.16}$$

We define $\Gamma^D_{p:c_1\to c_2}(f)$ to be the second moment of the effects of mutation $c_1 \to c_2$ at position $p$ across genetic backgrounds $s$ and $s'$ that differ by an exact set of mutations $D$, not including position $p$ ($(p, s_p, s'_p) \notin D$):

$$\begin{aligned}
\Gamma^D_{p:c_1\to c_2}(f) &= \mathbb{E}_{s,s':\mathcal{D}(s,s')=D}\left[w_{p:c_1\to c_2}(s|f) w_{p:c_1\to c_2}(s'|f)\right] \\
&= \frac{1}{N_D} \sum_{s,s':\mathcal{D}(s,s')=D} \left[f(t_p^{c_2}(s)) - f(t_p^{c_1}(s))\right]\left[f(t_p^{c_2}(s')) - f(t_p^{c_1}(s'))\right] \\
&= \frac{1}{N_D} \sum_{s,s':\mathcal{D}(s,s')=D} [f(t_p^{c_2}(s)) f(t_p^{c_2}(s')) - f(t_p^{c_1}(s)) f(t_p^{c_2}(s')) \\
&\qquad - f(t_p^{c_2}(s)) f(t_p^{c_1}(s')) + f(t_p^{c_1}(s)) f(t_p^{c_1}(s'))],
\end{aligned} \tag{SM1.17}$$

where $N_D$ is the number of pairs of sequences differing at mutation set $D$. Next, we ask what is the expected value of this $\Gamma^D_{p:c_1\to c_2}(f)$ statistic under the prior defined by the FA kernel. Using the fact that $\mathbb{E}_{f\sim\mathcal{N}(0,K)}[f(s)f(s')] = k(s, s')$ since the mean of the prior is zero, we get:

$$\begin{aligned}
&\mathbb{E}_{f\sim\mathcal{N}(0,K)}\left[\Gamma^D_{p:c_1\to c_2}(f)\right] \\
&= \frac{1}{N_D} \sum_{s,s':\mathcal{D}(s,s')=D} \mathbb{E}_{f\sim\mathcal{N}(0,K)}[f(t_p^{c_2}(s)) f(t_p^{c_2}(s')) - f(t_p^{c_1}(s)) f(t_p^{c_2}(s')) \\
&\qquad - f(t_p^{c_2}(s)) f(t_p^{c_1}(s')) + f(t_p^{c_1}(s)) f(t_p^{c_1}(s'))] \\
&= \frac{1}{N_D} \sum_{s,s':\mathcal{D}(s,s')=D} k\left(t_p^{c_2}(s), t_p^{c_2}(s')\right) - k\left(t_p^{c_1}(s), t_p^{c_2}(s')\right) \\
&\qquad - k\left(t_p^{c_2}(s), t_p^{c_1}(s')\right) + k\left(t_p^{c_1}(s), t_p^{c_1}(s')\right) \\
&= 2k(D) - k(D \cup \{(p, c_1, c_2)\}) - k(D \cup \{(p, c_2, c_1)\}),
\end{aligned} \tag{SM1.18}$$

where $k(D)$ denotes the kernel function for any pair of sequences differing by the mutation set $D$. If we write these terms under the FA prior, we obtain

$$k(D) = \sigma^2 \prod_{(q,c_3,c_4)\in D} \left(1-\delta_q^{c_3,c_4}\right) \prod_{\substack{(q,c_3,c_4),(r,c_5,c_6)\in D \\ r<q}} \left(1-\delta_{qr}^{c_3,c_4,c_5,c_6}\right)$$
$$k\left(D\cup\{(p,c_1,c_2)\}\right) = k(D)\left(1-\delta_p^{c_1,c_2}\right)\prod_{(q,c_3,c_4)\in D}\left(1-\delta_{pq}^{c_1,c_2,c_3,c_4}\right)$$
$$k\left(D\cup\{(p,c_2,c_1)\}\right) = k(D)\left(1-\delta_p^{c_2,c_1}\right)\prod_{(q,c_3,c_4)\in D}\left(1-\delta_{pq}^{c_1,c_2,c_4,c_3}\right).$$

Then, we find that

$$\mathbb{E}_{f\sim\mathcal{N}(0,K)}\left[\Gamma^D_{p:c_1\to c_2}(f)\right] = k(D)C_{p:c_1\to c_2}(D), \tag{SM1.19}$$

where

$$C_{p:c_1\to c_2}(D) = 2-\left(1-\delta_p^{c_1,c_2}\right)\left[\prod_{(q,c_3,c_4)\in D}\left(1-\delta_{pq}^{c_1,c_2,c_3,c_4}\right)+\prod_{(q,c_3,c_4)\in D}\left(1-\delta_{pq}^{c_1,c_2,c_4,c_3}\right)\right]. \tag{SM1.20}$$

Using the reciprocal relationship above and $R_p(D)=\prod_{(q,c_3,c_4)\in D}\left(1-\delta_{pq}^{c_1,c_2,c_3,c_4}\right)$, we can write

$$C_{p:c_1\to c_2}(D) = 2-\left(1-\delta_p^{c_1,c_2}\right)\left[R_p(D)+R_p(D)^{-1}\right] \tag{SM1.21}$$
$$= 2-\left(1-\delta_p^{c_1,c_2}\right)\left[R_p(D)+R_p(D)^{-1}+2-2\right] \tag{SM1.22}$$
$$= 2-2\left(1-\delta_p^{c_1,c_2}\right)-\left(1-\delta_p^{c_1,c_2}\right)\frac{R_p(D)^2+1-2R_p(D)}{R_p(D)} \tag{SM1.23}$$
$$= 2\delta_p^{c_1,c_2}-\left(1-\delta_p^{c_1,c_2}\right)\frac{\left(1-R_p(D)\right)^2}{R_p(D)} \tag{SM1.24}$$
$$= 2\delta_p^{c_1,c_2}-\left(1-\delta_p^{c_1,c_2}\right)\frac{\left(1-\prod_{(q,c_3,c_4)\in D}\left(1-\delta_{pq}^{c_1,c_2,c_3,c_4}\right)\right)^2}{\prod_{(q,c_3,c_4)\in D}\left(1-\delta_{pq}^{c_1,c_2,c_3,c_4}\right)}. \tag{SM1.25}$$

Now let $\gamma^D_{p:c_1\to c_2}(f)$ be the normalized statistic:

$$\gamma^D_{p:c_1\to c_2}(f) = \frac{\mathbb{E}_{s,s':\mathcal{D}(s,s')=D}\left[w_{p:c_1\to c_2}(s|f)w_{p:c_1\to c_2}(s'|f)\right]}{\sqrt{\mathbb{E}_{s,s':\mathcal{D}(s,s')=D}\left[w_{p:c_1\to c_2}(s|f)^2\right]}\sqrt{\mathbb{E}_{s,s':\mathcal{D}(s,s')=D}\left[w_{p:c_1\to c_2}(s'|f)^2\right]}}, \tag{SM1.26}$$

Let us now consider the expected value of average squared mutational effect across each set of backgrounds that appear in the denominator:

$$\mathbb{E}_{f\sim\mathcal{N}(0,K)}\left[\mathbb{E}_{s,s':\mathcal{D}(s,s')=D}\left[w_{p:c_1\to c_2}(s|f)^2\right]\right] \tag{SM1.27}$$

$$= \frac{1}{N_D}\sum_{s,s':\mathcal{D}(s,s')=D}\mathbb{E}_{f\sim\mathcal{N}(0,K)}\left[\left(f(t_p^{c_2}(s)) - f(t_p^{c_1}(s))\right)^2\right] \tag{SM1.28}$$

$$= \frac{1}{N_D}\sum_{s,s':\mathcal{D}(s,s')=D}\mathbb{E}_{f\sim\mathcal{N}(0,K)}\left[f(t_p^{c_2}(s))^2 - 2f(t_p^{c_2}(s))f(t_p^{c_1}(s)) + f(t_p^{c_1}(s))^2\right] \tag{SM1.29}$$

$$= \frac{1}{N_D}\sum_{s,s':\mathcal{D}(s,s')=D}\left[k(t_p^{c_2}(s),t_p^{c_2}(s)) - 2k(t_p^{c_2}(s),t_p^{c_1}(s)) + k(t_p^{c_1}(s),t_p^{c_1}(s))\right] \tag{SM1.30}$$

$$= 2k(\emptyset) - 2k(\{(p,c_1,c_2)\}) = 2\sigma^2\left(1-(1-\delta_p^{c_1,c_2})\right) = 2\sigma^2\delta_p^{c_1,c_2}. \tag{SM1.31}$$

Note that the expected value of the average squared mutational effects of mutation $p : c_1 \to c_2$ is the same across backgrounds containing one set of alleles or another from the mutation set $D$, so that

$$\mathbb{E}_{f\sim\mathcal{N}(0,K)}\left[\mathbb{E}_{s,s':\mathcal{D}(s,s')=D}\left[w_{p:c_1\to c_2}(s|f)^2\right]\right] = \mathbb{E}_{f\sim\mathcal{N}(0,K)}\left[\mathbb{E}_{s,s':\mathcal{D}(s,s')=D}\left[w_{p:c_1\to c_2}(s'|f)^2\right]\right] \tag{SM1.32}$$

We approximate the expected normalized second-moment statistic by replacing the background averaged product and each background averaged squared effect in Equation SM1.26 with their respective expectations under the FA prior given by equations SM1.19 and SM1.27:

$$\mathbb{E}_{f\sim\mathcal{N}(0,K)}\left[\gamma^D_{p:c_1\to c_2}(f)\right] \approx \frac{k(D)C_{p:c_1\to c_2}(D)}{\sqrt{2\sigma^2\delta_p^{c_1,c_2}}\sqrt{2\sigma^2\delta_p^{c_1,c_2}}} = \frac{k(D)}{\sigma^2}\frac{C_{p:c_1\to c_2}(D)}{2\delta_p^{c_1,c_2}}. \tag{SM1.33}$$

This equation shows that the predictability of a given mutation depends not only on the sites at which the genetic backgrounds differ, but also on the specific decay factors associated with that mutation. However, if the decay factors for pairs of alleles involving a given mutation $c_1 \to c_2$ at position $p$ and any other mutation at which the two genetic backgrounds differ are set to zero ($\delta_{pq}^{c_1,c_2,c_3,c_4} = 0$ and $\delta_{pq}^{c_1,c_2,c_4,c_3} = 0$ for all $(q,c_3,c_4) \in D$), then the predictability of the effect of that particular mutation becomes independent of the specific mutation, and only depends on the set of mutations $D$ at which the genetic backgrounds $s$ and $s'$ differ.

For the specific case in which $D = \{(q,c_3,c_4)\}$, the term $C_{p:c_1\to c_2}(D)$ simplifies as follows:

$$\begin{aligned} C_{p:c_1\to c_2}(\{(q,c_3,c_4)\}) &= 2\delta_p^{c_1,c_2} - \left(1-\delta_p^{c_1,c_2}\right)\frac{\left(1-(1-\delta_{pq}^{c_1,c_2,c_3,c_4})\right)^2}{1-\delta_{pq}^{c_1,c_2,c_3,c_4}} \\ &= 2\delta_p^{c_1,c_2} - \left(1-\delta_p^{c_1,c_2}\right)\delta_{pq}^{c_1,c_2,c_3,c_4}\frac{\delta_{pq}^{c_1,c_2,c_3,c_4}}{1-\delta_{pq}^{c_1,c_2,c_3,c_4}}, \end{aligned}$$

allowing us to obtain an approximation to the expected value for the $\gamma^{(q,c_3,c_4)}_{p:c_1\to c_2}(f)$ statistic under the prior:

$$\mathbb{E}_{f\sim\mathcal{N}(0,K)}\left[\gamma^{(q,c_3,c_4)}_{p:c_1\to c_2}(f)\right] \approx \frac{\sigma^2(1-\delta_q^{c_3,c_4})}{\sigma^2}\frac{2\delta_p^{c_1,c_2}-(1-\delta_p^{c_1,c_2})\,\delta_{pq}^{c_1,c_2,c_3,c_4}\frac{\delta_{pq}^{c_1,c_2,c_3,c_4}}{1-\delta_{pq}^{c_1,c_2,c_3,c_4}}}{2\delta_p^{c_1,c_2}}$$
$$= (1-\delta_q^{c_3,c_4})\left[1-\frac{1}{2}\delta_{pq}^{c_1,c_2,c_3,c_4}\frac{(1-\delta_p^{c_1,c_2})}{\delta_p^{c_1,c_2}}\frac{\delta_{pq}^{c_1,c_2,c_3,c_4}}{1-\delta_{pq}^{c_1,c_2,c_3,c_4}}\right].$$

Given that $(1-\delta_p^{c_1,c_2})\geq 0$ and $(1-\delta_{pq}^{c_1,c_2,c_3,c_4})\geq 0$, the correction applied by the FA kernel can only be smaller than 1 and thus further decrease the predictability of mutational effects across backgrounds beyond what would be expected under a GPK prior.

This statistic is closely related to the previously proposed $\gamma_{(A_i,B_i)\to(A_j,B_j)}$ statistic [SM1], but differs in its treatment of mutation direction and normalization. The earlier definition averages over both directions of each focal mutation, including $c_1 \to c_2$ and $c_2 \to c_1$ with equal weight. Because their effects have opposite signs, this symmetry ensures that the mean mutational effect is zero, allowing the normalized second moment to coincide with a Pearson correlation. In contrast, our definition of $\gamma^D_{p:c_1\to c_2}(f)$ fixes the direction of the focal mutation, whose mean effect across backgrounds need not be zero. We therefore interpret this statistic as a normalized second moment rather than a Pearson correlation. Centering the effects within each landscape would introduce additional background-averaged terms, complicating the calculation of its expectation under the prior.

The definitions also differ in their normalization. The denominator in [SM1] pools the squared mutational effects across both sets of backgrounds, corresponding to the arithmetic mean of their respective mean squared effects. Equation SM1.26 instead uses the product of the root mean squared effects in the two sets of backgrounds, corresponding to the geometric mean of their mean squared effects. Our statistic is therefore equivalent to the cosine similarity between the two vectors of paired mutational effects (specifically, the vectors $u$ and $v$ indexed by pairs of sequences $(s,s')$ with $\mathcal{D}(s,s') = D$ in which $u_{(s,s')} = w_{p:c_1\to c_2}(s|f)$ and $v_{(s,s')} = w_{p:c_1\to c_2}(s'|f)$) and measures their alignment: it equals 1 when the vectors are positively proportional, $-1$ when they are negatively proportional, and 0 when they are orthogonal.

**SM1.5. Predictability of the effects of different mutations.** In this section, we derive the expected relationship in the effects of different mutations when they are introduced in the same genetic background. This provides further detail to our discussion of Section 3.2 in the main text.

As in the previous section, let $w_{p:c_1\to c_2}(s|f)$ be the effect of mutation $c_1 \to c_2$ at position $p$ in genetic background $s$ obtained by subtracting the function $f$ evaluated at sequence $s$ with allele $c_1$ compared to sequence $s$ with allele $c_2$ at position $p$:

$$w_{p:c_1\to c_2}(s|f) = f(t_p^{c_2}(s)) - f(t_p^{c_1}(s)), \tag{SM1.34}$$

where $t_p^c(s) = s_1 \dots s_{p-1} c s_{p+1} \dots s_\ell$ is the sequence resulting from placing allele $c$ at position $p$ in sequence $s$.

Let $\Gamma_{p:c_1\to c_2;\, q:c_3\to c_4}(f)$ be the second moment between the effect of mutation $c_1 \to c_2$ at position $p$, $w_{p:c_1\to c_2}(s|f)$, and the effect of mutation $c_3 \to c_4$ at position $q$, $w_{q:c_3\to c_4}(s|f)$, across all possible genetic backgrounds $s$ containing alleles $c_1$ and $c_3$ at positions $p$ and $q$, respectively:

$$\begin{aligned}
\Gamma_{p:c_1\to c_2;\,q:c_3\to c_4}(f) &= \mathbb{E}_{s:s_p=c_1,s_q=c_3}[w_{p:c_1\to c_2}(s|f)w_{q:c_3\to c_4}(s|f)] \\
&= \frac{1}{N}\sum_{s:s_p=c_1,s_q=c_3} w_{p:c_1\to c_2}(s|f)w_{q:c_3\to c_4}(s|f) \\
&= \frac{1}{N}\sum_{s:s_p=c_1,s_q=c_3} [f(t_p^{c_2}(s)) - f(s)][f(t_q^{c_4}(s)) - f(s)] \\
&= \frac{1}{N}\sum_{s:s_p=c_1,s_q=c_3} [f(t_p^{c_2}(s))f(t_q^{c_4}(s)) - f(s)f(t_q^{c_4}(s)) \\
&\quad - f(t_p^{c_2}(s))f(s) + f(s)f(s)],
\end{aligned} \tag{SM1.35}$$

where $N$ is the number of sequences satisfying $s_p = c_1$ and $s_q = c_3$, and $t_p^c(s)$ returns the sequence resulting from placing allele $c$ at position $p$ in sequence $s$. We then consider the expectation of this statistic under the prior to obtain

$$\begin{aligned}
&\mathbb{E}_{f\sim\mathcal{N}(0,K)}\left[\Gamma_{p:c_1\to c_2;\,q:c_3\to c_4}(f)\right] \\
&\quad= \frac{1}{N}\sum_{s:s_p=c_1,s_q=c_3} \mathbb{E}_{f\sim\mathcal{N}(0,K)}[f(t_p^{c_2}(s))f(t_q^{c_4}(s)) - f(s)f(t_q^{c_4}(s)) \\
&\quad - f(t_p^{c_2}(s))f(s) + f(s)f(s)] \\
&\quad= \frac{1}{N}\sum_{s:s_p=c_1,s_q=c_3} \Big[k\left(t_p^{c_2}(s), t_q^{c_4}(s)\right) - k\left(s, t_q^{c_4}(s)\right) \\
&\quad - k\left(s, t_p^{c_2}(s)\right) + k\left(s,s\right)\Big],
\end{aligned} \tag{SM1.36}$$

where we used the linearity of expectation and the fact that the expectation of the second moments corresponds exactly to the prior covariance between pairs of sequences.

Let us start by considering the case where the two mutations take place at the same position and are introduced in the same exact sequence ($p = q$, $c_3 = c_1$ and $c_4 \neq c_2$). Then the covariances only depend on the mutation-specific decay factors for those specific mutations, and not on decay factors for pairs of mutations. Thus, if we denote $c_4$ as $c_3$ for consistent numbering, we find that

$$\begin{aligned}
\mathbb{E}_{f\sim\mathcal{N}(0,K)}\left[\Gamma_{p:c_1\to c_2;\,p:c_1\to c_3}(f)\right] &= \sigma^2\left[1 - \left(1-\delta_p^{c_1,c_2}\right) - \left(1-\delta_p^{c_1,c_3}\right) + \left(1-\delta_p^{c_2,c_3}\right)\right] \\
&= \sigma^2\left[\delta_p^{c_1,c_2} + \delta_p^{c_1,c_3} - \delta_p^{c_2,c_3}\right].
\end{aligned} \tag{SM1.37}$$

We then approximate the expected normalized statistic $\gamma_{p:c_1\to c_2;\,p:c_1\to c_3}(f)$ by replacing the background-averaged product and each background-averaged squared effect with their expectations under the prior:

(SM1.38)

$$\begin{aligned}
&\mathbb{E}_{f\sim\mathcal{N}(0,K)}\left[\gamma_{p:c_1\to c_2;\,p:c_1\to c_3}(f)\right] \\
&\qquad\approx \frac{\mathbb{E}_{f\sim\mathcal{N}(0,K)}\left[\Gamma_{p:c_1\to c_2;\,p:c_1\to c_3}(f)\right]}{\sqrt{\mathbb{E}_{f\sim\mathcal{N}(0,K)}\left[\mathbb{E}_{s:s_p=c_1}[w_{p:c_1\to c_2}(s|f)^2]\right]\mathbb{E}_{f\sim\mathcal{N}(0,K)}\left[\mathbb{E}_{s:s_p=c_1}[w_{p:c_1\to c_3}(s|f)^2]\right]}} \\
&\qquad= \frac{\delta_p^{c_1,c_2}+\delta_p^{c_1,c_3}-\delta_p^{c_2,c_3}}{2\sqrt{\delta_p^{c_1,c_2}\delta_p^{c_1,c_3}}}.
\end{aligned}$$

Note that Equation SM1.38 is independent of any decay factor for pairs of mutations, showing that the FA kernel is as expressive as the GPK model in terms of the relationship between the effects of mutations at the same site. It is easy to see, for instance, that both models allow the effects of different mutations to be positively proportional to one another when $\delta_p^{c_2,c_3}=\delta_p^{c_1,c_2}+\delta_p^{c_1,c_3}-2\sqrt{\delta_p^{c_1,c_2}\delta_p^{c_1,c_3}}$.

Finally, we consider the case in which mutations take place at different positions ($p\neq q$) but in the same genetic background $s$, so that we only consider backgrounds where $s_p=c_1$ and $s_q=c_3$. In this case, by replacing the corresponding prior covariances in Equation SM1.36, we find that

(SM1.39)

$$\begin{aligned}
&\mathbb{E}_{f\sim\mathcal{N}(0,K)}\left[\Gamma_{p:c_1\to c_2;\,q:c_3\to c_4}(f)\right] \\
&\qquad= \frac{1}{N}\sum_{s:s_p=c_1,s_q=c_3} k\left(t_{pq}^{c_2,c_3}(s),t_{pq}^{c_1,c_4}(s)\right)-k\left(t_{pq}^{c_1,c_3}(s),t_{pq}^{c_1,c_4}(s)\right) \\
&\qquad\quad -k\left(t_{pq}^{c_1,c_3}(s),t_{pq}^{c_2,c_3}(s)\right)+k\left(t_{pq}^{c_1,c_3}(s),t_{pq}^{c_1,c_3}(s)\right) \\
&\qquad= k\left(\{(p,c_1,c_2),(q,c_4,c_3)\}\right)-k\left(\{(q,c_3,c_4)\}\right)-k\left(\{(p,c_1,c_2)\}\right)+k\left(\emptyset\right) \\
&\qquad= \sigma^2\left[\left(1-\delta_p^{c_1,c_2}\right)\left(1-\delta_q^{c_3,c_4}\right)\left(1-\delta_{pq}^{c_1,c_2,c_4,c_3}\right)-\left(1-\delta_q^{c_3,c_4}\right)-\left(1-\delta_p^{c_1,c_2}\right)+1\right] \\
&\qquad= \sigma^2\left[\delta_p^{c_1,c_2}\delta_q^{c_3,c_4}-\delta_{pq}^{c_1,c_2,c_4,c_3}\left(1-\delta_p^{c_1,c_2}\right)\left(1-\delta_q^{c_3,c_4}\right)\right].
\end{aligned}$$

Applying the same approximation to the normalized statistic gives:

(SM1.40)

$$\begin{aligned}
\mathbb{E}_{f\sim\mathcal{N}(0,K)}\left[\gamma_{p:c_1\to c_2;\,q:c_3\to c_4}(f)\right] &\approx \frac{\mathbb{E}_f\left[\Gamma_{p:c_1\to c_2;\,q:c_3\to c_4}(f)\right]}{\sqrt{\mathbb{E}_f\left[\mathbb{E}_{s:s_p=c_1,s_q=c_3}[w_{p:c_1\to c_2}(s|f)^2]\right]\mathbb{E}_f\left[\mathbb{E}_{s:s_p=c_1,s_q=c_3}[w_{q:c_3\to c_4}(s|f)^2]\right]}} \\
&= \frac{\delta_p^{c_1,c_2}\delta_q^{c_3,c_4}-\delta_{pq}^{c_1,c_2,c_4,c_3}(1-\delta_p^{c_1,c_2})(1-\delta_q^{c_3,c_4})}{2\sqrt{\delta_p^{c_1,c_2}\delta_q^{c_3,c_4}}} \\
&= \frac{1}{2}\sqrt{\delta_p^{c_1,c_2}\delta_q^{c_3,c_4}}-\delta_{pq}^{c_1,c_2,c_4,c_3}\frac{(1-\delta_p^{c_1,c_2})(1-\delta_q^{c_3,c_4})}{2\sqrt{\delta_p^{c_1,c_2}\delta_q^{c_3,c_4}}} \\
&= \frac{1}{2}\sqrt{\delta_p^{c_1,c_2}\delta_q^{c_3,c_4}}+\frac{\delta_{pq}^{c_1,c_2,c_3,c_4}}{1-\delta_{pq}^{c_1,c_2,c_3,c_4}}\frac{(1-\delta_p^{c_1,c_2})(1-\delta_q^{c_3,c_4})}{2\sqrt{\delta_p^{c_1,c_2}\delta_q^{c_3,c_4}}}.
\end{aligned}$$

Since $1-\delta_{pq}^{c_1,c_2,c_3,c_4}>0$, the sign of $\delta_{pq}^{c_1,c_2,c_3,c_4}$ determines whether the approximation to the expected similarity between the two mutational effects is higher or lower than under a GPK prior with the same single-mutation decay factors. Positive pairwise decay factors increase this similarity, whereas negative pairwise decay factors decrease it.

Finally, note that this new $\gamma_{p:c_1\to c_2;\,q:c_3\to c_4}(f)$ statistic can also be understood as the cosine similarity between vectors of mutational effects and thus can be used to quantify the degree of alignment of the effects of every pair of mutations across every possible background in which they can be introduced in an arbitrary sequence-function relationship $f$.

**SM1.6. Relationship with global epistasis models.** We illustrate in this section the FA kernel's ability to capture global or nonspecific epistasis patterns in single and multiple dimensions by considering some synthetic examples. The examples use a small sequence space of size $4^6$ (alphabet size 4, length 6). We will first consider a one-dimensional case where the FA kernel learns a one dimensional embedding $A$ that accurately represents the latent function $\phi$ in the GE model [SM21]. Then, we also consider a multi-dimensional case where the FA kernel learns a two dimensional embedding $A$, accurately learning two different latent functions $\phi_1$ and $\phi_2$ associated with Factor 1 and Factor 2. In this more complicated example, we demonstrate the FA kernel's ability to learn global epistasis without the single dimension restriction of the GE model.

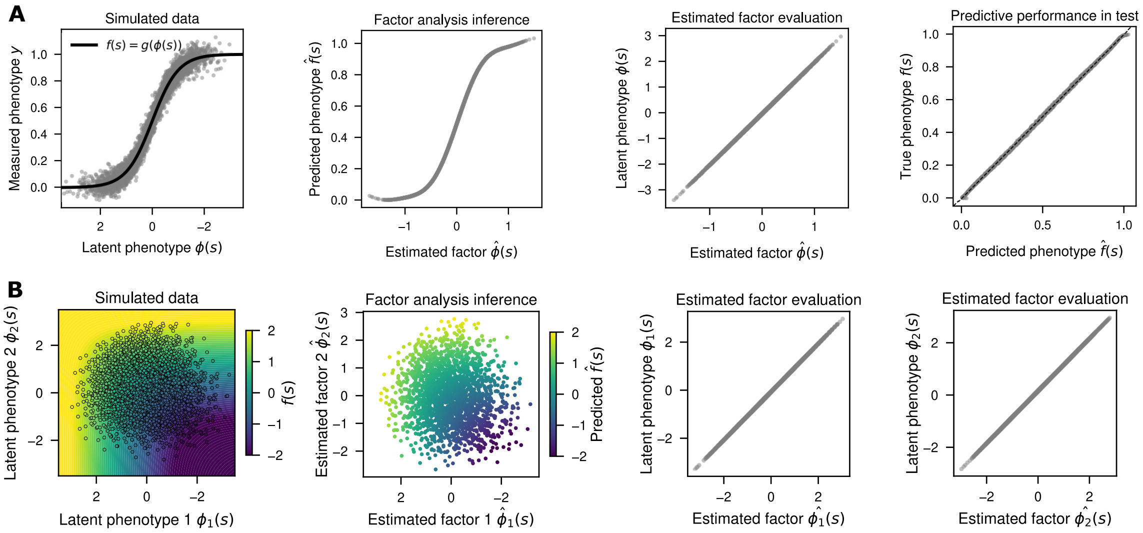


**Figure SM1.** *Examples demonstrating the FA kernel capturing global epistasis. (A) One-dimensional data is simulated to create DNA sequences of length six, whose fitness function resembles Equation SM1.41 (left). The FA kernel captures this behavior with one factor (left, center). The performance of the learned factor and the latent phenotype are compared as well as the predicted fitness against the true fitness (right, center). (B) The same analysis is done in two dimensions defined by Equation SM1.43.*

First, in Figure SM1A, we consider an example where $f(s) = g(\phi(s))$, where $g(\cdot)$ is a monotonic function and $\phi(s)$ is an additive function of the sequence $s$. In particular, we use a logistic function as it is commonly implemented for modeling epistatic sequence-function relationships:

$$g(\phi) = \frac{1}{1 + e^{-2\phi}}, \tag{SM1.41}$$

where $\phi$ is drawn randomly from the space of possible additive functions and is normalized to have a standard deviation of one (Figure SM1A, left). We then sample measurements $y$ with Gaussian error given by $\sigma_n^2 = 0.05$:

$$y_x \sim \mathcal{N}(f(x), \sigma_n^2), \tag{SM1.42}$$

where we use 50% of the data points for training and fit a Gaussian process model with an FA kernel that learns the linear embedding space $A$ via evidence maximization. We see that it recreates the global epistasis shape (Figure SM1A, center left) and note that the learned linear subspace accurately recapitulates the latent additive function $\phi$ (Figure SM1A, center right) and accurately predicts phenotype (Figure SM1A, right).

Next, we generalize these results to higher dimensions, since the FA kernel can have multiple factors. This enables the FA kernel to capture more expressive global epistatic relationships for cases like multi-peaked landscapes. We consider a model in which $\phi = [\phi_1, \phi_2]$ is a two-dimensional additive function and $(g \circ \phi)$ maps sequences to a single value. In this setting, we set $(g \circ \phi)$ to a two-dimensional function that commonly appears as a way to model two different valid configurations for functionality:

$$g(\phi_1, \phi_2) = \log(e^{\phi_1} + e^{\phi_2}) \tag{SM1.43}$$

We then sample measurements $y$ with Gaussian error given by $\sigma_n^2 = 0.01$:

$$y_x \sim \mathcal{N}(f(x), \sigma_n^2), \tag{SM1.44}$$

where we use 50% of the data points for fitting a Gaussian process model with a two-dimensional FA kernel. The learned factors produce an embedding that reflects the original $\phi$ after finding the linear transformation of the factors that minimizes the distances (Figure SM1B). These results suggest that the FA kernel can capture multidimensional global nonlinearities up to a linear transformation.

While the transformation was designed specifically to orthogonalize the two factors and maximize the variance explained by the first dimension, the resulting representation may not reflect true cause-and-effect relationships in the data. However, regardless of this chosen representation, the FA kernel's predictions are excellent in the two-dimensional latent phenotype model (Figure SM1).

**SM1.7. Relationship with LANTERN.** LANTERN [SM22] is another Gaussian process method that learns a linear latent space embedding of sequences and defines the prior covariance between sequences in terms of distances in this latent space. Here, we highlight the differences between the models and their implementations.

The LANTERN method defines a prior distribution on the latent space through the relative sizes of the embedding dimensions. The model learns an embedding matrix $W$ whose columns represent latent mutation effects, with $z(s) = Wx(s)$, where $x(s)$ is the one-hot encoding of sequence $s$. A prior is set on the elements of $W$, where $w_{k,j} \sim N(0, \alpha_k^{-1})$, with $\alpha_k$ being the precision (or inverse variance) of dimension $k$. The goal is to learn a sparse $W$ that is interpretable with few dimensions. In their paper, they learn up to eight dimensions. In contrast, our model learns the embedding without a hierarchical prior on the relative importance of the embedding dimensions. By not restricting these dimensions, the FA model retains the flexibility of the full embedding space. When we instead learn a low-rank embedding space, we restrict the class of sequence-function relationships that can be represented. In our paper, we consider both options by learning the full embedding space for short sequence motifs and by learning low-rank approximations for longer sequences, ranging from one to eight dimensions.

Furthermore, while LANTERN and our FA kernel both define sequence embeddings in a latent space, we use different kernel functions for determining the prior covariance from the latent space distances. Our kernel function decays exponentially with the squared distance in latent space $d(s, s')^2 = ||\phi(s) - \phi(s')||_2^2$,

$$k(s, s') = \sigma^2 \exp(-c^2 d(s, s')^2),$$

whereas LANTERN uses a rational quadratic (RQ) kernel of the form

$$k(s, s') = \sigma^2 \left(1 + \frac{d(s, s')^2}{2\eta}\right)^{-\eta}.$$

As the shape parameter $\eta$ tends to infinity, the RQ kernel converges to the FA kernel with $c = 1/\sqrt{2}$. However, in practice, [SM22] does not take this limit. Using the exponential decay rather than the RQ allows us to mathematically relate the FA kernel to other classical kernels defined in the literature (see Section 3 in the main text and Section SM1.1 in this document for more details).

Finally, there are differences in the implementation of these two models. LANTERN learns the embedding space via variational inference and inducing points, whereas we use evidence maximization with exact inference. Furthermore, we parameterize our models differently. LANTERN works in an overparameterized space and regularizes this space by setting priors on the mutational effect space. We address the degrees of freedom arising from the unconstrained directions of the one-hot encoding and the nonidentifiability of the embedding matrix by fixing a gauge and establishing a systematic approach to optimizing and interpreting the inferred embedding matrix (see Section 5.1 in the main text for more details).

**SM1.8. Precomputed embedding details.** We expand on the discussion in the main text (Section 3.3) about applying precomputed features to our Gaussian process model. Here, we construct an embedding kernel using features obtained by singular value decomposition (SVD) [SM2] of natural sequence data. The leading singular vectors capture the dominant structure in these datasets. Truncating the SVD to $k$ directions reduces the embedding dimension from $\alpha\ell$ to $k$, allowing us to choose a smaller embedding dimension independent of the sequence length. However, one caveat is that patterns in evolutionary data may not reflect the same patterns in measured laboratory data. We therefore use this precomputed embedding to define the covariance kernel of our Gaussian process prior and learn the relative scales of its features from the training data.

The matrix on which we perform the SVD can be constructed using one-hot encodings of multiple sequence alignments (MSAs) sequences or nonlinear representations from protein language models such as ESM [SM8]. These representations provide a matrix $X$ with dimensions $N \times \alpha\ell$ for one-hot encoded sequences or $N \times 1280\ell$ for 1280-dimensional ESM representations per position where $N$ is the number of sequences. For simplicity, we describe the construction using a generic encoding $x(s)$ of a sequence $s$.

We compute the SVD

$$X = U\Lambda V^T, \tag{SM1.45}$$

where U has orthonormal columns derived by computing the eigenvectors of $XX^T$ and has dimension $N \times N$, $\Lambda$ is rectangular with singular values $\sigma_i$ along the main diagonal corresponding to the square root eigenvalues of $X^T X$ and $XX^T$, and $V^T$ has orthonormal rows by computing the eigenvectors of $X^T X$ and has dimension $\alpha\ell \times \alpha\ell$. In practice, we reduce the dimensionality of this representation by selecting the top $k$ singular values in $\Lambda$ and their associated right singular vectors represented by the rows of $V^T$.

Multiplying Equation SM1.45 by $V$ gives

$$XV = U\Lambda. \tag{SM1.46}$$

If $x(s)^i$ is the $i$th encoded sequence of $X$ and $v_j$ is the $j$th right singular vector of V, then we can rewrite the above Equation SM1.46 as:

$$x(s)^i \cdot v_j = \lambda_j u_{ij} \tag{SM1.47}$$

By defining $\beta_j(x(s)^i) = x(s)^i \cdot v_j$, we derive our $j$th feature as

$$\boldsymbol{\beta}^j = Xv_j \tag{SM1.48}$$

This provides a way to go between sequence space and SVD space, where each feature $\boldsymbol{\beta}^j$ is the projection of the encoded sequences onto the right singular vector $v_j$. Hence, these features capture the importance of certain positions of the select sequences.

The dimension of $V$, which determines the number of features that describe the evolutionary patterns, depends on both the number of sequences in $X$, $N$, and the number of dimensions used to encode each sequence relative to $N$. This can differ between encoding methods. In the context of one-hot encoding of MSA sequences, if $X$ is $N \times \alpha\ell$ and $N < \alpha\ell$, then $V$ has dimension $\alpha\ell \times N$. In other words, there are $N$ features. On the other hand, if $N \geq \alpha\ell$, then $V$ has dimension $\alpha\ell \times \alpha\ell$; there are $\alpha\ell$ features. The same reasoning can be used for $ESM$ embeddings, in which the encoding dimension is $1280\ell$. In the case of the GFP dataset, on which we test this method, we have that $N < \alpha\ell$ and $N < 1280\ell$, so we only consider the $N$ feature case. When we truncate the number of features for $k \ll \alpha\ell$ or $k \ll 1280\ell$, this leads to $k$ total features.

Using these features, we define the embedding kernel from Equation 3.1 in the main text

$$k(s, s') = \sigma^2 \exp\left(-\|\phi(s) - \phi(s')\|^2\right), \tag{SM1.49}$$

where $\phi(s) = Ax(s) = D_k^{1/2} V_k^T x(s)$ for the top $k$ residue eigenvectors of $V$, $V_k$, and diagonal matrix $D_k = \text{diag}\left(c_1^2, \ldots, c_k^2\right)$ describing the embedding scale parameters $c_r > 0$ for each feature.

We find that this kernel directly describes an ARD kernel by substituting in $\phi$:

$$k(s, s') = \sigma^2 \exp\left(-\sum_{r=1}^{k} c_r^2 (\beta_r(x(s)) - \beta_r(x(s')))^2\right). \tag{SM1.50}$$

This definition highlights the embedding scale parameters describing the contributions of each feature to the overall covariance. We learn the embedding scale values and the prior variance $\sigma^2$ by maximizing the Gaussian process marginal likelihood on the training data.

**SM1.8.1. Results on GFP using precomputed features.** We compare our results of learning features directly on the data via the FA kernel with precomputing them from natural sequence data. We consider two methods in which to precompute features: 1) finding the eigenvectors from the jackhmmer MSA (described in Section SM2.1.4), and 2) the per-position embeddings (a vector of length $1280\ell$ for each sequence) or the mean embeddings (a vector of length 1280 for each sequence) from the ESM2 model [SM8] on lab derived training data. Both the MSA and per position ESM embeddings use the SVD method described in Section SM1.8. The mean ESM embedding, however, is defined by the squared difference between the embeddings:

$$k(s, s') = \sigma^2 \exp\left(-||ESM(s) - ESM(s')||_2^2\right) \tag{SM1.51}$$

where $ESM(s)$ is the 1280-dimensional embedding of sequence $s$, obtained by averaging its $\ell$ position-specific embedding vectors, each of dimension 1280. Since no SVD is being performed, there is no truncation on the number of features we define in this setting. Thus, we compare the predictive performance of all the precomputed methods with an increasing number of features (10, 50, and 250) in Figure SM2.

We compare the performance of these precomputed features with a randomly generated feature set, described by finding the eigenvectors of randomly generated sequences in which each amino acid at each position was chosen uniformly at random. This feature set defines a baseline of how well the precomputed features should perform. As we would expect, all methods employing the SVD truncation increase performance as the number of features increases, and in particular, with enough features, the random set spans enough of sequence space to perform well, if not better in many instances, than the precomputed methods (Figure SM2). The ESM mean embedding performs the same throughout as there is no SVD to truncate the number of features. These results suggest that the precomputed features considered here do not provide an advantage over randomly generated features for predicting the sequence-function relationships in these datasets. Comparing these results with the learned embeddings in the main text (Section 6.4), we find that the one FA feature outperforms even the 250 features of the precomputed methods. This suggests it is better to learn features directly on this dataset, rather than precomputing them, and that perhaps Euclidean distances in the ESM latent space may not be the best proxies for fluorescence similarity.

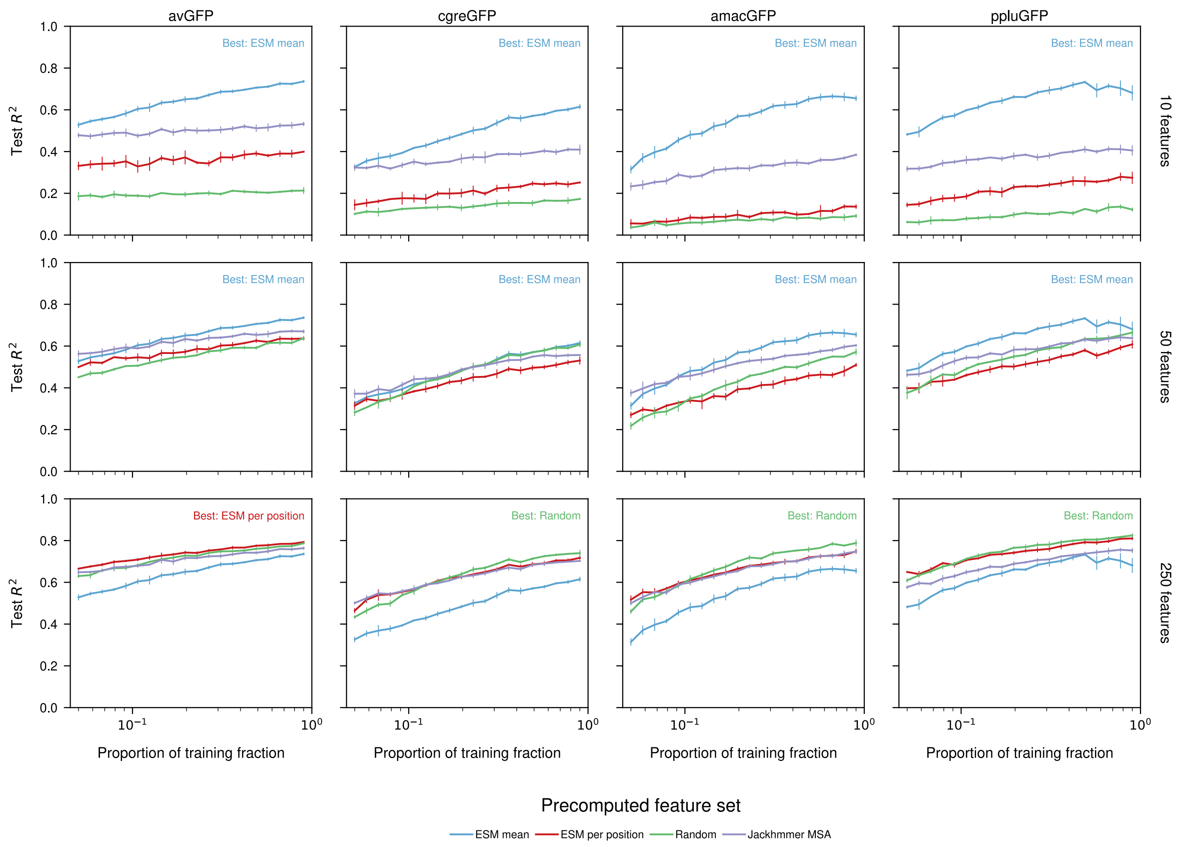


**Figure SM2.** *Predictive performance of precomputed feature sets with an increase of features. Rows compare a particular number of feature from the SVD truncation. Columns define a particular GFP protein. Each line represents a different precomputed method.*

**SM2. Supplementary Methods.** Here, we describe the details of implementing our Gaussian process model by discussing how we process each dataset, how we train our model and deal with missing alleles in the GFP dataset, and how we analyze our results by making predictions and assessments.

### SM2.1. Processing data.

**SM2.1.1. GB1.** We processed the GB1 dataset following the procedure of earlier work [SM26, SM25, SM24]. Using raw sequencing counts for each variant $s$ before selection ($c^{input}$) and after selection ($c^{sel}$) [SM23], we defined the binding strength $y_s$ as the log-enrichment of the variant normalized to that of the wild-type sequence. We also estimated the error variance $\sigma_s^2$ of each value from the counts [SM17]:

$$y_s = \log\left(\frac{c_s^{sel} + 0.5}{c_s^{input} + 0.5}\right) - \log\left(\frac{c_{wt}^{sel} + 0.5}{c_{wt}^{input} + 0.5}\right) \tag{SM2.1}$$

$$\sigma_s^2 = \frac{1}{c_s^{input} + 0.5} + \frac{1}{c_s^{sel} + 0.5} + \frac{1}{c_{wt}^{input} + 0.5} + \frac{1}{c_{wt}^{sel} + 0.5} \tag{SM2.2}$$

**SM2.1.2. AAV2.** The AAV2 dataset was downloaded and used directly from the ProteinGym database [SM11] without externally supplied estimates of the experimental error variance. For this dataset, we assumed a common experimental variance $\sigma_n^2$ that is inferred together with the kernel parameters as described in Section SM2.2.1.

**SM2.1.3. Self-splicing intron.** We processed the self-splicing intron dataset from [SM19], which contains the combinatorially-complete library at the eight variable nucleotide positions 2-5 and 18-21. We retained all $4^8 = 65,536$ DNA sequences and used the reported $\log_2$ fold-change at 30°C as the phenotype $y_s$. We did not assign experimental error variances to these measurements; instead, a single noise variance $\sigma_n^2$, shared across all sequences, was learned during Gaussian process fitting.

**SM2.1.4. GFP.**

*GFP: Lab-derived data.* We obtained the avGFP dataset from [SM18] and the amacGFP, cgreGFP, and ppluGFP2 datasets from [SM7]. Each dataset reports nucleotide genotypes and their corresponding amino-acid genotypes as mutations relative to the wild-type sequence. We reconstructed full amino-acid sequences by applying the reported amino-acid mutations to the corresponding wild-type sequence. Experimental replicates had already been summarized as one brightness score per nucleotide genotype. The avGFP scores were reported on a log-brightness scale; for the other proteins, we used the log of the reported brightness. We removed amino-acid sequences containing a stop codon (*) or unknown residue (.).

Because different nucleotide genotypes can encode the same amino-acid sequence, we grouped the measurements by amino-acid sequence. For each amino-acid sequence $s$, we defined its phenotype as

$$y_s = \frac{1}{n_s} \sum_{s^{\text{nt}} \in \{s^{nt} | T(s^{nt}) = s\}} b_{s^{\text{nt}}}. \tag{SM2.3}$$

where $n_s$ is the number of distinct nucleotide sequences $s^{nt}$ whose translation $T(s^{nt})$ forms the same amino-acid sequence $s$ and $b_{s^{nt}}$ is the log-brightness reported for each $s^{nt}$.

Replicate standard deviations were available for only a small fraction of the avGFP nucleotide sequences. We therefore estimated one error variance per wild-type dataset from the variation among distinct nucleotide sequences that encode the same amino-acid sequence. We pooled this variation over amino-acid sequences with more than one nucleotide sequence present in the dataset to give a general and more reliable estimate of the experimental error variance,

$$\hat{\sigma}^2 = \frac{1}{n - m} \sum_s \sum_{s^{\text{nt}} \in \{s^{nt} | T(s^{nt}) = s\}} \left(b_{s^{\text{nt}}} - y_s\right)^2, \tag{SM2.4}$$

where $n$ is the number of nucleotide sequences and $m$ is the number of amino acid sequences present in the dataset. We then assumed that the experimental variance for each sequence $s$

was the same $\sigma_s^2 = \hat{\sigma}^2$ without dividing by the number of contributing nucleotide genotypes. This approximation improved numerical stability during model fitting. For avGFP, 1202 amino-acid sequences have $n_s \geq 2$, encoded by 5573 nucleotide sequences in total, giving $\sigma_s^2 = 0.0106$. The other proteins gave $\sigma_s^2 = 0.00135$ (amacGFP), 0.00150 (cgreGFP), and 0.00098 (ppluGFP2).

*GFP: multiple sequence alignments.* We used the 71 homologous sequences reported by [SM7] to construct a multiple sequence alignment (MSA). We searched UniProt using jackhmmer [SM14]. We removed fragments and sequences identified as nonfluorescent, appended the four wild-type sequences (avGFP, amacGFP, cgreGFP, and ppluGFP2), and aligned the sequences using MUSCLE 3.8.31 [SM5]. This MSA contained 390 sequences. For each wild-type reference, we removed alignment columns corresponding to gaps in that reference, leaving 238 positions for avGFP and amacGFP, 235 for cgreGFP, and 222 for ppluGFP2. To reduce the influence of overrepresented groups of similar sequences, we clustered the sequences at an 80% sequence-identity threshold using MMseqs2 [SM20]. Let $c_k$ be the number of sequences in cluster $k$. For sequence $s_i$ in cluster $k(i)$, we assigned the weight $w_i = 1/c_{k(i)}$. We constructed a matrix $X$ whose $i$th row is the one-hot encoding of $s_i$ over the 20 standard amino acids at each retained position. Gaps and nonstandard characters were represented by an all-zero block at the corresponding position. We then multiplied each row by its sequence weight, giving $X_w = D_w^{1/2} X$, where $D_w = \mathrm{diag}(w_1, \ldots, w_{390})$. We computed the reduced singular value decomposition $X_w = U \Lambda V^T$. The columns of $V$ are directions in the one-hot feature space, ordered by decreasing singular value. The rank of $X_w$ was 369 for avGFP and amacGFP, 368 for cgreGFP, and 367 for ppluGFP2. All feature truncations used in the models, including 250 features, were below these full ranks.

### SM2.2. Gaussian process model fitting.

**SM2.2.1. Training parameters.** We used EpiK [SM24] to fit Gaussian process models to each dataset. EpiK builds on GPyTorch [SM6] and KeOps [SM3] to run inference on GPUs. We used a zero-mean prior. For datasets with experimental error-variance estimates, the observation-noise variance for sequence $s$ was $\sigma_s^2 + \sigma_n^2$, where $\sigma_n^2$ is a learned term common to all sequences. For GFP, $\sigma_s^2$ was the pooled estimate for the corresponding protein. When no experimental error-variance estimate was available, the observation-noise variance was $\sigma_n^2$. This learned term captures sequence-independent noise not accounted for by the supplied estimates [SM24]. Kernel hyperparameters were fit with Adam optimizer with a learning rate 0.02 to maximize the marginal likelihood, for 200 iterations on GB1, 500 on the self-splicing intron, 300 on AAV2 (500 for Jenga and GPK), and 1000 on GFP. The log-determinant term in the evidence was approximated with 300 Lanczos vectors and 50 random vectors for the trace estimate, with a conjugate-gradient tolerance of 0.1 for hyperparameter training and $10^{-4}$ for computing the posterior estimates [SM24]. For cross-validation fits of the FA kernel on GFP and AAV2, we used a KeOps [SM3] covariance with 100 Lanczos vectors. The Jenga-FA kernel used 20 trace samples.

The initialization of the hyperparameters, and the number of restarts, differed between the site-specific product kernels and the factor analysis kernels. For the site-specific product kernels (Geometric, Connectedness, Jenga, and GPK), the prior variance started at the empirical variance of the phenotype, and all decay factors started at one value shared across sites and mutations. This value was chosen so that the prior correlation falls exponentially with the number of mutations and reaches 0.1 between the two most distant sequences. Because the evidence is non-convex and may have local optima, we fit the complete datasets from 3 random initializations and kept the run with the highest evidence. In practice, the random initializations produced similar optimums [SM15]. The FA kernel was initialized with the same empirical prior variance. Its factors were obtained from a deterministic factorization that places equal weight on every allele observed in the training data, with the factor scales initialized to 1. We fit models parametrized by a full rank $M$ matrix for GB1 and the self-splicing intron, and models parametrized by one factor for each GFP protein and eight factors for AAV2. The Jenga-FA kernel used the same factor initialization. Its prior variance was initialized to 1, and its additional per-allele diagonal weights were initialized to 1. On AAV2 this kernel used three factors. Both FA kernels were fit from this single initialization.

**SM2.2.2. Missing data in GFP.** When using full-length protein datasets in our models, we must account for unobserved allele-position combinations. Some allele-position combinations are absent from the training data, and we therefore constrain their embedding coefficients to remain zero during optimization. In particular, the GFP dataset has many such missing combinations. Therefore, when we project a sequence into the gauge-fixed space, we want to ensure that missing allele-positions that are masked by zero remain masked after applying $PP^T$. We do this by considering a masked version of $Q$ described in the main text in Section 5.1.

Consider the original gauge-fixing matrix $Q = \text{blockdiag}(I_\alpha - \frac{1}{\alpha}\mathbf{1}_\alpha\mathbf{1}_\alpha^T)$. We construct an analogous matrix $\tilde{Q}$ that fixes the gauge after zeroing values associated to allele-position combinations that do not appear in the training set. To keep the sum of the alleles at each position zero, we replace $\frac{1}{\alpha}$ with $\frac{1}{\alpha_p}$, where $\alpha_p = |S_p|$ is the number of seen alleles at position $p$, $S_p$ is the set of alleles seen at position $p$, and $a_i$ is a particular allele in the alphabet.

We define $\tilde{Q}$ as the block diagonal matrix with

$$\tilde{Q}_p = \begin{bmatrix} \left(1-\frac{1}{\alpha_p}\right)\mathbb{1}_{a_1\in S_p} & \left(-\frac{1}{\alpha_p}\right)\mathbb{1}_{a_1\in S_p}\mathbb{1}_{a_2\in S_p} & \cdots & \left(-\frac{1}{\alpha_p}\right)\mathbb{1}_{a_1\in S_p}\mathbb{1}_{a_\alpha\in S_p} \\ (-\frac{1}{\alpha_p})\mathbb{1}_{a_2\in S_p}\mathbb{1}_{a_1\in S_p} & (1-\frac{1}{\alpha_p})\mathbb{1}_{a_2\in S_p} & \cdots & \left(-\frac{1}{\alpha_p}\right)\mathbb{1}_{a_2\in S_p}\mathbb{1}_{a_\alpha\in S_p} \\ \vdots & \vdots & \ddots & \vdots \\ \left(-\frac{1}{\alpha_p}\right)\mathbb{1}_{a_\alpha\in S_p}\mathbb{1}_{a_1\in S_p} & \left(-\frac{1}{\alpha_p}\right)\mathbb{1}_{a_\alpha\in S_p}\mathbb{1}_{a_2\in S_p} & \cdots & \left(1-\frac{1}{\alpha_p}\right)\mathbb{1}_{a_\alpha\in S_p} \end{bmatrix} \tag{SM2.5}$$

along its diagonal for each position $p$ where $\tilde{Q}_p \in \mathbb{R}^{\alpha\times\alpha}$. This can be thought of as a $\mathbb{R}^{\alpha_p\times\alpha_p}$ matrix with $I_{\alpha_p} - \frac{1}{\alpha_p}\mathbf{1}_{\alpha_p}\mathbf{1}_{\alpha_p}^T$ indexed only on the seen alleles and then inserted into a $\mathbb{R}^{\alpha\times\alpha}$ zero matrix at the appropriate allele-positions, keeping the rows and columns of the unseen alleles zero.

Let $S_p^c$ be the set of these unseen alleles at position $p$ and $S^c$ be the set of unseen allele-position indices, so that $|S^c| = k = \sum_{p=1}^{\ell}(\alpha-\alpha_p)$ is the total number of missing alleles. When there are no missing allele-position combinations, we perform our optimization of the latent factors in an $(\alpha-1)\ell$ dimensional space whose dimensionality matches the dimensionality of sequence space and then transform these factors into the gauge-fixed $\alpha\ell$ dimensional space via a matrix $P$. Eliminating $k$ allele-position combinations decreases the dimensionality of sequence space in which we want to optimize by $k$ to $r = \sum_p(\alpha_p - 1) = (\alpha - 1)\ell - k$. We analogously compute $\tilde{P}$ a matrix that brings the factors from the $r$-dimensional optimization space into the gauge-fixed $\alpha\ell$ dimensional space. To define $\tilde{P}$, we write the singular value decomposition of $\tilde{Q}$ as $\tilde{Q} = U\Lambda V^T$. It has exactly $r$ nonzero singular values, so keeping only those gives the truncated SVD $\tilde{Q} = U_r\Lambda_r V_r^T$, where $U_r, V_r \in \mathbb{R}^{\alpha\ell\times r}$ have orthonormal columns and $\Lambda_r$ is the invertible $r \times r$ diagonal matrix of nonzero singular values. We take $\tilde{P} := U_r \in \mathbb{R}^{\alpha\ell\times((\alpha-1)\ell-k)}$.

Here we establish that $\tilde{P}$ maps to gauge-fixed vectors with zeros at the unseen positions.

Claim SM2.1. *$Im(\tilde{P}) = \{v \in \mathbb{R}^{\alpha\ell} | \mathbf{1}_\alpha^T v_p = 0$ for $v_p \in \mathbb{R}^\alpha$, and $v_{a,p} = 0$ when $a \in S_p^c\}$.*

First note that $Im(\tilde{P}) = Im(\tilde{Q})$. Indeed, $\tilde{Q}x = \tilde{P}(\Lambda_r V_r^T x)$ for every $x$, so $Im(\tilde{Q}) \subseteq Im(\tilde{P})$. Conversely, since $V_r^T V_r = I_r$, we have $\tilde{Q}\,(V_r\Lambda_r^{-1}) = U_r\Lambda_r V_r^T V_r\Lambda_r^{-1} = U_r = \tilde{P}$, so every column of $\tilde{P}$ lies in $Im(\tilde{Q})$, which gives $Im(\tilde{P}) \subseteq Im(\tilde{Q})$.

Let $G \subseteq \mathbb{R}^{\alpha\ell}$ be the gauge-fixed space and $F \subseteq \mathbb{R}^{\alpha\ell}$ be vectors with the unseen allele-position entries frozen at zero. Let $\tilde{Q}$ be the masked version of $Q$ and $\tilde{P}$ be the orthonormal basis of $Im(\tilde{Q})$. Then $Im(\tilde{P}) = F \cap G$.

*Proof.* To say that $Im(\tilde{P}) = F \cap G$ would mean the columns of $\tilde{P}$ span the gauge-fixed vectors whose entries corresponding to the missing allele-position combinations are masked as zero. Since $Im(\tilde{P}) = Im(\tilde{Q})$, it suffices to show $Im(\tilde{Q}) = F \cap G$.

Since $\tilde{Q}$ is block diagonal, for any $x \in \mathbb{R}^{\alpha\ell}$ the $p$th subvector of $\tilde{Q}x$ is $\tilde{Q}_p x_p$, where $x_p \in \mathbb{R}^\alpha$ is the subvector of $x$ for position $p$. We will use this in both directions.

First we will show that $Im(\tilde{Q}) \subseteq F \cap G$. Suppose $v \in Im(\tilde{Q})$. Then, $v = \tilde{Q}x$ for some $x \in \mathbb{R}^{\alpha\ell}$, and $v_p = \tilde{Q}_p x_p$ for each position $p$. The rows of $\tilde{Q}_p$ corresponding to unseen alleles are zero, so $v_{a,p} = 0$ whenever $a \in S_p^c$, and hence $v \in F$. Furthermore, the submatrix indexed on the seen alleles has column sums $\mathbf{1}_{\alpha_p}^T - \frac{1}{\alpha_p}(\mathbf{1}_{\alpha_p}^T \mathbf{1}_{\alpha_p})\mathbf{1}_{\alpha_p}^T = 0$, and on the unseen alleles they contribute zero so their sum is also zero. Hence $\mathbf{1}_\alpha^T \tilde{Q}_p = 0$. Therefore, $\mathbf{1}_\alpha^T v_p = \mathbf{1}_\alpha^T \tilde{Q}_p x_p = 0$ for each $p$, and hence $v \in G$. Therefore $Im(\tilde{Q}) \subseteq F \cap G$.

Now, we will show $F \cap G \subseteq Im(\tilde{Q})$. Let $v \in F \cap G \subseteq \mathbb{R}^{\alpha\ell}$. Then $v_{a,p} = 0$ whenever $a \in S_p^c$ and $\mathbf{1}_\alpha^T v_p = 0$ for each position $p$. Fix $p$. We have that $\tilde{Q}_p v_p$ acts only on the seen alleles. Because the entries of $v_p$ sum to zero, this gives the submatrix on only the seen alleles $(I_{\alpha_p} - \frac{1}{\alpha_p}\mathbf{1}_{\alpha_p}\mathbf{1}_{\alpha_p}^T)v_p|_{S_p} = v_p|_{S_p} - \frac{1}{\alpha_p}\mathbf{1}_{\alpha_p}(\mathbf{1}_{\alpha_p}^T v_p|_{S_p}) = v_p|_{S_p}$ where $v_p|_{S_p}$ is the vector $v_p$ restricted to the seen alleles at position $p$. Since the unseen entries of $v_p$ and the corresponding rows of $\tilde{Q}_p$ are zero, $(\tilde{Q}_p v_p)|_{S_p^c} = 0 = v_p|_{S_p^c}$. Thus, $\tilde{Q}_p v_p = v_p$ for every $p$. Since $\tilde{Q}$ is block diagonal, $\tilde{Q}v = v$, so $v \in Im(\tilde{Q})$. Therefore $F \cap G \subseteq Im(\tilde{Q})$.

Thus $Im(\tilde{Q}) = F \cap G$, and since $Im(\tilde{P}) = Im(\tilde{Q})$, we get the result we wanted, $Im(\tilde{P}) = F \cap G$. In other words, $\tilde{P}$ preserves the masking of missing allele-position combinations and the gauge-fixing. ■

**SM2.2.3. Predictions and analysis.** We assessed predictive performance using the same random seeds to create train-test splits for each of the datasets and each of the kernels, ranging from 5% to 90% of the data for training. This resulted in 60 splits per dataset and we randomly selected up to 5,000 of the remaining sequences for testing. For each split, we refit the kernel hyperparameters by maximizing the marginal likelihood using only the training data and the optimization settings described above. Predictions were defined by the posterior mean of the Gaussian process. We compared our prediction to the sequences' true measurements by using the $R^2$ across the three random seed splits at each training fraction.

**SM2.3. Global epistasis models.** Global epistasis models for each of the four GFP data set were fit using MAVE-NN v1.0.2 [SM21] with an unregularized additive genotype-phenotype map, a monotonic global epistasis non-linearity with a regularization of 0.01 on the measurement process parameters, and Gaussian homoskedastic noise. Models were trained by minimizing the variational information with a batch size equal to the size of the dataset for 5000 epochs with a learning rate of 0.01 to ensure model convergence. All other parameters were set to their default values.

**SM2.4. Visualization of GB1 and self-splicing intron landscapes.** We generated low-dimensional representations of the inferred sequence-function relationships using gpmap-tools [SM9]. In the resulting visualization, squared distances optimally approximate the commute time—the sum of the expected times to evolve from one sequence to another and back—under a model of molecular evolution where the population evolves under selection for higher phenotypic values. The different diffusion axes capture directions in which evolution is the slowest and typically separate qualitatively different sets of sequences with high phenotypic values that are separated by at least partial fitness valleys [SM10, SM25, SM4].

The coordinates of the visualization were computed using the maximum a posteriori (MAP) estimate for the complete sequence-function map of protein GB1 and self-splicing intron inferred with the complete dataset under the full rank FA model. We used uniform mutation rates for every possible mutation and selection strength was set such that the stationary distribution of the resulting Markov chain had a phenotypic mean that matches the wild-type phenotypic value for the GB1 landscape and an average $\log_2$ fold-change score of 1.7 for the self-splicing intron landscape (corresponding to the 97% percentile in the distribution of $\log_2$ fold-change values).

**SM2.5. GFP posterior sampling.** In Figure 6B (left) in the main text, we subsample the points to draw the posterior distribution draws. Here, we will describe this process. The posterior is conditioned on every sequence in the training split. The posterior mean is the predicted mean for each test sequence. To draw functions from this posterior, the test sequences are sorted by their Factor 1 coordinate, $a_1^\top x(s)$, and 120 sequences are retained at equal intervals along that order with an additional 51 sequences more finely covering the sparser region associated with Factor 1 values $-5$ to $-15$. This enables the curves to cover the observed range of Factor 1. Eight samples are then drawn from the joint posterior for the 161 sequences, which are represented by the grey curves in Figure 6B.

**SM3. Supplementary Figures.** In this section, we include additional figures corresponding to the main text's Section 6.3. We find no evidence of sparse structure for the embedding matrix of AAV2 by applying the rotation test (Figure SM3) [SM12] and show all the orthogonal factors derived from the eigendecomposition of the estimated rank-eight $\tilde{M}$ matrix (Figure SM4).

In Figure SM3, the similarity between the reconstruction of the rotated $\tilde{M}$ line and the non-rotated $\tilde{M}$ line suggests there is no sparse structure in this dataset [SM12].

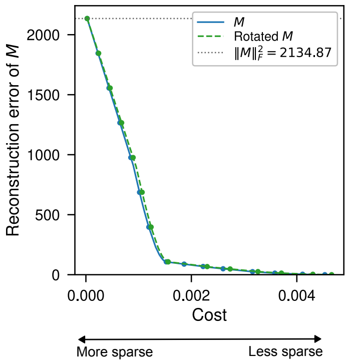


**Figure SM3.** *Evaluation of the sparse structure decomposition (SSD) of the learned $\tilde{M}$ matrix. Comparison of the reconstruction error and the total cost function for a wide range of sparsity penalties of the $M$ matrix (solid blue line) and a rotated version (dashed red line). Error is quantified by the Frobenius norm of the difference between the $M$ matrix and the reconstructed matrix from the sparse factors.*

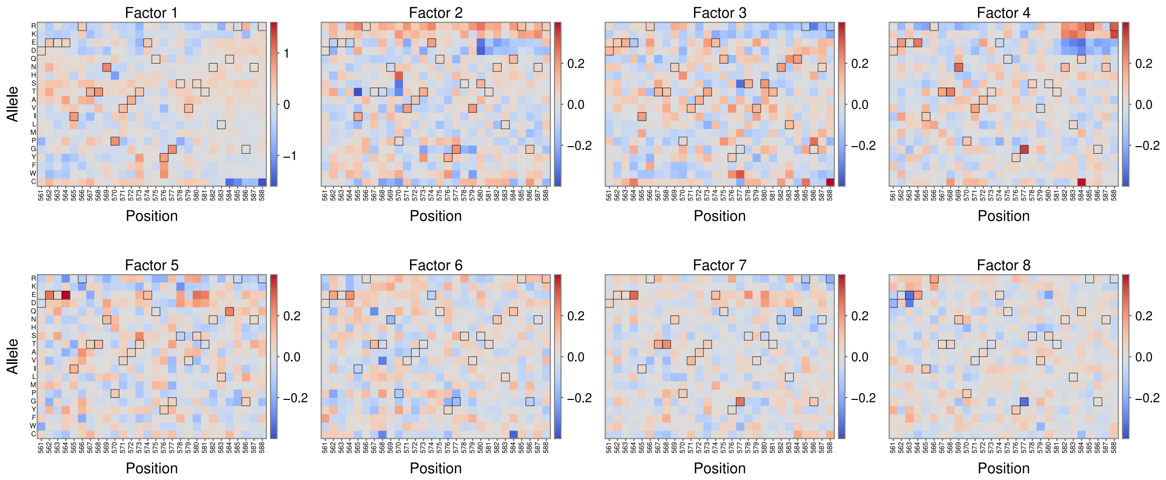


**Figure SM4.** *Applying the FA kernel on the AAV2 capsid assembly. Heatmaps representing the independent contributions that each allele at each position has for all eight factors.*

### REFERENCES


[1] C. Bank, S. Matuszewski, R. T. Hietpas, and J. D. Jensen, *On the (un)predictability of a large intragenic fitness landscape*, Proceedings of the National Academy of Sciences, 113 (2016), pp. 14085–14090, https://doi.org/10.1073/pnas.1612676113, https://pnas.org/doi/full/10.1073/pnas.1612676113 (accessed 2024-01-09).

[2] A. R. Baxter-Koenigs, G. El Nesr, and D. Barrick, *Singular value decomposition of protein sequences as a method to visualize sequence and residue space*, Protein Science, 31 (2022), p. e4422, https://doi.org/10.1002/pro.4422, https://onlinelibrary.wiley.com/doi/10.1002/pro.4422.

[3] B. Charlier, J. Feydy, J. A. Glaunès, F.-D. Collin, and G. Durif, *Kernel operations on the GPU, with autodiff, without memory overflows*, Journal of Machine Learning Research, 22 (2021), pp. 1–6.

[4] W.-c. Chen, J. Zhou, J. M. Sheltzer, J. B. Kinney, and D. M. Mccandlish, *Field-theoretic density*

*estimation for biological sequence space with applications to 5 splice site diversity and aneuploidy in cancer*, Proc. Natl. Acad. Sci. USA, (2021), https://doi.org/10.1073/pnas.2025782118.

[5] R. C. Edgar, *MUSCLE: multiple sequence alignment with high accuracy and high throughput*, Nucleic Acids Research, 32 (2004), pp. 1792–1797, https://doi.org/10.1093/nar/gkh340.

[6] J. Gardner, G. Pleiss, K. Q. Weinberger, D. Bindel, and A. G. Wilson, *Gpytorch: Blackbox matrix-matrix gaussian process inference with gpu acceleration*, Advances in neural information processing systems, 31 (2018).

[7] L. Gonzalez Somermeyer, A. Fleiss, A. S. Mishin, N. G. Bozhanova, A. A. Igolkina, J. Meiler, M.-E. Alaball Pujol, E. V. Putintseva, K. S. Sarkisyan, and F. A. Kondrashov, *Heterogeneity of the GFP fitness landscape and data-driven protein design*, eLife, 11 (2022), p. e75842, https://doi.org/10.7554/eLife.75842, https://elifesciences.org/articles/75842.

[8] Z. Lin, H. Akin, R. Rao, B. Hie, Z. Zhu, W. Lu, N. Smetanin, R. Verkuil, O. Kabeli, Y. Shmueli, A. Dos Santos Costa, M. Fazel-Zarandi, T. Sercu, S. Candido, and A. Rives, *Evolutionary-scale prediction of atomic-level protein structure with a language model*, Science, 379 (2023), pp. 1123–1130, https://doi.org/10.1126/science.ade2574, https://www.science.org/doi/10.1126/science.ade2574.

[9] C. Martí-Gómez, J. Zhou, W.-C. Chen, A. Stoltzfus, J. B. Kinney, and D. M. McCandlish, *Inference and Visualization of Complex Genotype–Phenotype Maps*, Molecular Biology and Evolution, 43 (2026), p. msag023, https://doi.org/10.1093/molbev/msag023, https://academic.oup.com/mbe/article/doi/10.1093/molbev/msag023/8456298.

[10] D. M. McCandlish, *Visualizing Fitness Landscapes*, Evolution, 65 (2011), pp. 1544–1558, https://doi.org/10.1111/j.1558-5646.2011.01236.x, https://academic.oup.com/evolut/article/65/6/1544/6854390.

[11] P. Notin, A. Kollasch, D. Ritter, L. V. Niekerk, S. Paul, H. Spinner, N. Rollins, A. Shaw, R. Orenbuch, R. Weitzman, J. Frazer, M. Dias, D. Franceschi, Y. Gal, and D. Marks, *ProteinGym: Large-Scale Benchmarks for Protein Fitness Prediction and Design*, in Advances in Neural Information Processing Systems 36, New Orleans, Louisiana, USA, 2023, Neural Information Processing Systems Foundation, Inc. (NeurIPS), pp. 64331–64379, https://doi.org/10.52202/075280-2810, http://www.proceedings.com/075280-2810.html.

[12] S. Petti, G. Reddy, and M. M. Desai, *Inferring sparse structure in genotype–phenotype maps*, GENETICS, 225 (2023), p. iyad127, https://doi.org/10.1093/genetics/iyad127, https://academic.oup.com/genetics/article/doi/10.1093/genetics/iyad127/7223405.

[13] A. Posfai, J. Zhou, D. M. McCandlish, and J. B. Kinney, *Gauge fixing for sequence-function relationships*, PLOS Computational Biology, 21 (2025), p. e1012818, https://doi.org/10.1371/journal.pcbi.1012818, https://dx.plos.org/10.1371/journal.pcbi.1012818.

[14] S. C. Potter, A. Luciani, S. R. Eddy, Y. Park, R. Lopez, and R. D. Finn, *HMMER web server: 2018 update*, Nucleic Acids Research, 46 (2018), pp. W200–W204, https://doi.org/10.1093/nar/gky448, https://academic.oup.com/nar/article/46/W1/W200/5037715.

[15] C. E. Rasmussen and C. K. I. Williams, *Gaussian Processes for Machine Learning*, Adaptive Computation and Machine Learning, MIT Press, Cambridge, Mass., 3. print ed., 2008.

[16] G. Reddy and M. M. Desai, *Global epistasis emerges from a generic model of a complex trait*, eLife, 10 (2021), p. e64740, https://doi.org/10.7554/eLife.64740, https://elifesciences.org/articles/64740.

[17] A. F. Rubin, H. Gelman, N. Lucas, S. M. Bajjalieh, A. T. Papenfuss, T. P. Speed, and D. M. Fowler, *A statistical framework for analyzing deep mutational scanning data*, Genome biology, 18 (2017), p. 150.

[18] K. S. Sarkisyan, D. A. Bolotin, M. V. Meer, D. R. Usmanova, A. S. Mishin, G. V. Sharonov, D. N. Ivankov, N. G. Bozhanova, M. S. Baranov, O. Soylemez, N. S. Bogatyreva, P. K. Vlasov, E. S. Egorov, M. D. Logacheva, A. S. Kondrashov, D. M. Chudakov, E. V. Putintseva, I. Z. Mamedov, D. S. Tawfik, K. A. Lukyanov, and F. A. Kondrashov, *Local fitness landscape of the green fluorescent protein*, Nature, 533 (2016), pp. 397–401, https://doi.org/10.1038/nature17995, https://www.nature.com/articles/nature17995.

[19] V. W. C. Soo, J. B. Swadling, A. J. Faure, and T. Warnecke, *Fitness landscape of a dynamic RNA structure*, PLOS Genetics, 17 (2021), p. e1009353, https://doi.org/10.1371/journal.pgen.1009353, https://dx.plos.org/10.1371/journal.pgen.1009353.

[20] M. Steinegger and J. Söding, *MMseqs2 enables sensitive protein sequence searching for the analysis of massive data sets*, Nature Biotechnology, 35 (2017), pp. 1026–1028, https://doi.org/10.1038/nbt.3988.

[21] A. Tareen, M. Kooshkbaghi, A. Posfai, W. T. Ireland, D. M. McCandlish, and J. B. Kinney, *MAVE-NN: Learning genotype-phenotype maps from multiplex assays of variant effect*, Genome Biology, 23 (2022), p. 98, https://doi.org/10.1186/s13059-022-02661-7, https://genomebiology.biomedcentral.com/articles/10.1186/s13059-022-02661-7.

[22] P. D. Tonner, A. Pressman, and D. Ross, *Interpretable modeling of genotype–phenotype landscapes with state-of-the-art predictive power*, Proceedings of the National Academy of Sciences, 119 (2022), p. e2114021119, https://doi.org/10.1073/pnas.2114021119, https://pnas.org/doi/full/10.1073/pnas.2114021119.

[23] N. C. Wu, C. A. Olson, and R. Sun, *High-throughput identification of protein mutant stability computed from a double mutant fitness landscape: High-Throughput Identification of Protein Mutant Stability*, Protein Science, 25 (2016), pp. 530–539, https://doi.org/10.1002/pro.2840, https://onlinelibrary.wiley.com/doi/10.1002/pro.2840.

[24] J. Zhou, C. Martí-Gómez, S. Petti, and D. M. McCandlish, *Learning sequence-function relationships with scalable, interpretable Gaussian processes*, Nov. 2025, https://doi.org/10.7554/eLife.108964.1, https://elifesciences.org/reviewed-preprints/108964v1.

[25] J. Zhou and D. M. McCandlish, *Minimum epistasis interpolation for sequence-function relationships*, Nature Communications, 11 (2020), p. 1782, https://doi.org/10.1038/s41467-020-15512-5, https://www.nature.com/articles/s41467-020-15512-5.

[26] J. Zhou, M. S. Wong, W.-C. Chen, A. R. Krainer, J. B. Kinney, and D. M. McCandlish, *Higher-order epistasis and phenotypic prediction*, Proceedings of the National Academy of Sciences, 119 (2022), p. e2204233119, https://doi.org/10.1073/pnas.2204233119, https://pnas.org/doi/full/10.1073/pnas.2204233119.
